# Excretory-secretory products of the fish-borne parasite *Anisakis simplex* L3 larvae possess allergens and unusual glycan modifications

**DOI:** 10.1101/2025.06.26.661700

**Authors:** Isabella Adduci, Xiaoxu Wang, Katharina Paschinger, Iain B. H. Wilson, Guofeng Cheng, Shi Yan

**Affiliations:** Institute of Parasitology, Department of Biological Sciences and Pathobiology, University of Veterinary Medicine Vienna, 1210 Vienna, Austria; School of Biotechnology Jiangsu University of Science and Technology, Zhen Jiang, 212100, China; Institute of Biochemistry, University of Natural Resources and Life Sciences, Vienna, 1190 Vienna, Austria; Shanghai Tenth People’s Hospital, Institute for Infectious Diseases and Vaccine Development, Tongji University School of Medicine, Shanghai, China; Clinical Center for Brain and Spinal Cord Research, Tongji University, Shanghai 200092, China; Affiliated Shanghai Blue Cross Brain hospital, School of Medicine, Tongji University, Shanghai 200020, China

**Keywords:** Parasitic nematode, *Anisakis simplex*, allergen, glycan, glycomics

## Abstract

*Anisakis simplex* is a parasitic aquatic nematode, which may cause mild-to-severe gastrointestinal allergic reactions (Anisakiasis) with clinical symptoms, such as rhinitis and urticaria in humans who accidentally consume raw or undercooked marine products contaminated with infective L3 *Anisakis* larvae. Several *Anisakis* excretory/secretory (E/S) products and somatic proteins are known to be involved in IgE-mediated allergic reactions. In comparison to vertebrates, nematodes have a distinct machinery to glycosylate their proteins and unusual glycan structures have been reported previously, many of which play immunogenic and immunomodulatory roles in host-parasite interactions. While an early study indicated that O-glycans participate the cross-reactivity of antibodies in allergy patients to *A. simplex* somatic antigens, the N-glycosylation pattern of *Anisakis* and the potential role of N-glycans in allergic reactions remained unknown. The aim of this study was to characterise N-glycans and the associated glycoproteins from *Anisakis* E/S products using mass spectrometry. We collected E/S products from larvae culture and released N-glycans from trypsinised proteins using PNGase Ar. Native glycans were pyridylaminated prior to HPLC separation and MALDI-TOF-MS/MS analysis. In addition, hydrofluoric acid and glycosidase digestions were performed to aid structural characterisation. MS data of 5h and 24h E/S products indicated the presence of pauci-mannose and core fucosylated N-glycans as major species; tri-fucosylated and methylated glycans as well as complex-type and phosphorylcholine-substituted glycans were also detected. In addition, E/S products were subject to proteomics analysis which revealed a set of proteins with conserved domains associated with allergens. Our study provides the first insight into the N-glycosylation machinery of *Anisakis* and highlights the needs for investigating whether and which N-glycans are indubitably involved in the modulation of allergic responses.

## Introduction

*Anisakis simplex* is a parasitic nematode that naturally infects fish and marine mammals. Consumption of raw or undercooked seafood containing the third-stage larvae (L3) of *Anisakis* species may cause anisakiasis and IgE-mediated acute allergic reactions in humans. The estimated global incidence of anisakiasis is 0.32 cases per 100,000 people [1]. After L3 infection of human, symptoms such as abdominal pain, nausea, vomiting, and diarrhoea may occur [2]. *Anisakis simplex*, once ingested, penetrates the gastric and intestinal mucosa, leading to the symptoms of Anisakiasis, which are clinically manifested as rhinitis, urticaria, and, in severe cases, anaphylactic shock [3–5]. Although cooking or freezing can kill the parasites, it may not reduce their allergenicity because *A. simplex*-derived allergens have strong heat and frost resistance, and allergic reactions may still occur after consumption of contaminated seafood [6]. At present, *A. simplex* is known to have the highest number of allergens among parasites [7]. Allergens can be divided into two categories: one from the larval body wall (somatic) and the other from the excretory-secretory (E/S) antigens released by *A. simplex*, with 14 species recognised by the World Health Organization and International Union of Immunological Societies (WHO/IUIS) Allergen Nomenclature Sub-Committee to date [8–13]. Most of these allergens are involved in IgE-mediated allergic reactions to *A. simplex*. In recent years, there has been a significant progress for understanding the immunobiology of *A. simplex*. For example, a series of molecules related to the parasite development and allergy-causing to host have been identified through genomic [14], transcriptomic [15], and proteomic analyses [16], and some of their functions have been studied *in vitro* and *in vivo* [17].

Glycan modifications in allergens represent key structural features that can significantly influence immune recognition and regulation. In particular, N-glycans produced by plants, insects, and helminths often contain atypical epitopes, such as core α1,3-fucose, that are absent in mammalian glycoproteins. These non-mammalian glycoepitopes are capable of binding specifically to IgE antibodies present in the sera of allergic individuals, thereby contributing to allergenicity. Nematodes, in contrast to vertebrates, exhibit unique mechanisms of protein glycosylation, which are implicated in both their development and immune evasion strategies. While current research on N- and O-glycans in parasitic nematodes has primarily focused on identifying immunogenic glycoconjugates [18], the free-living nematode *Caenorhabditis elegans* has been used as an excellent model for dissecting the biosynthetic pathways and functional roles of nematode-specific glycosylation [19]. Studies have shown that *C. elegans* expresses a set of functional polypeptide *N-*acetyl-galactosaminyltransferases (ppGaNTases), which are involved in the initiation of mucin-type O-glycans [20] that play an important role in the embryonic development of the worm. In the parasitic nematode *Anisakis simplex*, O-glycosylation of the antigens UA2R and UA3R (Ani s 7) has been reported to influence diagnostic specificity, indicating that O-glycans may increase the capability of recognition by host immune system [21]. Another study suggested *O*-glycosylated forms of Ani s 7 and Ani s 1 may alter allergic reactions in human hosts [17]. Despite these findings, little is known about the N-glycosylation potential of *A. simplex* or the role of N-glycans in its allergenicity. In contrast, the N-glycomes of a few related nematodes within the same clade (clade III), including the closely related species *Ascaris suum*, have already been characterised [22].

In this study, we sought to analyse N-glycan modifications of E/S products derived from *A. simplex* L3 larvae and identified associated glycoproteins. An analytical workflow optimised for studying nematode N-glycans was developed, which revealed a series of N-glycan structures. Therefore, we used mass spectrometry to determine the characteristics of N-glycans in *Anisakis simplex* E/S products. Additionally, chemical and glycosidase digestions were performed to determine structural features. Concurrently, proteomic analysis of the E/S products was conducted, revealing a set of proteins with conserved domains related to allergens. Our study is the first to reveal the N-glycans structures of *Anisakis simplex* and the N-glycosylation machinery behind, providing new insights into the potential role of glycan modifications in allergic reactions triggered by *Anisakis simplex*.

## Results

### Proteins and allergens detected in E/S products

To identify proteins present in the excretory-secretory (E/S) products of Anisakis simplex L3 larvae, samples were collected from *in vitro*-cultured larvae and subjected to tryptic digestion followed by deglycosylation prior to mass spectrometry analysis (**Figure 1A**). The resulting LC-MS/MS raw data were first searched against all *A. simplex* protein sequences available in the UniProtKB database. In total, 582 unique proteins were identified: 510 proteins in E/S fraction and 188 in extracellular vesicles (EVs), with 116 proteins shared among the two sample types (**Figure 1B**). Approximately one-third of the identified proteins are of unknown function and therefore could not be classified using the PANTHER system (**Figure 1C**). The remaining proteins were primarily associated with two major molecular function categories: catalytic activity (GO:0003824) and binding (GO:0005488), consistent with previous proteomic analyses [23, 24]. The most abundant proteins were: Apolipophorins (A0A0M3IXZ4 and A0A0M3IYH2), an uncharacterised protein (A0A0M3K1Y0, predicted to be a Vitellogenin domain-containing protein), the Ani s 13 allergen haemoglobin (A0A1W7HP34 and A0A1W7HP31), a globin-like protein (A0A0M3KIW7) highly homologous to Ani s 13 (201-332 aa), Glutamate dehydrogenase (A0A0M3K4H2), and the major allergen Ani s 1 (L7V0K9) (**Figure 1D-F**). Additionally, ten of the fourteen WHO/IUIS-registered *Anisakis* allergens were identified, corresponding to a total of sixteen protein sequences (**Table 1**). Notably, six out of these allergens are predicted to carry N-glycosylation sites, and three (Ani s 3, Ani s 6 and Ani s 13) were detected in our samples (**Supplementary Table 1**). These findings prompted further investigation into the N-glycan structures associated with *A. simplex* E/S glycoproteins.

**Figure 1.**
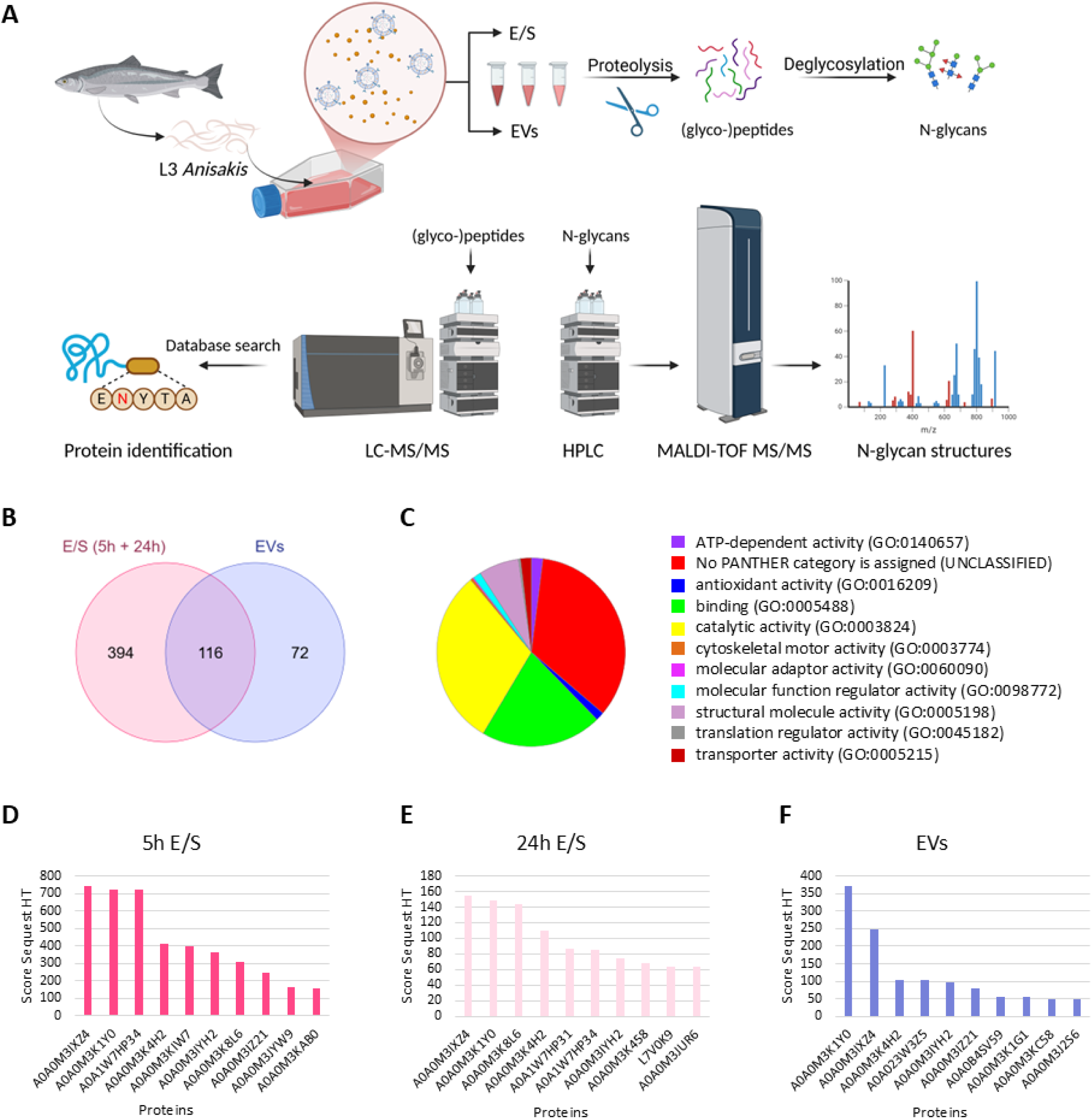
Analytical workflow and proteomic summary of Anisakis simplex excretory/secretory products (E/S) and extracellular vesicles (EVs) identified by LC-MS/MS. (**A**) Schematic overview of the sample preparation and mass spectrometry-based proteomic and glycomic analyses (created with BioRender.com). (**B**) Venn diagram showing the numbers of identified proteins unique to or shared by different samples. (**C**) Classification of molecular functions for all 582 identified proteins by PANTHER [48]. (**D-F**) Top ten proteins of each sample, ranked by Sequest HT Scores (see **Supplementary Table 1** for full dataset).

**Table 1.**
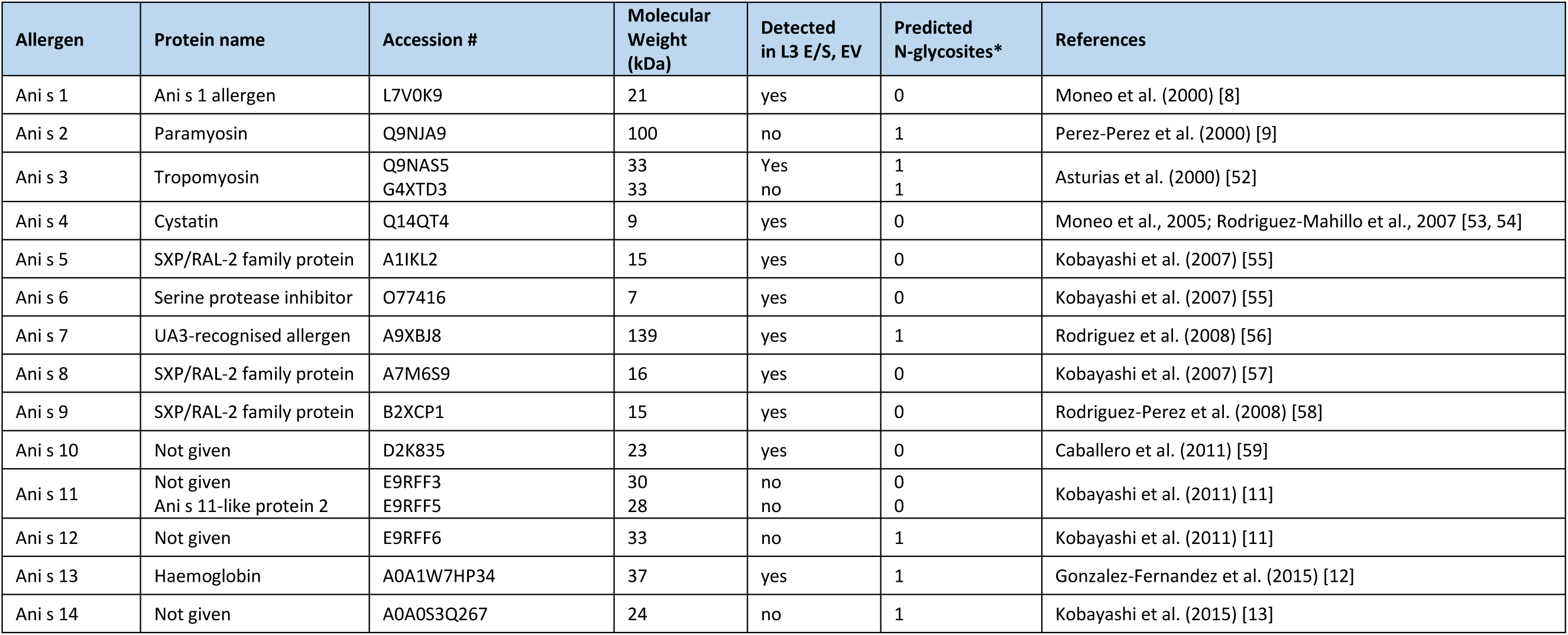
WHO/IUIS registered Anisakis allergenic proteins and their presence in L3 E/S products. Allergens listed here are accepted in the International Union of Immunological Societies (IUIS) Database [50]. * N-glycosylation sites are predicted using the NetNGlyc-1.0 server [51].

### Major N-glycans detected in E/S products

Many parasitic nematodes carry core α1,3-linked fucose on N-glycans, which is a classic cross-reactive carbohydrate determinant (CCD) shared with pollen and insect-derived allergens [25, 26]. In *C. elegans*, this epitope can be further modified with α-linked galactose, and N-glycans carrying such a disaccharide unit can only be released by hydrazinolysis or by the rice PNGase Ar [27]. To maximise glycan release efficiency, we optimised the workflow and used PNGase Ar to release N-glycans from C18-purified tryptic peptides of the E/S products, collected after 5-hour (5h E/S) and 24-hour incubation (24h E/S). 2-aminopyridine (PA) labelling of native glycans was also conducted at a smaller scale. MS spectra of PA-glycans showed an obvious difference between the two samples (**Figure 2**). In 5h E/S, one predominant peak at *m/z* 973.4 and four major peaks at *m/z* 811.4, 1119.4, 1133.5 and 1135.4 were observed, representing glycan compositions of mono- and di-fucosylated structures (Hex_1-3_HexNAc_2_Fuc_1-2_Me_0-1_); the less abundant masses detected at *m/z* 665.3, 827.4, 1151.4 and 1313.5 indicated the presence of paucimannosidic glycans (Hex_1-5_HexNAc_2_), whereas a minor portion of oligomannosidic structures (Man_7-9_GlcNAc_2_) were also observed. In contrast, at least six major glycan masses were observed on the MS spectrum of 24h E/S. The predominant structures in 24h E/S are paucimannosidic glycans as judged by masses of *m/z* 665.3, 827.4 and 1151.4 (Hex_1_HexNAc_2_, Hex_2_HexNAc_2_ and Hex_4_HexNAc_2_). Various mono- and di-fucosylated structures were observed at *m/z* 973.4, 987.4, 1119.4, 1133.5 and 1135.4 (Hex_2-3_HexNAc_2_Fuc_1-2_Me_0-1_) and one mass representing a Man_7_GlcNAc_2_ structure was detected at *m/z* 1637.6. A glycan observed at *m/z* 1427.5 in both samples indicated the presence of a tri-fucosylated glycan (Hex_3_HexNAc_2_Fuc_3_).

**Figure 2.**
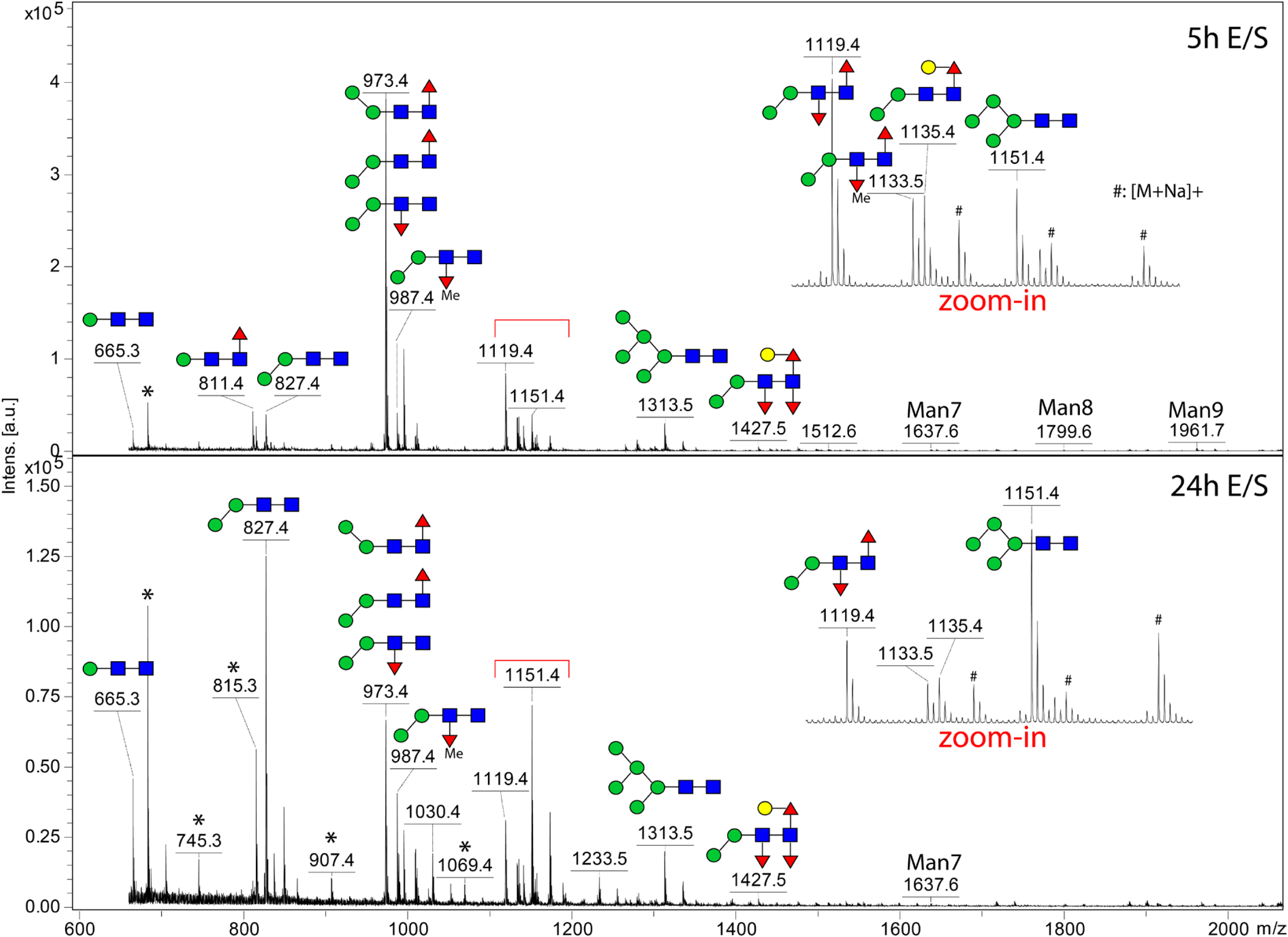
Comparison of PA-labelled N-glycans released from Anisakis E/S products. Prior to HPLC separation, *Anisakis* N-glycans were measured using MALDI-TOF MS. A comparison of the 5-hour and 24-hour samples indicated distinct MS profiles. Spectrum of the 5h E/S is dominated by mono- and di-fucosylated glycans (m/z 973.4 and 1119.4), whereas in 24h E/S paucimannosidic glycans are abundant (m/z 665, 827 and 1151). Expect for tri-fucosylated species, most of the glycans don’t carry the α1,3-fucose on the Asn-linked GlcNAc. Asterisks indicate impurities, which are hexose isomers; hashtags indicate sodiated adducts of glycans ([M+Na]^+^).

To gain more structural insights, the two PA-labelled glycan pools were fractionated on HPLC prior to MALDI-TOF MS/MS analysis. Separation of 5h E/S glycans (**Figure 3A**) was carried out using a RP-amide column with the same setting employed in previous studies [28, 29]. Therefore, many glycan structures could be proposed using their MS/MS fragmentation patterns and glucose units (g.u.) aligned with a PA-labelled dextran standards. Separation of 24h E/S glycans (**Figure 3B**) was done using the same gradient but the flow rate was set at 0.4 ml/min, which resulted in a delayed elution of glycans in comparison to the separation of 5h E/S (0.8 ml/min). While a small portion of oligomannosidic glycans (Man_6-9_GlcNAc_2_) was resolved in early eluted fractions (g.u. 5.0 – 6.5), major glycans were eluted in fractions between g.u. 6.0 and g.u. 10. Chromatograms of 5h and 24h E/S displayed difference elution patterns. The major peak of 5h E/S eluted at 8.7 - 8.9 g.u. contained three major fucosylated structures, whereas two major peaks of 24h E/S (g.u. 6.0 and 7.3) contained primarily pauci-mannosidic structures (Man_1-5_GlcNAc_2_). Phosphorylcholine (PC)-substituted glycans were primarily detected in 5h E/S (*m/z* values of 1503 and 1560), indicating the presence of two classic structures (Man_3_GlcNAc_3_Fuc_1_PC_1_ and Man_3_GlcNAc_4_PC_1_) carrying antennal PC-modifications (**Supplementary Table 2** and their MS/MS spectra provided in **Supplementary Information**). Moreover, all HPLC fractions were analysed by MALDI-TOF MS in negative ion mode, which did not reveal any N-glycans bearing negatively charged modifications, such as glucuronic acid or sialic acid.

**Figure 3.**
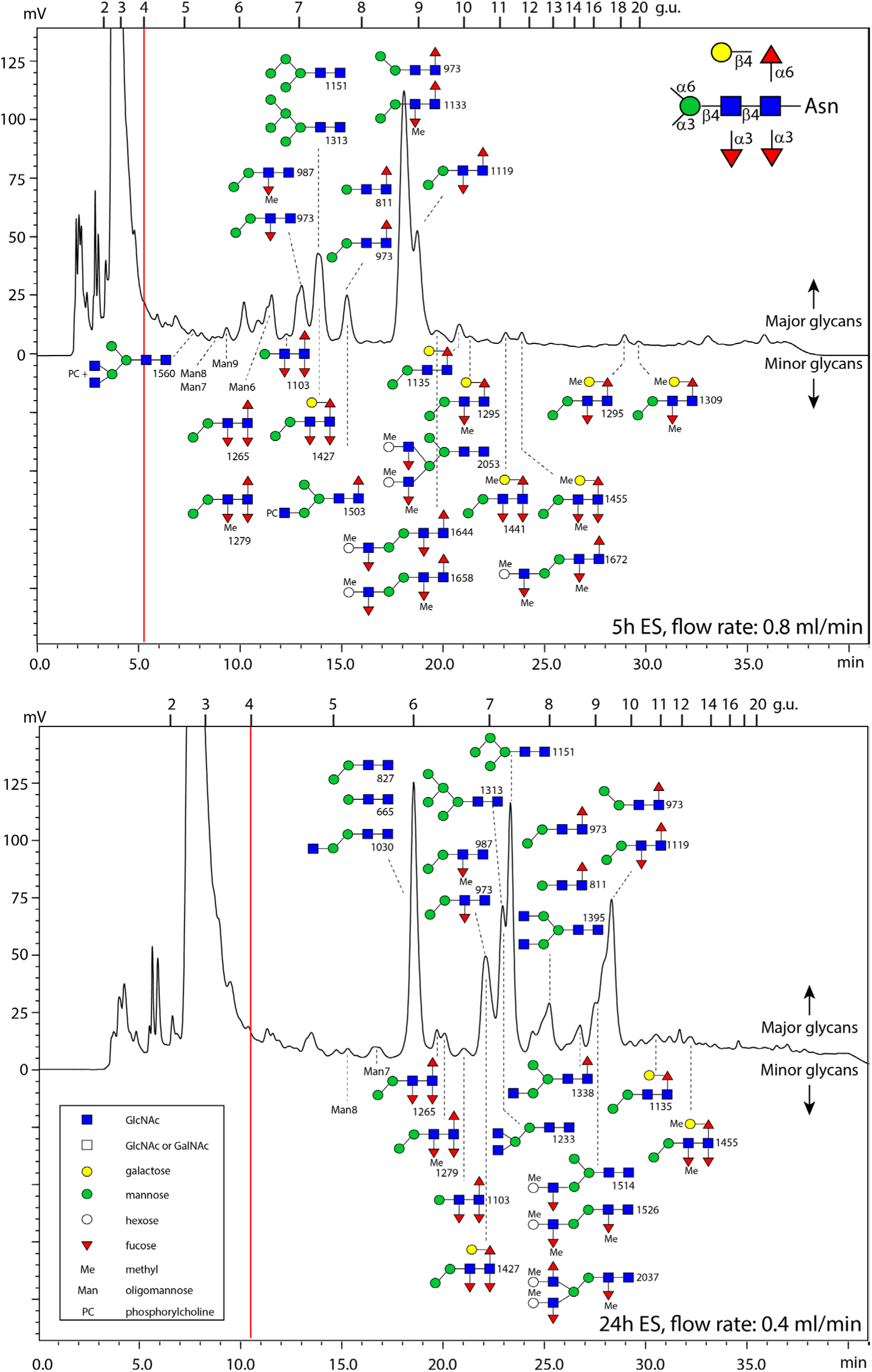
HPLC chromatograms of Anisakis N-glycans. PA-labelled N-glycans were separated by HPLC using a Ascentis^®^ RP-Amide column and a PA-labelled dextran standard was used to calibrate the column. Glycans were manually collected and subject to MALDI-TOF MS/MS analysis. Glycan structures annotated on the chromatograms were proposed based on their glucose units (g.u.), MS/MS fragmentation patterns and sensitivities to chemical and/or glycosidase digestions.

### Fucosylated N-glycans

Core and antennal fucosylated N-glycans are commonly found in nematodes and other invertebrates [30]. On the MALDI-TOF MS spectra (**Figure 2**), structures carrying up to three fucoses were identified. After RP-HPLC separation, the glycans were subjected to MALDI-TOF MS/MS experiments, which revealed additional structural details (key fragments are provided in **Supplementary Table 2**). The predominant mono-fucosylated structure Hex_2_HexNAc_2_Fuc_1_ (*m/z* 973) in 5h E/S was composed of three isomeric glycans, separated at g.u. 7.0, 7.6 and 8.7 (**Figure 4A, C and D**). The most abundant form (g.u. 8.7) carries a core α1,6-fucose and the α1,6-mannose ‘upper arm’; the other two forms eluted earlier share the same di-mannosyl core with the α1,3-mannose ‘lower arm’ and carry either a α1,3-fucose on the distal GlcNAc residue or a core α1,6-fucose on the proximal GlcNAc. Di-fucosylated glycans were primarily two structures with compositions of Hex_2_HexNAc_2_Fuc_2_ (*m/z* 1119, g.u. 8.9) and Hex_2_HexNAc_2_Fuc_2_Me_1_ (*m/z* 1133, g.u. 8.7), which carry (methyl-)fucose on both positions (**Figure 4E-F**). The same structures were also detected in three fractions in 24h E/S at later retention times due to the lower flow rate. Interestingly, a methylated fucose was found on the distal GlcNAc residue of two major glycans in 5h E/S (Hex_2_HexNAc_2_Fuc_1-2_Me_1_, **Figure 4 B and F**), which tended to co-elute with their non-methylated versions. The small daughter ions appeared on MS/MS spectra, including m/z 446/460 and m/z 592/606 as fragmentation associated low-level ‘re-arrangement’ signals, indicated the presence of distal-GlcNAc modifying (methyl-)fucose (**Figure 4 A-B and E-F**).

**Figure 4.**
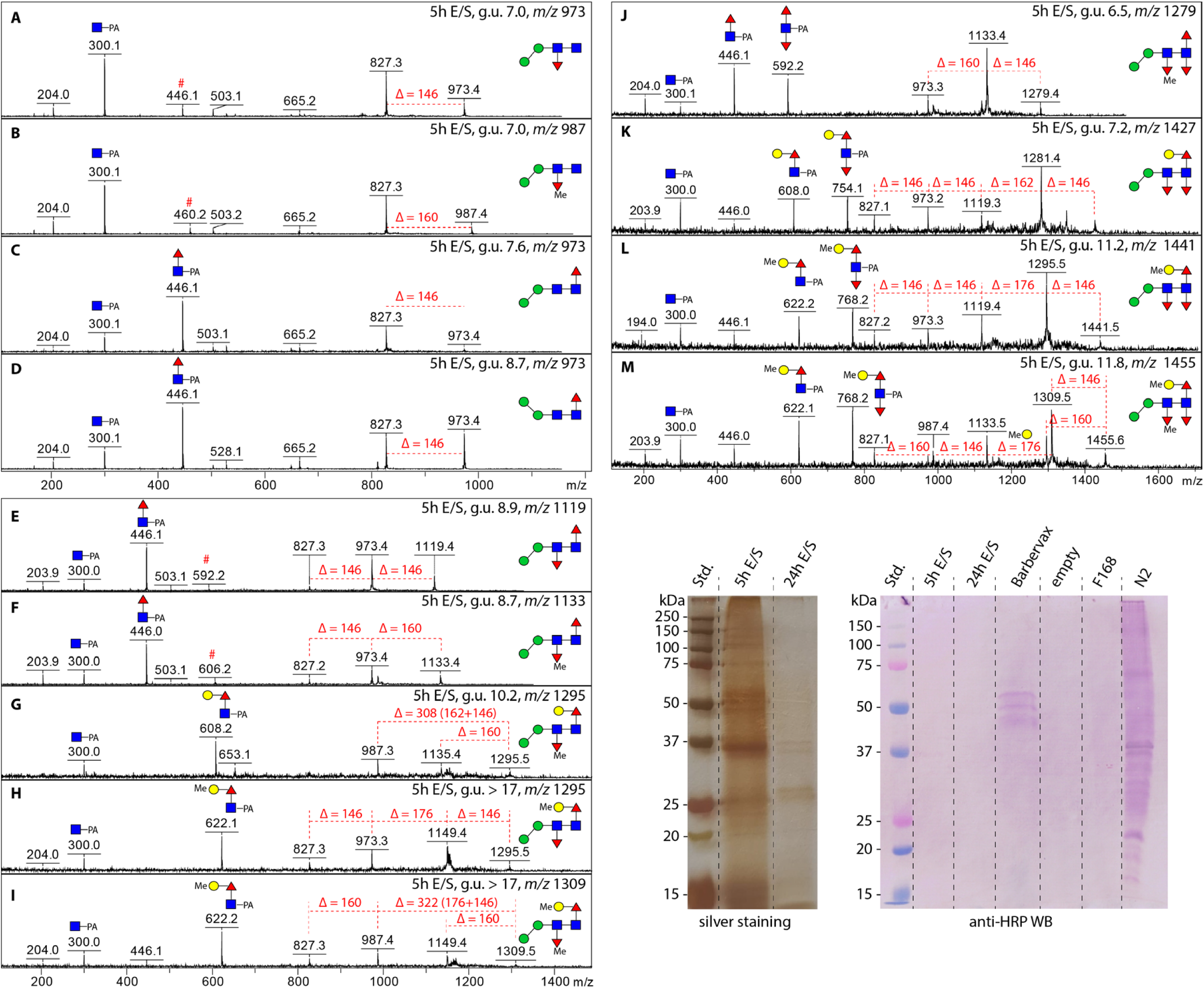
MALDI-TOF MS/MS spectra of fucosylated N-glycans and absence of anti-HRP epitope. RP-amide HPLC separated glycans carrying one (**A-D**), two (**E-I**) and three (**J-M**) fucose(s) were subject to MALDI-TOF MS/MS. Structural assessment of such glycans was based on their elution patterns (g.u.), diagnostic fragments, and in part sensitivities to hydrofluoric acid (HF) treatment (α1,3-fucose specific, MS/MS spectra of HF products are provided in **Supplementary Information**). Many structures were identified in both 5-hour and 24-hour E/S samples. Peaks marked with a red hashtag are fragment ions due to known in-source rearrangement [49]. Graphs of silver staining and Western blotting *A. simplex* proteins are provided, demonstrated no binding to the classic core α1,3-fucose CCD, also known as the anti-HRP epitope. Barbervax: glycoproteins purified from *Haemonchus contortus*; F168, a triple FUT knockout of *Caenorhabditis elegans*; N2, wild type *C. elegans*.

A series of tri-fucosylated structures (Hex_1-3_HexNAc_2_Fuc_3_Me_0-2_) were observed as a minor portion in both samples. A relative abundant species is the Hex_3_HexNAc_2_Fuc_3_ structure co-eluted with Man_4_GlcNAc_2_ and Man_5_GlcNAc_2_ at g.u. 7.2 (**Figure 4 K, Figure 3**), which displayed diagnostic ions at m/z 608 and 754 on the MS/MS spectrum, strongly indicated the presence of a Galβ1,4Fuc epitope and a core α1,3-fucose. HF treatment of this structure led to losses of two fucose residues and yielded a product of m/z 1135 (**Supplementary Information**). This structure accounts for the *m/z* 1427.5 peaks observed on MS spectra in **Figure 2**. Methylated versions of this structure as well as the de-galactosylated version were observed at *m/z* 1441, 1455 and 1279 as judged by the fragmentation patterns, the β1,4-galactose as well as the core α1,3-fucose could carry one methyl group (**Figure 4 J, L and M**). Additionally, Western blotting was carried out to detect the classic CCD core α1,3-fucose using the epitope-specific anti-HRP antibody, which indicated that there is no binding to the *A. simplex* E/S products (**Figure 4**).

### Novel glycan motifs

A subset of glycans bearing novel motifs at their non-reducing termini was detected in both E/S samples. These glycans consistently exhibited an uncommon fragment ion at *m/z* 380 in their MS/MS spectra (**Supplementary Table 2**) and were characterised by the presence of additional fucose or methylated fucose residues. The *m/z* 380 fragment is interpreted as a B-ion, consisting of a methyl-hexose (HexMe) linked to an *N*-acetyl-hexosamine (HexNAc), as indicated by the observed loss of 176 Da (corresponding to HexMe) and the presence of the characteristic *m/z* 204 fragment (**Figure 5**). The observation of m/z 526 and 540 fragments in some glycans indicated that the HexMe-HexNAc disaccharide motif is associated with either a fucose or a methyl-fucose (**Figure 5 C, G and I**). The loss of a (methyl-)fucose post HF treatment indicated α1,3-linkage (**Figure 5 D, H, J and L**). Presumably, these motifs are methylated versions of LacNAc (Galβ1-4GlcNAc) and Lewis x (Galβ1-4(Fucα1-3)GlcNAc) epitopes commonly found in humans. Although they are considered minor glycans in the whole E/S *N*-glycome, approximately 15 glycan compositions carrying such novel motifs could be identified in HPLC fractions (**Supplementary Table 2**).

**Figure 5.**
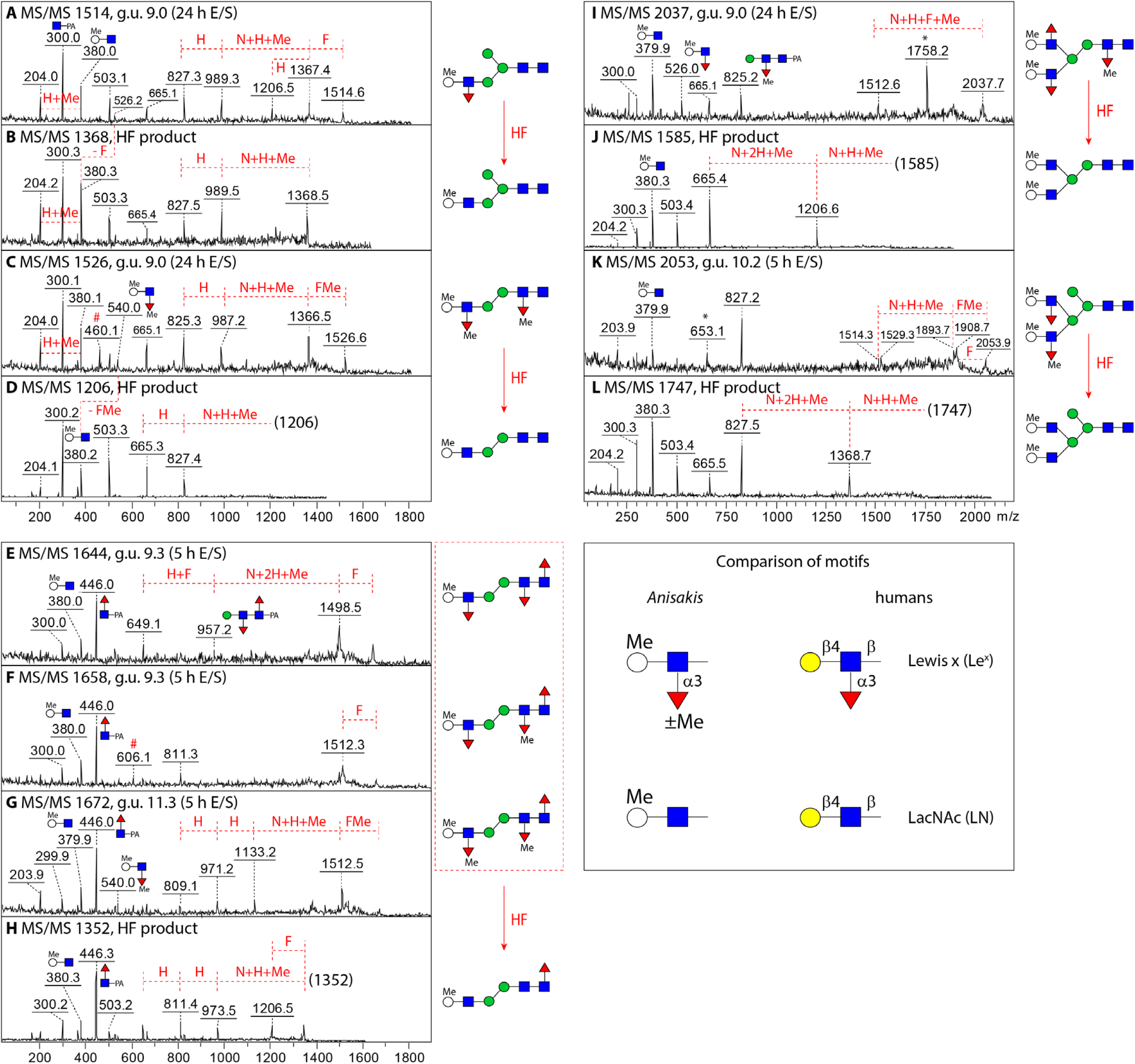
MS/MS spectra of glycans carrying novel motifs. Unusual glycans, carrying presumably a methylated Lewis-type motif, were also identified after HPLC separation. The diagnostic B-ion at *m/z* 380 was a unique signature of such structures (**A, C, E-G, I** and **K**). Loss of up to two (methyl-) fucose residues was observed post HF treatment (**B, D, H, J** and **L**); other diagnostic ions observed at *m/z* 526 and 540 (**A, C, G,** and **I,** Hex1HexNAc1Fuc1Me1-2) were absent from the MS/MS spectra of HF products. Asterisks indicate fragment ions originating from impurities.

### Major Glycoproteins in E/S products

Using FragPipe with glycoproteomics-optimised settings, a total of 156 glycopeptide-spectrum matches (glyco-PSMs) were identified, corresponding to approximately 30 distinct N-glycoproteins (**Supplementary Table 3**). Of those, 106 glyco-PSMs were detected in the 5-hour E/S samples, 44 in the 24-hour E/S samples, and 6 in the EV fraction. The most abundant glyco-PSMs were derived from haemoglobin homologs (A0A1W7HP34, A0A1W7HP35, A0A0M3KIW7), which are annotated as Ani s 13 allergens by the WHO/IUIS (**Figure 6A**). Notably, 20 glyco-PSMs originated from an SCP domain-containing protein (A0A0M3K1U4), while 35 glyco-PSMs were associated with three uncharacterised proteins (A0A0M3JWS2, A0A158PPR3, and A0A0M3KJU5), all predicted to possess multiple N-glycosylation sites. Among the 31 glyco-PSMs assigned to haemoglobin, paucimannose-type glycans with core fucosylation were predominant (**Figure 6B**). The most frequent glycan compositions included Hex₂HexNAc₂Fuc₁ (42%), Hex₁HexNAc₂Fuc₁ (29%), and Hex₃HexNAc₂Fuc₁ (13%). Analysis of LC-MS/MS spectra of a haemoglobin-derived glycopeptide (GRENYTAEDVQKDEFFVK) revealed a series of diagnostic fragment ions, strongly supporting the presence of mono-fucosylated N-glycans, Gal-Fuc epitopes and phosphorylcholine (PC) modifications on the allergen (**Figure 6 C–E**). Moreover, glycopeptides bearing di- and tri-fucosylated N-glycans, some with up to three methyl groups, were also identified. These accounted for 29% of all glyco-PSMs, as exemplified by the MS/MS spectrum of the peptide VGCGIQNCTR from the SCP domain-containing protein A0A0M3K1U4 (**Supplementary Table 3** and **Figure 6F**).

**Figure 6.**
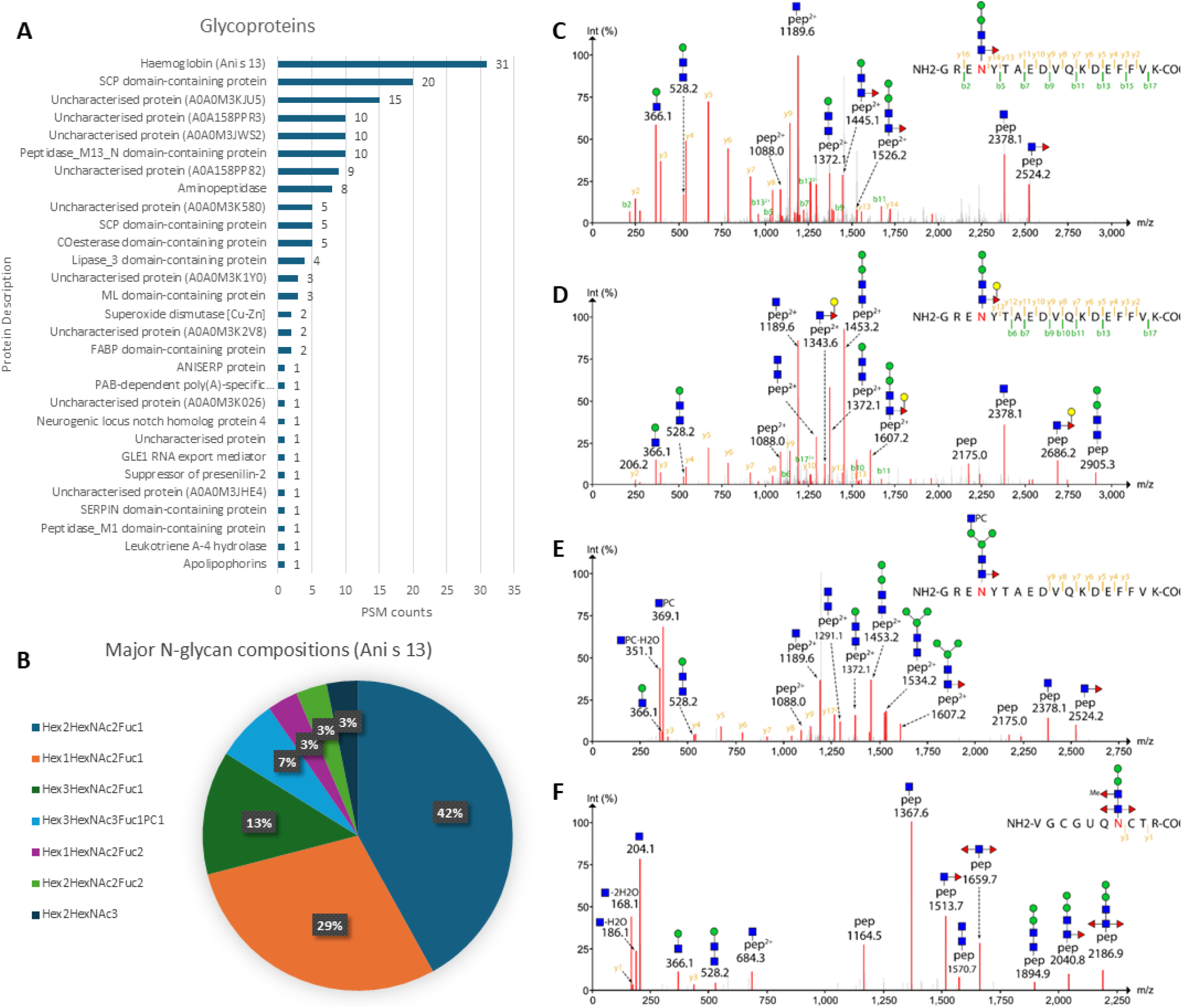
Glycoproteins, major glycan compositions and LC-MS/MS spectra of glycopeptides. (**A**) glycoproteins identified by FragPipe and sorted by PSM counts. (**B**) A pie chart of N-glycan compositions observed on haemoglobin (Ani s 13). (**C-E**) Selected MS/MS spectra of glycopeptides of haemoglobin carrying the major glycan (Man2GlcNAc2Fuc1), a glycan with Galβ1,4Fuc epitope and a glycan with phosphorylcholine (PC). (**F**) MS/MS spectrum of a glycopeptide with tri-fucosylated N-glycan. This peptide is associated with the SCP domain-containing protein (A0A0M3K1U4).

### Enzymatic basis of Anisakis N-glycan biosynthesis

Both glycomic and glycoproteome data suggested that *A. simplex* possesses a set of Golgi glycosyltransferases (GTs) and Glycoside hydrolases (GHs) that participate in core and antennal modification of N-glycans. Using sequences of characterised glycoenzymes of *C. elegans* and *homo sapiens*, BLASTp searches were performed against *A. simplex* proteins, which resulted in a range of orthologues, nine of which carrying a N-terminal transmembrane domain (detailed in **Supplementary Information**). These include four core-modifying GTs (**Figure 7**): VDK45403.1 (441 AA) and VDK47458.1 (383 AA) as the core α1,3-fucosyltransferases FUT-1 and FUT-6, a merged sequence of VDK56627.1 (289 AA) and VDK58441.1 (239 AA) as the core α1,6-fucosyltransferase FUT-8 (528 AA), VDK50184.1 (411 AA) as GT92 β1,4-galactosyltransferase GALT-1 that synthesis the Galβ1,4Fuc epitope. Four GnT homologs were also predicted to exist, including GnT I (VDK49579.1, 478 AA), GnT II (VDK17387.1, 481 AA) and GnT IV (VDK43945.1, 526 AA) and GnT V (VDK44765.1, 1268 AA).

**Figure 7.**
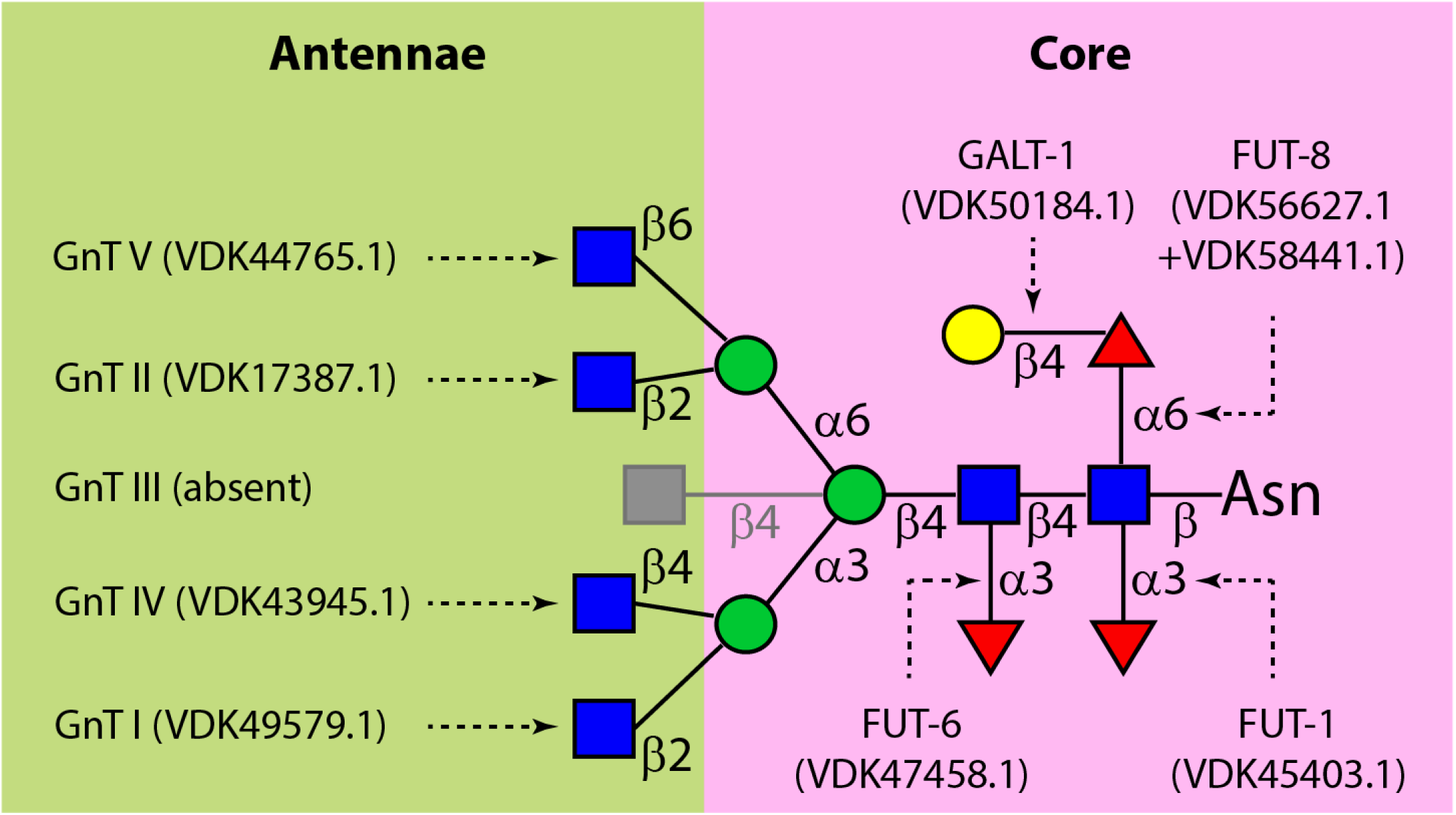
A summary of predicted glycosyltransferases involved in antennal and core modifications of *A. simplex* N-glycans. Major classes of Golgi glycosyltransferases, including fucosyltransferases (FUTs), a β1,4-galacotyltransferase (GALT-1) and N-acetyl-glucosaminyltransferases (GnT I-V) are listed. GenBank accession numbers of predicted *A. simplex* proteins are provided in the brackets.

In addition, VDK43914.1 (329 AA) was identified as a homolog of BRE-4, a *C. elegans* β1,4-N-acetylgalactosaminyltransferase involved in extending both N- and O-glycan antennae [31]. Orthologues of Golgi-associated hexosaminidases (HEX-2, 3, 4 and 5) and α-mannosidases (AMAN-2 and AMAN-3) were also identified, including VDK57457.1 (542 AA), VDK49770.1 (394 AA), VDK44985.1 (1122 AA), although these sequences were predicted to lack transmembrane domains.

## Discussion

Nematodes are capable of synthesising a wide range of glycoconjugates, and some of the glycans produced by parasitic nematodes are known to be immunogenic [22]. It is hypothesised that *A. simplex* similarly generates molecules carrying diverse N- and O-glycan structures, which may contribute to allergic reactions through cross-reactive carbohydrate determinants (CCDs) documented today. To date, at least 14 *A. simplex*-derived allergens have been identified, representing the largest number of allergens reported for nematode species. Of these, six allergenic proteins are predicted to contain N-glycosylation sites, and Ani s 7 is also thought to be O-glycosylated [21, 32].

In this study, we analysed the protein composition of excretory-secretory (E/S) products collected from *in vitro* culture of L3 larvae. The E/S products were enriched in proteins with catalytic and binding functions, highlighting their likely roles in host-parasite interactions. Notably, apolipophorins (A0A0M3IXZ4 and A0A0M3IYH2) and a vitellogenin domain-containing protein (A0A0M3K1Y0) were among the most abundant proteins. These lipid transporters belong to allergen family 92 (AF092) [33], and have homologs known to act as allergens in other species, such as the German cockroach and house dust mite [34]. Therefore, their *A. simplex* homologs were proposed as potential novel allergens in a previous study [35]. Furthermore. we identified 10 out of the 14 WHO/IUIS -recognised *Anisakis* allergens, with Ani s 13 and Ani s 1 allergens being among the most abundant. These findings provide additional evidence that L3 larvae release allergenic proteins during the early stages of host exposure, potentially contributing to immediate allergic responses in sensitised individuals. By analysing N-glycans released from E/S proteins, we characterised for the first time the N-glycosylation patterns of *A. simplex*. Our data revealed that pauci-mannose and core fucosylated glycans represent the major glycoforms of E/S products, whereas high-mannose, complex-type were also detected, albeit in lower abundance. Notably, we identified multiple N-glycan structures containing tri-fucosylated cores. Some of the tri-fucosylated glycans were reported previously in other clade V parasitic nematodes such as *Haemonchus contortus* [29], *Pristionchus pacificus* [28] and *Oesophagostomum dentatum* [36]. To our knowledge, *Anisakis* is the first nematode in clade III reported to possess such tri-fucosylated core structures.

Phylogenetically, *A.* simplex belongs to clade III and is closely related to other Ascaridida species such as *Ascaris suum, Ascaris lumbricoides, and Toxocara canis.* The N-glycosylation pattern of *A. simplex* observed in this study shared some glycomic features with *A. suum* [37, 38], including core fucosylation, the presence of Galβ1,4Fuc epitope, and PC-modifications. In addition, we detected novel motifs on N-glycan antennae, *i.e.*, a HexMe-HexNAc disaccharide with or without α1,3-linked (methyl-)fucose. A particularly striking feature of the *A. simplex* E/S N-glycome was the abundance of methylated glycans, with up to five methyl groups decorating individual glycan structures. This extensive methylation is reminiscent of findings in the free-living *C. elegans*, where L4 larvae grown in liquid culture exhibited higher levels of methylated N-glycans than those cultivated on agar plates, a phenomenon likely linked to oxidative stress [39]. In parasitic nematodes, such as *Toxocara canis* and *Heligmosomoides polygyrus*, methylated O-glycans have been shown to be highly immunogenic and capable of eliciting glycan-specific antibody responses [40, 41]. Taken together, our findings suggest that methylated N-glycans of *A. simplex*, particularly those with unusual tri-fucosylated cores or novel antennae, may play a role in modulating host immune responses and potentially contribute to allergenicity. Future studies should explore the immunological significance of these methylated glycans and assess their potential as cross-reactive carbohydrate determinants (CCDs) implicated in *Anisakis*-induced allergic reactions.

Taking advantage of glycopeptide search engine integrated in FragPipe, we identified Ani s 13 haemoglobin and its homologs (A0A1W7HP35, A0A1W7HP34, A0A1W7HP31, A0A221C790, K9USK2, A0A0M3JEL6, A0A0M3KIW7) as the most abundant glycoproteins in the E/S products. These seven proteins are highly homologous and share a conserved N-glycopeptide sequence (GRE<u>N</u>YTAEDVQKDEFFVK). The dominant glycan composition identified at this site was a core α1,6-fucosylated paucimannose structure, rather than the canonical core α1,3-fucosylated form typically associated with cross-reactive carbohydrate determinants (CCDs). Notably, several N-glycans attached to this site contained immunologically relevant motifs, including phosphorylcholine and Galβ1,4Fuc epitope. Interestingly, in our recent study, the globin-like protein A0A0M3KIW7 (also referred to as Ani941, GenBank number VDK75941.1) exhibited strong IgG reactivity in sera from *Anisakis pegreffii*-infected rats [42], suggesting its potential application as a sensitive serological marker for anisakiasis. Although it remains to be determined whether N-glycan modifications of haemoglobin and its homologs influence antibody recognition or allergenic potential, our findings provide the first direct evidence that Ani s 13 carries nematode-specific N-glycans. These results support the hypothesis that glycosylation plays a critical role in shaping the antigenic and immunomodulatory properties of *A. simplex* E/S proteins.

Protein N-glycosylation, a complex process occurring in the endoplasmic reticulum and Golgi apparatus of eukaryotes, require glycosyltransferase and glycosidase activities. In this study, we predict that *A. simplex* encodes multiple glycosyltransferases, including an orthologue of the *C. elegans* core α1,3-fucosyltransferase FUT-1, which can introduce the classic CCD core α1,3-fucose to the innermost *N-*acetyl-glucosamine residue of N*-*glycoproteins. Additionally, the presence of FUT-6 orthologue, another α1,3-fucosyltransferase known to add α1,3-linked fucose to the distal core *N-*acetyl-glucosamine in *C. elegans* and act as a Lewis-type fucosyltransferase *in vitro* [43], and an orthologue of core α1,6-fucosyltransferase (FUT-8) provide enzymatic basis for biosynthesis of tri-fucosylated structures observed in this study. Interestingly, immunoblot data did not support the presence of the classic CCD core α1,3-fucose, although this epitope was detected as part of tri-fucosylated structures by mass spectrometry. This discrepancy may be attributed to potential steric hindrance caused by additional (methyl-)fucose residues, particularly when positioned at the distal core site, which could limit the accessibility of the anti-HRP antibody.

In summary, our data reveal a diverse repertoire of N-glycan modifications in the food-borne parasite *Anisakis*, including novel antennal motifs and a subset of glycoproteins found in the excretory-secretory fraction. These nematode-specific glycans and glycoepitopes, such as phosphorylcholine (PC) and Galβ1,4Fuc, may represent previously uncharacterised classes of cross-reactive carbohydrate determinants (CCDs), in addition to the conventional CCDs implicated in allergic responses. Our findings highlight the significance of glycosylation in parasite development and host interaction, and lay the groundwork for future investigations into their immunological roles.

## Materials and Methods

### Preparation of Excretory-secretory products of Anisakis simplex

*Trichiurus haumela* fish were obtained from a local market in Shanghai were dissected and *Anisakis* L3 larvae were collected from the abdominal cavity. Worms were washed three times in 0.9% saline solution after the removal of host tissue. Afterwards, worms were transferred to T75 flasks in RPMI-1640 culture medium supplied with 1% Penicillin-Streptomycin and incubated in a CO_2_ incubator (5 %) at 37°C. After 30 minutes, culture medium was discarded to avoid remaining components from the host that are released by the worms. Subsequently, fresh RPMI-1640 medium with antibiotics was added and worms were incubated under that same condition. Half of the culture supernatant, which contains excretory-secretory (E/S) products, was collected after 5 hours (5h E/S), and the remaining half was collected after 24 hours (24h E/S). Proteins in E/S products were precipitated using pre-cooled acetone at a ratio of 1:4 (v/v). Post 30 minutes incubation at -20°C, solid precipitates were obtained by centrifugation (12,000 g, 30 min). An aliquot of protein samples was dissolved in a loading buffer and examined by silver staining post gel electrophoresis. Additionally, a small portion of extracellular vesicles (EVs) were obtained by isolating exosomes from *Anisakis* L3 larvae following a modified protocol [44]. Briefly, L3 larvae were isolated from fish as described above and cultured in PBS; culture supernatant was collected every 12 hours, and exosomes were isolated and purified using differential ultracentrifugation (100,000 × g, 4°C).

### Tryptic digestion and Proteomics analysis

80% of the collected E/S proteins were trypsinised by performing in-solution digestion procedure. Briefly, lyophilised samples were solubilised in 8 M urea in 50mM ammonium bicarbonate (pH 7.8, containing 5mM DTT) and reduced at 37°C for 1h. Subsequently, 15 mM iodoacetamide (IAA) was added to the sample followed by a short incubation in dark at room temperature (30 min). Overnight digestion with Promega Trypsin Gold was carried out at 37°C (protease: protein ratio of 1:100). Tryptic peptides were desalted using hand packed C18 cartridges (LiChroprep RP-18, 25-40 µm, Sigma-Merck) using 50% ACN with 0.1% TFA as eluent. Due to low protein concentration, the EV sample was trypsinised using a modified protocol: 33.6 µl sample was filled up to 500 µl with 8 M Urea in 50 mM Tris-buffer and was loaded on to an Pall 10kDa filter. The solution was centrifuged 2x 20 min at 10000 rcf. The proteins were first reduced with 200 mM DTT (37°C, 30 min) and then alkylated with 500 mM IAA (37°C, 30 min) on the filter. After two washes with 100 µl 50 mM Tris-buffer, digestion was carried out using Trypsin/LysC Mix (Protease:Protein ratio of 1:25) over night. Digested peptides were recovered with 3 times 50 µl of 50 mM Tris and acidified with 1µl concentrated TFA. Before LC-MS analysis peptide extracts were desalted and cleaned up using C18 spin columns (Pierce) according to the manufacturers protocol. The dried peptides were redissolved in 0.1% TFA, and 3 µl of the sample was injected to the LC-MS/MS system.

### LC-MS/MS

Peptides were separated on a nano-HPLC Ultimate 3000 RSLC system (Dionex). Sample pre-concentration and desalting was accomplished with a 5 mm Acclaim PepMap μ-Precolumn (300 µm inner diameter, 5 µm particle size, and 100 Å pore size) (Dionex). For sample loading and desalting 2% ACN in ultra-pure H2O with 0.05% TFA was used as a mobile phase with a flow rate of 5 µl/min. Separation of peptides was performed on a 25 cm Acclaim PepMap C18 column (75 µm inner diameter, 2 µm particle size, and 100 Å pore size) with a flow rate of 300 nl/min. The gradient started with 4% B (80% ACN with 0.08% formic acid) for 7min, increased to 31% in 30min and to 44% in additional 5min. It was followed by a washing step with 95% B. Mobile Phase A consisted of ultra-pure H2O with 0.1% formic acid.

For mass spectrometric analysis the LC was directly coupled to a high-resolution Q Exactive HF Orbitrap mass spectrometer (Thermo Fisher Scientific). MS full scans were performed in the ultrahigh-field Orbitrap mass analyser in ranges m/z 350−2000 with a resolution of 60 000, the maximum injection time (MIT) was 50 ms and the automatic gain control (AGC) was set to 3e^^6^. The top ten intense ions were subjected to Orbitrap for further fragmentation via high energy collision dissociation (HCD) activation over a mass range between m/z 200 and 2000 at a resolution of 15 000 with the intensity threshold at 4e^^3^. Ions with charge state +1, +7, +8 and >+8 were excluded. Normalised collision energy (NCE) was set at 28. For each scan, the AGC was set at 5e^^4^ and the MIT was 50 ms. Dynamic exclusion of precursor ion masses over a time window of 30s was used to suppress repeated peak fragmentation.

### Identification of proteins and glycoproteins

For protein identification, database searches were conducted using Proteome Discoverer (version 2.4.0.305, Thermo Fisher Scientific). The search was performed against a comprehensive protein database consisting of 25816 sequences downloaded from UniProt (https://www.uniprot.org/taxonomy/6268) and a contaminant database (crap.fasta, available at https://www.thegpm.org/crap). Trypsin was specified as the proteolytic enzyme, allowing up to two missed cleavage sites. The precursor mass tolerance was set to 10 ppm, and the fragment mass tolerance to 0.02 Da. Variable (dynamic) modifications included oxidation of methionine (+15.995 Da), deamidation of asparagine and glutamine (+0.984 Da), N-terminal acetylation (+42.011 Da), N-terminal methionine loss (−131.040 Da), and N-terminal methionine loss with acetylation (−89.030 Da). Carbamidomethylation of cysteine residues (+57.021 Da) was set as a fixed (static) modification. Peptide-spectrum matches were validated using a decoy database search with a strict false discovery rate (FDR) threshold of 1% and a relaxed FDR of 5%. Search results were exported to Microsoft Excel and comparison of proteins presented in different samples carried out using accession numbers by Multiple List Comparator (https://molbiotools.com/listcompare.php). The PANTHER server (v19.0, https://pantherdb.org/) was used to perform gene list analysis and determine GO terms. Accession numbers were used as Enter ID and *A. simplex* was selected as reference.

To identify N-glycoproteins, raw LC-MS/MS data were analysed using the open-source FragPipe software suite (v22.0), incorporating MSFragger (v4.1) and Philosopher (v5.1.1). The ‘glyco-N-HCD’ workflow was employed with default settings: one missed tryptic cleavage permitted, cysteine carbamidomethylation as a fixed modification, and methionine oxidation, as well as HexNAc (including its neutral loss), as variable modifications. Peptide- and glycan-level false discovery rates (FDRs) were set at 1%. To enhance identification of phosphorylcholine (PC)-modified glycans, additional PC-related oxonium ions (184.0728, 351.1310, 369.1416, 531.1950, 572.2210, 734.2738 m/z) were added to both the Glyco/Labile Mods and PTMs (Diagnostic Feature Extraction) tabs in MSFragger. These were supplemented to the default panel of N-glycan-specific oxonium ions (e.g., 128.0549, 138.0550, 144.0655, 163.0601, 168.0655, 186.0761, 204.0866, 243.0264, 274.0921, 290.0870, 292.1027, 308.0976, 323.2240, and 366.1395 m/z). Spectral matching was performed with a mass tolerance of ±20 ppm at both the precursor and MS/MS levels. PTM-Shepherd was configured in ‘Glyco Search’ mode using a custom glycan database containing 58 entries corresponding to N-glycan compositions identified in this study (**Supplementary Table 2**). Additionally, masses of methyl-group (Me, 14.0157 Da) and phosphorylcholine (PC, 165.0555 Da) were included into glycan Mods in the glycan database folder to support engine-level recognition of such residues.

### Release of N-glycans and PA-labelling

Glycopeptides of E/S samples were enriched by cation exchange chromatography (Dowex 50W×8; elution with 0.5 M ammonium acetate, pH 6.0) and desalted by gel filtration (Sephadex G25, 0.5% acetic acid as solvent). N-linked glycan were released using a recombinant *Oryza sativa* PNGase A (New England Biolabs) in 50 mM ammonium acetate buffer (pH 5.0) overnight at 37°C. Native glycans were separated from residual peptides by cation exchange chromatography (Dowex 50W×8); glycans collected in the filtrate fraction (as no longer bund to the column) were further purified using hand-packed C18 cartridges and nPGC cartridges prior to analysis. Native glycans were labelled with 2-aminopyridine (PA) to introduce a fluorescent tag at the reducing ends and the excess linker was removed by gel filtration (Sephadex G15, 0.5% acetic acid as solvent) as previously described [45].

### HPLC fractionation of glycans

A previously established off-line HPLC-MALDI-TOF MS analytical pipeline was followed to conduct glycomics study with minor modifications [28]. PA-labelled glycan pools were fractionated using a Shimadzu Nexera UPLC system equipped with a RF 20AXS fluorescence detector (excitation/emission: 320 nm/400 nm). N-glycans released from 5h E/S were fractionated over a Ascentis® RP-amide column (2.7 µm, 15 cm × 4.6 mm attached to a 5 cm guard column) on HPLC. A gradient of 30% (v/v) methanol (buffer B) in 0.1 M ammonium acetate, pH 4.0 (buffer A) was applied at a flow rate of 0.8 ml/min as follows: 1% buffer B per minute over 35 minutes. Similarly, N-glycans from 24h E/S were fractionated using the same gradient but at a lower flow rate of 0.4 ml/min. For column calibration, a PA-labelled partial dextran hydrolysate (2-20 glucose units) was injected prior to loading worm glycans.

### MALDI-TOF MS and MS/MS

Lyophilised glycan samples, either in native or PA-labelled forms, were dissolved in HPLC grade water and subject to MALDI-TOF MS and MS/MS analyses on a Autoflex Speed instrument (1000 Hz Smartbeam-II laser, Bruker Daltonics) using 6-aza-2-thiothymine as a matrix (Alfa-Aesar, Thermo Scientific). Calibration of the instrument was routinely performed using the Bruker Peptide Calibration Standard II to cover the MS range between 700 and 3200 Da. The detector voltage was normally set at 1977 V for MS and 2133 V for MS/MS; typically, 3000 shots from different regions of the sample spots were summed. To obtain MS/MS spectra, parent ions were fragmented by laser-induced dissociation (LID) without applying a collision gas (precursor ion selector was generally ±0.6%). For PA-labelled glycans, [M+H]^+^ ions were favoured for LID fragmentation. MS and MS/MS data are further processed with the Flexanalysis (version 3.3.80, Bruker) using the SNAP2 algorithm and manually interpreted using an in-house glycan database of nematodes.

### Chemical and exo-glycosidase digestion of glycans

48% v/v hydrofluoric acid (HF) was used to verify structures potentially modified with α1,3-linked (methyl-)fucose. Selected glycans were dried and incubated overnight at 0°C with 3 μl HF prior to evaporation; treated samples were diluted in water and re-evaporated twice, prior to MALDI-TOF MS/MS analysis. Exo-glycosidases for determining arm-specificity and the presence of the Galβ1,4Fuc epitope included recombinant *Xanthomonas manihotis* α1,2/3-mannosidase and α1,6-mannosidase (New England Biolabs) and a recombinant *Aspergillus niger* β-galactosidase prepared in-house [46]. Reactions were carried out in 25 mM ammonium acetate (pH 5.0) overnight at 37°C.

### Western blotting

Worm-derived proteins, including the E/S products of *A. simplex* L3 larvae, native antigens of *H. contortus*, whole worm lysates of *C. elegans* (N2 and F168), were dissolved in SDS-loading dye, denatured at 95°C for 10 min and resolved on 12% SDS-PAGE gels. Precision Plus Protein™ Dual Color Standards (Bio-Rad Laboratories) was used as protein ladder. Post electrophoresis at 200 V, proteins were transferred onto nitrocellulose membranes using a TransBlot® Turbo™ Transfer System (Bio-Rad Laboratories). After blocking with 0.5% BSA, immunodetection of core α1,3 fucose was performed using the anti-HRP polyclonal antibody (1:10,000, Sigma-Aldrich), followed by incubation of alkaline phosphatase-conjugated goat anti rabbit IgG antibody (1:2,000, Vector Labs), with SIGMAFAST™ BCIP®/NBT as the substrate for colour development.

### Bioinformatic analysis

Prediction of enzymes putatively involved in the N-glycan biosynthesis of A. simplex were carried out by standard protein BLAST searches (BLASTp) against all non-redundant protein sequences of *Anisakis* (taxid:6268), using selected sequences of *C. elegans* and human glycosyltransferases and glycosidases as template. These sequences include NP_001369885.1 (FUT-1), NP_494823.2 (FUT-6), NP_504555.2 (FUT-8), NP_504545.2 (GALT-1), NP_490872.1 (BRE-4), NP_741838.1 (GLY-12), NP_509566.1 (GLY-13), NP_497719.1 (GLY-14), NP_491874.1 (GLY-2), NP_505864.1 (GLY-20), NP_001091740.1 (MGAT3), NP_001253751.1 (MGAT4B), NP_001358386.1 (MGAT-5), NP_505995.2 (AMAN-2), NP_001361919.1 (AMAN-3), NP_504489.3 (HEX-2), NP_499390.3 (HEX-3), NP_740792.3 (HEX-4), NP_001366664.1 (HEX-5). Transmembrane domains of *A. simplex* proteins were predicted using the TMHMM - 2.0 server [47].

## Data availability

MALDI-TOF MS and MS/MS spectra are available on GlycoPost (GPST000396; Preview URL: https://glycopost.glycosmos.org/preview/1371834393685563bb160ef; PIN CODE: 7927). Raw LC-MS/MS data have been deposited in the ProteomeXchange Consortium via the PRIDE partner repository (accession number PXD065269); the password to access the data is available from the corresponding author upon request.

## Acknowledgement

This work has been funded by the Programme “Top Vet Science” financed by the Vetmeduni Vienna to S.Y., the Key Program for International S&T Cooperation Projects of China (2021YFE0191600 to G.C.) and the Austrian Science Fund (FWF, P32572 to K.P.). The authors gratefully acknowledge Karin Hummel at VetCore Proteomics for providing LC-MS/MS services and the BOKU Core Facility Mass Spectrometry for the access to their MALDI-TOF MS instrument. We also thank Jorick Vanbeselaere at BOKU Vienna for his assistance with LC-MS/MS data analysis.

## Author Contributions

Conceptualisation: G.C., S.Y.

Funding acquisition: G.C., S.Y.

Investigation: I.A., X.W., S.Y.

Formal analysis: S.Y., K.P., I.B.H.W.

Methodology: I.A., S.Y.

Supervision: S.Y., G.C.

Visualisation: S.Y.

Writing – original draft: S.Y., G.C., I.A.

Writing – review & editing: all authors

