## Supplementary material for "Excretory-secretory products of the fish-borne parasite *Anisakis simplex* L3 larvae possess allergens and unusual glycan modifications": Supplymentary Information

#### Supplementary Figure

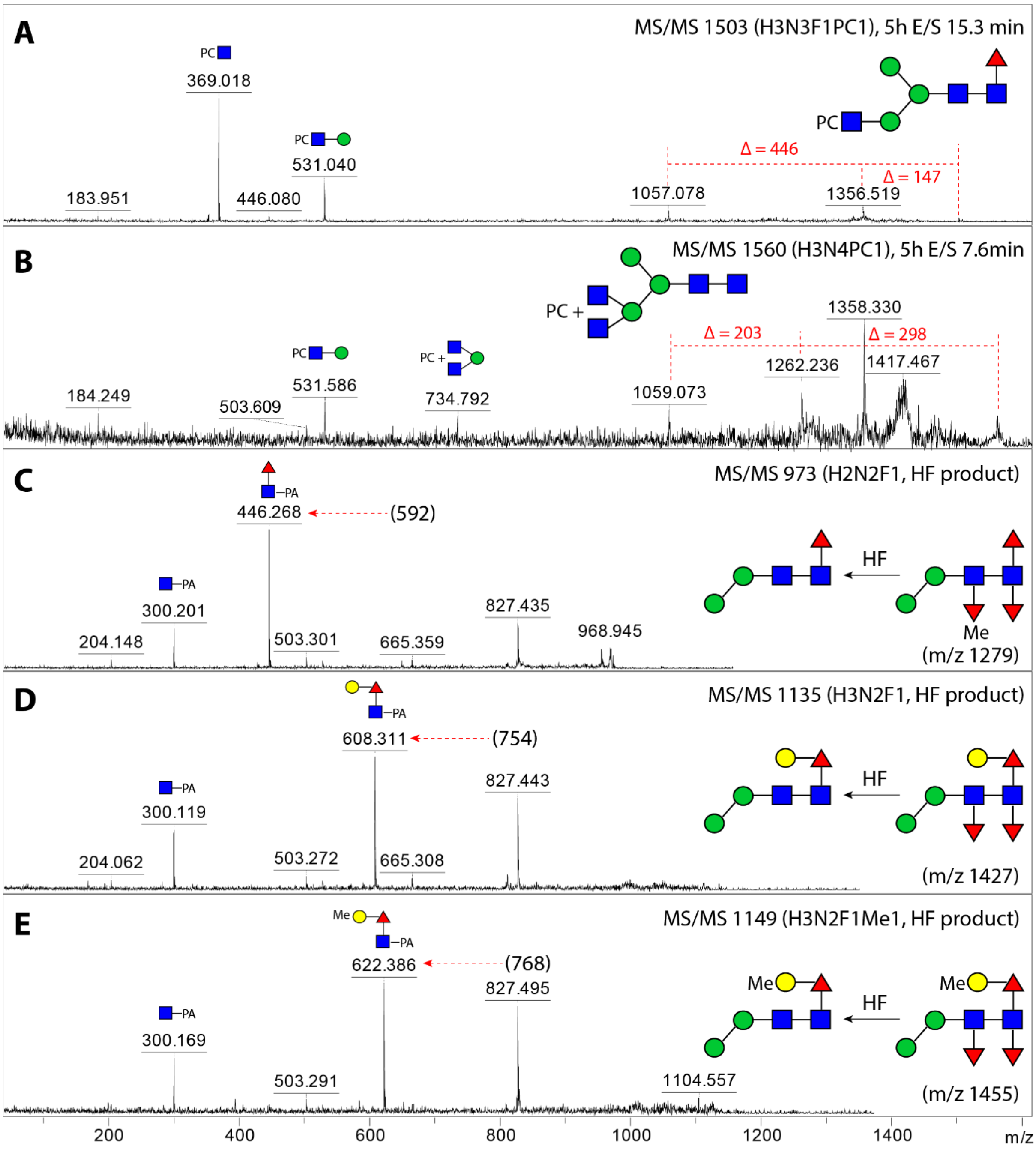


MALDI-TOF MS/MS spectra of phosphorylcholine (PC)-modified N-glycans and tri-fucosylated glycan products following hydrofluoric acid (HF) treatment. (**A–B**) Two PC-modified N-glycans identified in the 5-hour excretory-secretory (E/S) products of *Anisakis simplex*. (**C–E**) HF treatment resulted in the removal of core α1,3-linked (methyl-)fucose residues, leading to the disappearance of diagnostic fragment ions at m/z 592, 754, and 768, as shown in **Figure 4J**, **4K**, and **4M**.

#### Prediction of glycoenzymes

Summary of homologous glycoenzymes in *A. simplex*:

| Enzymes | Accession # | Length (aa) | TMD (Yes/No) | Note |
|---|---|---|---|---|
| **Glycosyltransferase (GT)** | | | | |
| FUT-1 | VDK45403.1 | 441 | Yes | Identities 42 % |
| FUT-6 | VDK47458.1 | 383 | Yes | Identities 39 % |
| FUT-8 | VDK56627.1 (1-289)<br>VDK58441.1 (290-528) | 528 | Yes | Identities 58 % |
| GALT-1 | VDK50184.1 | 411 | Yes | Identities 28 % |
| GLY-12 (GnT I) | VDK49579.1<br>VDK67433.1<br>VDK30835.1 | 478<br>263<br>57 | Yes<br>No<br>No | Identities 42 % |
| GLY-13 (GnT I) | VDK67433.1<br>VDK49579.1<br>VDK30835.1 | 263<br>478<br>57 | No<br>Yes<br>No | Identities 35 % |
| GLY-14 (GnT I) | VDK67433.1<br>VDK49579.1<br>VDK30835.1 | 263<br>478<br>57 | No<br>Yes<br>No | Identities 36 % |
| GLY-2 (GnT V) | VDK44765.1 | 1268 | Yes | Identities 42 % |
| GLY-20 (GnT II) | VDK17387.1 | 481 | Yes | Identities 50 % |
| MGAT3 (GnT III) | | | | No hits |
| MGAT4B (GnT IV) | VDK42989.1<br>VDK43945.1<br>AZA15233.1 | 468<br>526<br>252 | No<br>Yes<br>No | Identities 34 % |
| MGAT5 (GnT V) | VDK44765.1 | 1268 | Yes | Identities 43 %<br>GLY-2 homolog |
| BRE-4 | VDK43914.1 | 329 | Yes | Identities 65 % |
| **Glycoside hydrolase (GH)** | | | | |
| HEX-2 | VDK57457.1 | 542 | No | Identities 39 % |
| HEX-3 | VDK57457.1 | 542 | No | Identities 34 % |
| HEX-4 | VDK49770.1 | 394 | No | Identities 45 % |
| HEX-5 | VDK57457.1 | 542 | No | Identities 32 % |
| AMAN-2 | VDK44985.1 | 1122 | No | Identities 43 % |
| AMAN-3 | VDK44985.1 | 1122 | No | Identities 31 % |

#### FUT-1

Alpha-(1,3)-fucosyltransferase

*C. elegans*: 433 aa NP_001369885.1

>NP_001369885.1 Alpha-(1,3)-fucosyltransferase fut-1 [Caenorhabditis elegans]
MTARSIKLFFARWKYLMFACCITYLLVIYAPISKSEQKDWKEGEIELSNDHELDVPILQKEELKPQQRPS
FEENVPKKKTFNFNPVGKEPFDVEEVLTSSDIKLEERMTATVIPGQKRLILSWNAGHSQDNLQGCPDWNC
EFTQVRARAPDADAVLIAHMDNDFVPKPNQYVVYFSQESPANSGIQIPRPDYINMTLGFRHDTPAGSPYG
YTVKLGAKSRKTGQVVDANLVNGKAKGAAWFVSHCQTNSKREDFVKKLQKHLQIDIYGGCGPMKCARGDS
KCDTMLDTDYHFYVTFENSICEDYVTEKLWKSGYQNTIIPLVLKRKLVEPFVPPNSFIAIDDFKSVKEMG
DYLNYLMNNKTAYMEYFEWRHDYKVVFLDGSHHDVLERPWGFCQVCRMAWTEPRQKVLIPNWDAYWRQTC
EKDGTLVDSIPLD

##### TMHMM result

```
# CAA91285.2 Length: 433
# CAA91285.2 Number of predicted TMHs:  1
# CAA91285.2 Exp number of AAs in TMHs: 19.56876
# CAA91285.2 Exp number, first 60 AAs:  19.56876
# CAA91285.2 Total prob of N-in:        0.94664
# CAA91285.2 POSSIBLE N-term signal sequence
CAA91285.2      TMHMM2.0        inside       1    12
CAA91285.2      TMHMM2.0        TMhelix     13    32
CAA91285.2      TMHMM2.0        outside     33   433
```

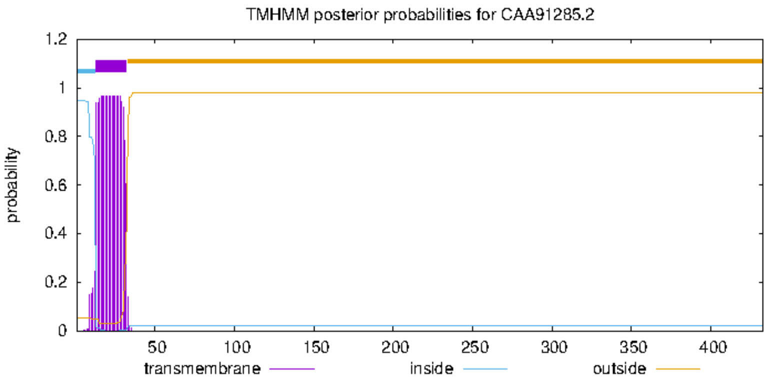


#### unnamed protein product [Anisakis simplex]

**Sequence ID: VDK45403.1Length: 441Number of Matches: 1**

>VDK45403.1 unnamed protein product [Anisakis simplex]
MFQRWCARLSWYLAHRFQYALFVIALLSTFLILPYGISTYSIFKRYARNFGTITIAPSSRRRNENAKMSN
TISLYESCGMCGSHGCTCLVSAAPKLIVSLSPDWKSNNLNGCLEWRCELSSNTNDLQRADAIIGNSALHG
FVNITKPRQYNVFYSQESPINTALPSSTKHFNLSLSYRRDSPTSSPYGYTVKLAPRSRNSRRIPDKNLLN
AKSRPLVWLLLLFKTYHTSKSALQKLTDQKDFFQQYIDVDIYGDCGQLRCTRKTSCEDVLDSKYYFYIAF
ENSICHDYVTEKLWGKGFNRLIVPIVLQRSILEAYAPPHSFIAADDFVDIKQLADHLKYLMRNSTAYREY
FDWRRDYAAIFLDGNTHDQLEQPWGICQLCRLLWQQPQKQYTIENFNEWWTKCEEPGDLVKRLIKNDSIN
DNNDHINERKLVKNKSQEQYK

### TMHMM result

```
# VDK45403.1 Length: 441
# VDK45403.1 Number of predicted TMHs:  1
# VDK45403.1 Exp number of AAs in TMHs: 22.70497
# VDK45403.1 Exp number, first 60 AAs:  22.67067
# VDK45403.1 Total prob of N-in:        0.99299
# VDK45403.1 POSSIBLE N-term signal sequence
VDK45403.1      TMHMM2.0        inside       1    20
VDK45403.1      TMHMM2.0        TMhelix     21    43
VDK45403.1      TMHMM2.0        outside     44   441
```

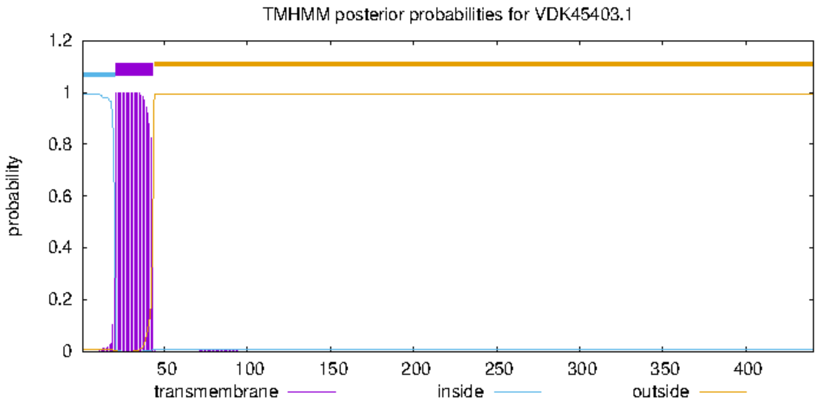


Uniprot: A0A0M3JV82_ANISI

#### PTM/Processing

##### Keywords

PTM | #Glycoprotein Automatic Annotation

#### Structure

**Model Confidence:**

- Very high (pLDDT > 90)
- Confident (90 > pLDDT > 70)
- Low (70 > pLDDT > 50)
- Very low (pLDDT < 50)

AlphaFold produces a per-residue confidence score (pLDDT) between 0 and 100. Some regions with low pLDDT may be unstructured in isolation.

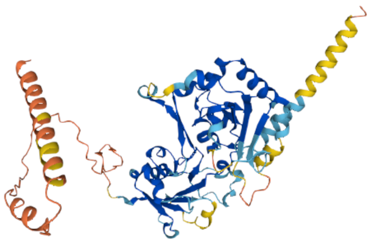

### FUT-6

Gene ID: 188084

```
>NP_494823.2 Alpha-(1,3)-fucosyltransferase fut-6 [Caenorhabditis elegans]
MSQIGGATCTWRYLGRFVTLGIYASVALFVWYTLVPTRSKHKDSIAINNNNADPATALIPVHTKNVVIYA
ATKFFGHPITTERFLATCPDVQNYCRITQEESEFDNADAVLFHNADYRGSTDKFKKMKSQRKPGVPYVLW
SLESPTNDMFRPDSHMINWTMTYRTDSDVWAPYGTIVKLKNPVEVDLNAIWEGKTKTATWLASNCITQNH
RFDLIKKIIDNGFEIDIWGNCGKQVSQCAGVDNQESPCVLELIKPYKFYISMENSNCKDYVTEKFWKALN
DRMTIPIVLARKYYKDLGVPDSAYIAVDDYATLDEFLAHVKKVNKEKDLFLSYHQWRKEWKVIIGSGFSG
WCTLCNKLQDKDYILKNPKSYKDVAWWHSFEMCNNQIASKYL
```

```
# NP_494823.2 Length: 392
# NP_494823.2 Number of predicted TMHs:  1
# NP_494823.2 Exp number of AAs in TMHs: 22.87217
# NP_494823.2 Exp number, first 60 AAs:  22.80954
# NP_494823.2 Total prob of N-in:        0.99567
# NP_494823.2 POSSIBLE N-term signal sequence
NP_494823.2     TMHMM2.0        inside       1    12
NP_494823.2     TMHMM2.0        TMhelix     13    35
NP_494823.2     TMHMM2.0        outside     36   392
```

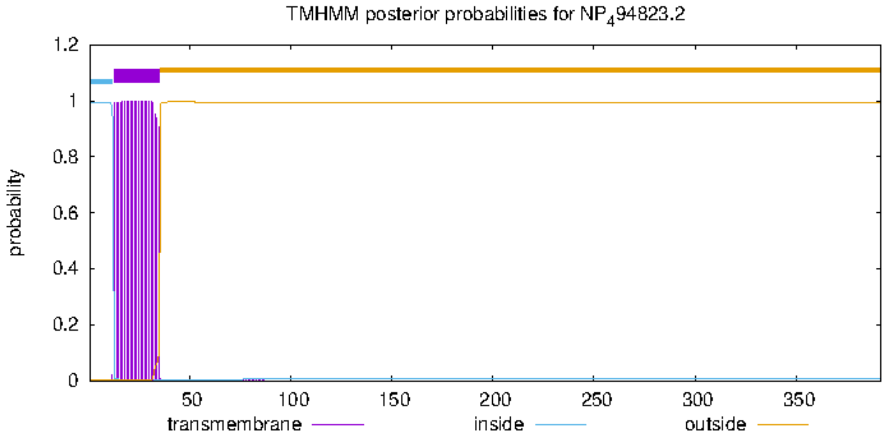


**unnamed protein product [Anisakis simplex]**

Sequence ID: VDK47458.1 Length: **383** Number of Matches: **1**

**Range 1: 4 to 360** GenPept Graphics ▼ Next Match ▲ Prev

| Score | Expect | Method | Identities | Positives | Gaps |
|---|---|---|---|---|---|
| 252 bits(644) | 4e-81 | Compositional matrix adjust. | 148/382(39%) | 211/382(55%) | 35/382(9%) |

```
>VDK47458.1 unnamed protein product [Anisakis simplex]
MAPRFKNRYLVIFIAALIGLFLLDIISSFLVGHSSSDLSSTKTTDAYRNKNVRLFAATPFFGRPIDNSWL
SKCSEYCKLVDNSESADAVMYHIPDYRFLSTNELKSNQIAVIWSLESPIYQHLSRELRGRINWTMTYRRD
SDVWFPYGVIRKRDKPIQIDYDKIWRDKKRMVVWLVSNCGHANGRLILGRALQKSGLELDIFGACGQHKT
PNDCDGVKKQSDQCVAELFMPYMFALSFENSLCKDYITEKFFEVLQKRYAIPIVMRRKDYEHIAAPPNSF
IAVDDYKNIDELISDIKTIASNKFTYLKYHRWRESYEIQSDYFHIDDTGFCALCKKLMRQRFSRKHYDDV
AAWWNSNEICEIPQDGFVNEFLSKSGMVVPQVR
```

```
# VDK47458.1 Length: 383
# VDK47458.1 Number of predicted TMHs:  1
# VDK47458.1 Exp number of AAs in TMHs: 22.5402
# VDK47458.1 Exp number, first 60 AAs:  22.43994
# VDK47458.1 Total prob of N-in:        0.97407
# VDK47458.1 POSSIBLE N-term signal sequence
VDK47458.1      TMHMM2.0        inside       1     8
VDK47458.1      TMHMM2.0        TMhelix      9    31
VDK47458.1      TMHMM2.0        outside     32   383
```

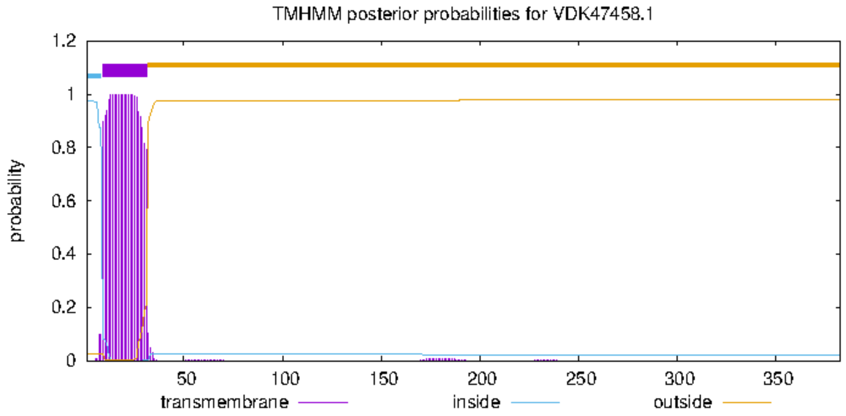


```
             1       10        20        30        40        50        60        70        80        90        100       110       120       130
             |--------+---------+---------+---------+---------+---------+---------+---------+---------+---------+---------+---------+---------|
NP_494823.2  MSQIGGATCTWRYLGRFVTLGIYASVALFVWYTLVPTRSKHKDSIAINNNNADPATALIPVHTKNVVIYAATKFFGHPITTERFLATCPDVQNYCRITQEESEFDNADAVLFHNADYRGSTDKFKKMKSQ
 VDK47458.1          MAPRFKNRYLVIFIAALIGLFLLDIISSFLVGHSSSDLSSTKTTDA------YRNKNVRLFAATPFFGRPIDNS-WLSKCSE---YCKLVDNS---ESADAVMYHIPDYRFLST------NE
  Consensus  ........capR%lnR%ltigIaAl!aLFlldiispfrsgHkdSdainnnnaDa......yrnKNVri%AATkFFGrPIdne.fLakCp#...YCrit##e...#nADAV$%HnaDYRglsd......n#

             131     140       150       160       170       180       190       200       210       220       230       240       250       260
             |--------+---------+---------+---------+---------+---------+---------+---------+---------+---------+---------+---------+---------|
NP_494823.2  RKPGVPYVLWSLESPT-NDMFRPDSHMINWTMTYRTDSDVWAPYGTIVKLKNPVEVDLNAIWEGKTKTATWLASNCITQNHRFDLIKKIIDNGFEIDIWGNCG--KQVSQCAGVDNQESPCVLELIKPYK
 VDK47458.1  LKSNQIAVIWSLESPIYQHLSRELRGRINWTMTYRRDSDVWFPYGVIRKRDKPIQIDYDKIWRDKKRMVVWLVSNCGHANGRLILGRALQKSGLELDIFGACGQHKTPNDCDGVKKQSDQCVAELFMPYM
  Consensus  rKpnqiaViWSLESPi.#d$fRedrgrINWTMTYRrDSDVWaPYGtIrKrdnP!#!Dl#aIWrdKkrmatWLaSNCghaNgRldLgraiqdnGlEiDIfGaCG..Kqpn#CaGVdnQedqCVaELikPYk

             261     270       280       290       300       310       320       330       340       350       360       370       380       390
             |--------+---------+---------+---------+---------+---------+---------+---------+---------+---------+---------+---------+---------|
NP_494823.2  FYISMENSNCKDYVTEKFWKALNDRMTIPIVLARKYYKDLGVPDSAYIAVDDYATLDEFLAHVKKVNKEKDLFLSYHQWRKEWKV---IIGSGFSGWCTLCNKLQDKDYILKNPKSYKDVA-WWHSFEMC
 VDK47458.1  FALSFENSLCKDYITEKFFEVLQKRYAIPIVMRRKDYEHIAAPPNSFIAVDDYKNIDELISDIKTIASNKFTYLKYHRWRESYEIQSDYFHIDDTGFCALCKKLMRQRF---SRKHYDDVAAWWNSNEIC
  Consensus  FaiSmENSnCKDY!TEKFfeaL#dRmaIPIV$aRKdYediaaPdna%IAVDDYaniDEliad!Kk!ak#Kdl%LkYHrWReewe!...iigiddsGfCaLCnKLqrqr%...nrKhYdDVA.WWnSnEiC

             391     400       410413
             |--------+---------+--|
NP_494823.2  N---NQIASKYL
 VDK47458.1  EIPQDGFVNEFLSKSGMVVPQVR
  Consensus  #...#qiane%L...........
```

### FUT-8

GenBank: NP_504555.2

```
>NP_504555.2 Alpha-(1,6)-fucosyltransferase [Caenorhabditis elegans]
MLKCIAAVGTVVWMTMFLFLYSQLSNNQSGGDSIRAWRQTKEAIDKLQEQNEDLKSILEKERQERNDQHK
KIMEQSHQLPPNPENPSLPKPEPVKEIISKPSILGPVQQEVQKRMLDDRIREMFYLLHSQTIENSTKILL
ETQMISLMGLSAQLEKLEGSEEERFKQRTAITQRIFKSIEKLQNPKACSEAKTLVCNLDKECGFGCQLHH
VTYCAITAFATQRMMVLKRDGSSWKYSSHGWTSVFKKLSKCSFDEAVGNTEAKPFAEPSPARVVSLGIVD
SLITKPTFLPQAVPEQLLESLTSLHSHPPAFFVGTFISYLMRFNSATQEKLDKALKSIPLDKGPIVGLQI
RRTDKVGTEAAFHALKEYMEWTEIWFKVEEKRQGKPLERRIFIASDDPTVVPEAKNDYPNYEVYGSTEIA
KTAQLNNRYTDASLMGVITDIYILSKVNYLVCTFSSQVCRMGYELRQPSGADDGSKFHSLDDIYYFGGQQ
AHEVIVIEDHIAQNNKEIDLKVGDKVGIAGNHWNGYSKGTNRQTYKEGVFPSYKVVNDWRKFKFEALLD
```

```
# CCD64126.1 Length: 559
# CCD64126.1 Number of predicted TMHs:  1
# CCD64126.1 Exp number of AAs in TMHs: 16.75743
# CCD64126.1 Exp number, first 60 AAs:  16.53576
# CCD64126.1 Total prob of N-in:        0.83363
# CCD64126.1 POSSIBLE N-term signal sequence
CCD64126.1     TMHMM2.0     inside      1     4
CCD64126.1     TMHMM2.0     TMhelix     5    24
CCD64126.1     TMHMM2.0     outside    25   559
```

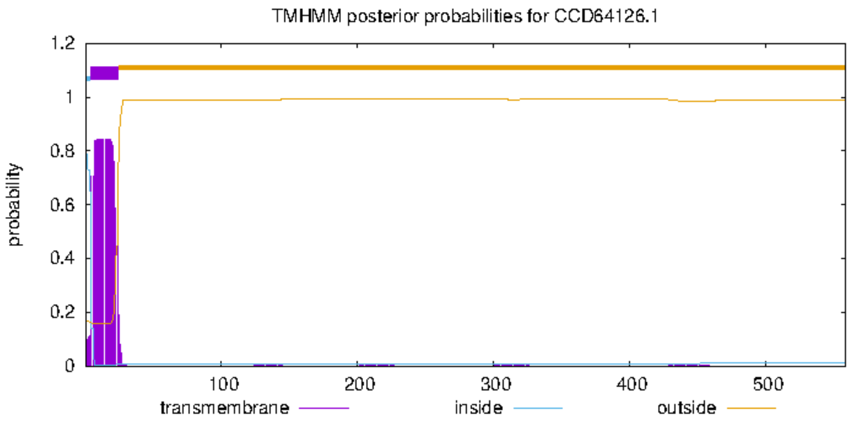


#### unnamed protein product [Anisakis simplex]

**Sequence ID:** VDK58441.1 **Length:** 239 **Number of Matches:** 1

**Range 1: 1 to 233** GenPept Graphics ▼ Next Match ▲ Previou

| Score | Expect | Method | Identities | Positives | Gaps |
|---|---|---|---|---|---|
| 304 bits(778) | 4e-101 | Compositional matrix adjust. | 135/233(58%) | 176/233(75%) | 0/233(0%) |

```
>VDK58441.1 unnamed protein product [Anisakis simplex]
MRYLMRPNSSLAKRITEFAAKVSFESGPIVGLQIRRTDKVGTEAEFHALSEYMKWTEYWFRIQEYRHGKA
VKRRIYVATDDPTVFSEARKKYPNYEVFGDAAISNTANTRSRYSIESLYGVIIDIEMLARCDYLVCTFSS
QVCRMGYELMQIRVGDAGDNFHSLDDLYYYGGQQAHEQVVVESYQAESKDEIDLEIGDTIGIAGNHWDGF
SKGTNRRNGAVGLYPSYKTREKWIIVPFP
```

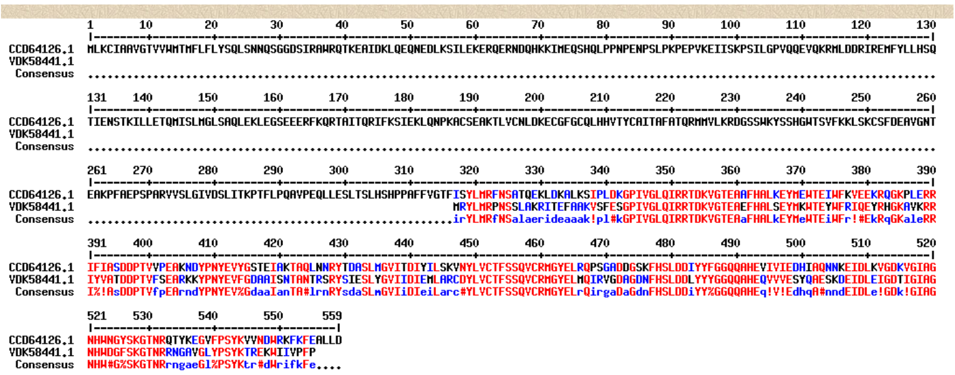

#### unnamed protein product [Anisakis simplex]

Sequence ID: VDK56627.1 Length: 289 Number of Matches: 1

Range 1: 16 to 276 GenPept Graphics ▼ Next Match ▲ Previous Match

| Score | Expect | Method | Identities | Positives | Gaps |
|---|---|---|---|---|---|
| 146 bits(368) | 4e-40 | Compositional matrix adjust. | 92/272(34%) | 145/272(53%) | 29/272(10%) |

```
>VDK56627.1 unnamed protein product [Anisakis simplex]
MITPGYGDVSRNMIQLKCSMRRATIAAVAIWMIIFIYLAISIFTLQSKEREDRNEPQKKTDPLVIKYEES
RKKIISLYENIDELKRLLSEKDALLSGNPKLKLIPIAGNGKSESRDEIPHTLYTKDHEMARRNLDNSIKE
LFYYLNSQFEHAKYLSFANHAINQTLSLIAQSAAFSSVDRADRWREKALASISEEMQLHLDKLQHPSDCA
TARALICTLNKGCGFGCQLHHVTYCFIVAYGTNRTLILHRDGKEWNYSERGWSAAFRPVSGCKHDQISKV
LHSDLCLRL
```

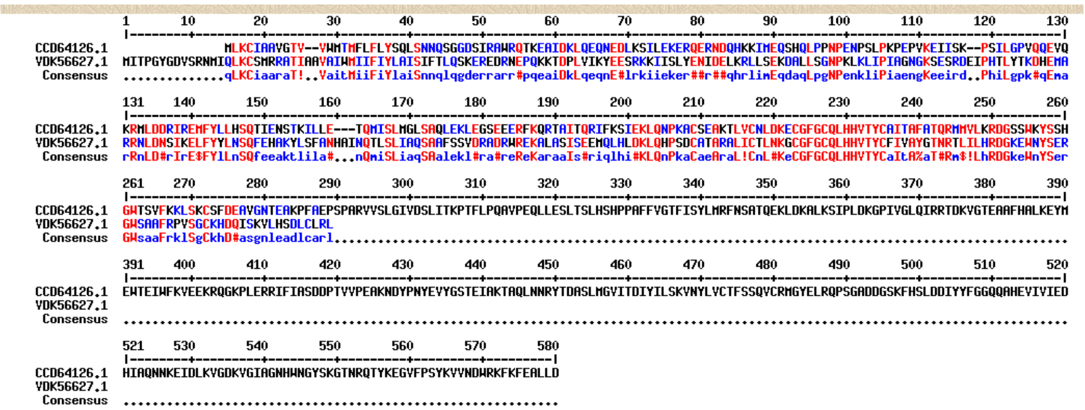

Merged:??

```
MITPGYGDVSRNMIQLKCSMRRATIAAVAIWMIIFIYLAISIFTLQSKEREDRNEPQKKTDPLVIKYEES
RKKIISLYENIDELKRLLSEKDALLSGNPKLKLIPIAGNGKSESRDEIPHTLYTKDHEMARRNLDNSIKE
LFYYLNSQFEHAKYLSFANHAINQTLSLIAQSAAFSSVDRADRWREKALASISEEMQLHLDKLQHPSDCA
TARALICTLNKGCGFGCQLHHVTYCFIVAYGTNRTLILHRDGKEWNYSERGWSAAFRPVSGCKHDQISKV
LHSDLCLRLMRYLMRPNSSLAKRITEFAAKVSFESGPIVGLQIRRTDKVGTEAEFHALSEYMKWTEYWFRIQEYR
HGKAVKRRIYVATDDPTVFSEARKKYPNYEVFGDAAISNTANTRSRYSIESLYGVIIDIEMLARCDYLVCTFSS
QVCRMGYELMQIRVGDAGDNFHSLDDLYYYGGQQAHEQVVVESYQAESKDEIDLEIGDTIGIAGNHWDGF
SKGTNRRNGAVGLYPSYKTREKWIIVPFP
```

```
# WEBSEQUENCE Length: 528
# WEBSEQUENCE Number of predicted TMHs:  1
# WEBSEQUENCE Exp number of AAs in TMHs: 22.53394
# WEBSEQUENCE Exp number, first 60 AAs:  22.29921
# WEBSEQUENCE Total prob of N-in:        0.96568
# WEBSEQUENCE POSSIBLE N-term signal sequence
WEBSEQUENCE     TMHMM2.0        inside       1    22
WEBSEQUENCE     TMHMM2.0        TMhelix     23    45
WEBSEQUENCE     TMHMM2.0        outside     46   528
```

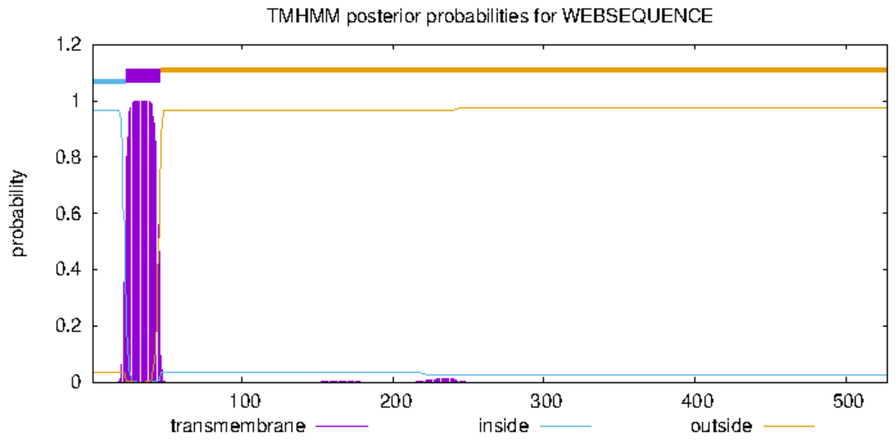


```
            1       10        20        30        40        50        60        70        80        90        100       110       120       130
            |--------+---------+---------+---------+---------+---------+---------+---------+---------+---------+---------+---------+---------|
CCD64126.1                MLKCIAAVGTV--VWMTMFLFLYSQLSNNQSGGDSIRAWRQTKEAIDKLQEQNEDLKSILEKERQERNDQHKKIMEQSHQLPPNPENPSLPKPEPVKEIISK--PSILGPVQQEVQ
AniMerged   MITPGYGDVSRNMIQLKCSMRRATIAAVAIWMIIFIYLAISIFTLQSKEREDRNEPQKKTDPLVIKYEESRKKIISLYENIDELKRLLSEKDALLSGNPKLKLIPIAGNGKSESRDEIPHTLYTKDHEMA
Consensus   ..............qLKCiaaraT!..VaitMiiFiYlaiSnnqlqgderrarr#pqeaiDkLqeqnE#lrkiieker##r##qhrlimEqdaqLpgNPenkliPiaengKeeird..PhiLgpk#qEma

            131     140       150       160       170       180       190       200       210       220       230       240       250       260
            |--------+---------+---------+---------+---------+---------+---------+---------+---------+---------+---------+---------+---------|
CCD64126.1  KRMLDDRIREMFYLLHSQTIENSTKILLE---TQMISLMGLSAQLEKLEGSEEERFKQRTAITQRIFKSIEKLQNPKACSEAKTLVCNLDKECGFGCQLHHVTYCAITAFATQRMMVLKRDGSSWKYSSH
AniMerged   RRNLDNSIKELFYYLNSQFEHAKYLSFANHAINQTLSLIAQSAAFSSVDRADRWREKALASISEEMQLHLDKLQHPSDCATARALICTLNKGCGFGCQLHHVTYCFIVAYGTNRTLILHRDGKEWNYSER
Consensus   rRnLD#rIrE$FYlLnSQfeeaktlila#...nQmiSLiaqSAalekl#ra#reReKaraaIs#riqlhi#KLQnPkaCaeAraL!CnL#KeCGFGCQLHHVTYCaItA%aT#Rm$!LhRDGkeWnYSer

            261     270       280       290       300       310       320       330       340       350       360       370       380       390
            |--------+---------+---------+---------+---------+---------+---------+---------+---------+---------+---------+---------+---------|
CCD64126.1  GWTSVFKKLSKCSFDEAVGNTEAKPFAEPSPARVVSLGIVDSLITKPTFLPQAVPEQLLESLTSLHSHPPAFFVGTFISYLMRFNSATQEKLDKALKSIPLDKGPIVGLQIRRTDKVGTEAAFHALKEYM
AniMerged   GWSAAFRPVSGCKHDQI--------------------------------------SKVLHSDLCLR----------LMRYLMRPNSSLAKRITEFAAKVSFESGPIVGLQIRRTDKVGTEAEFHALSEYM
Consensus   GWsaaFrklSgCkhD#a......................................eqlLeSdlcLr..........lirYLMRfNSalaerideaaak!pl#kGPIVGLQIRRTDKVGTEAaFHALkEYM

            391     400       410       420       430       440       450       460       470       480       490       500       510       520
            |--------+---------+---------+---------+---------+---------+---------+---------+---------+---------+---------+---------+---------|
CCD64126.1  EWTEIWFKVEEKRQGKPLERRIFIASDDPTVVPEAKNDYPNYEVYGSTEIAKTAQLNNRYTDASLMGVITDIYILSKVNYLVCTFSSQVCRMGYELRQPSGADDGSKFHSLDDIYYFGGQQAHEVIVIED
AniMerged   KWTEYWFRIQEYRHGKAVKRRIYVATDDPTVFSEARKKYPNYEVFGDAAISNTANTRSRYSIESLYGVIIDIEMLARCDYLVCTFSSQVCRMGYELMQIRVGDAGDNFHSLDDLYYYGGQQAHEQVVVES
Consensus   eWTEiWFr!#EkRqGKaleRRI%!AsDDPTVfpEArndYPNYEV%GdaaIanTA#lrnRYsdaSLmGVIiDIeiLarc#YLVCTFSSQVCRMGYELrQirgaDaGdnFHSLDDiYY%GGQQAHEq!V!Ed

            521     530       540       550       560       570       580
            |--------+---------+---------+---------+---------+---------|
CCD64126.1  HIAQNNKEIDLKVGDKVGIAGNHWNGYSKGTNRQTYKEGVFPSYKVVNDWRKFKFEALLD
AniMerged   YQAESKDEIDLEIGDTIGIAGNHWDGFSKGTNRRNGAVGLYPSYKTREKWIIVPFP
Consensus   hqA#nndEIDLe!GDk!GIAGNHW#G%SKGTNRrngaeGl%PSYKtr#dWrifkFe....
```

### GALT-1

NCBI Reference Sequence: NP_504545.2

```
>NP_504545.2 Beta-1,4-galactosyltransferase galt-1 [Caenorhabditis elegans]
MPRITASKIVLLIALSFCITVIYHFPIATRSSKEYDEYGNEYENVASIESDIKNVRRLLDEVPDPSQNRL
QFLKLDEHAFAFSAYTDDRNGNMGYKYVRVLMFITSQDNFSCEINGRKSTDVSLYEFSENHKMKWQMFIL
NCKLPDGIDFNNVSSVKVIRSTTKQFVDVPIRYRIQDEKIITPDEYDYKMSICVPALFGNGYDAKRIVEF
IELNTLQGIEKIYIYTNQKELDGSMKKTLKYYSDNHKITLIDYTLPFREDGVWYHGQLATVTDCLLRNTG
ITKYTFFNDFDEFFVPVIKSRTLFETISGLFEDPTIGSQRTALKYINAKIKSAPYSLKNIVSEKRIETRF
TKCVVRPEMVFEQGIHHTSRVIQDNYKTVSHGGSLLRVYHYKDKKYCCEDESLLKKRHGDQLREKFDSVV
GLLDL
```

```
# NP_504545.2 Length: 425
# NP_504545.2 Number of predicted TMHs:  1
# NP_504545.2 Exp number of AAs in TMHs: 20.35882
# NP_504545.2 Exp number, first 60 AAs:  20.35793
# NP_504545.2 Total prob of N-in:        0.98199
# NP_504545.2 POSSIBLE N-term signal sequence
NP_504545.2     TMHMM2.0        inside       1     8
NP_504545.2     TMHMM2.0        TMhelix      9    28
NP_504545.2     TMHMM2.0        outside     29   425
```

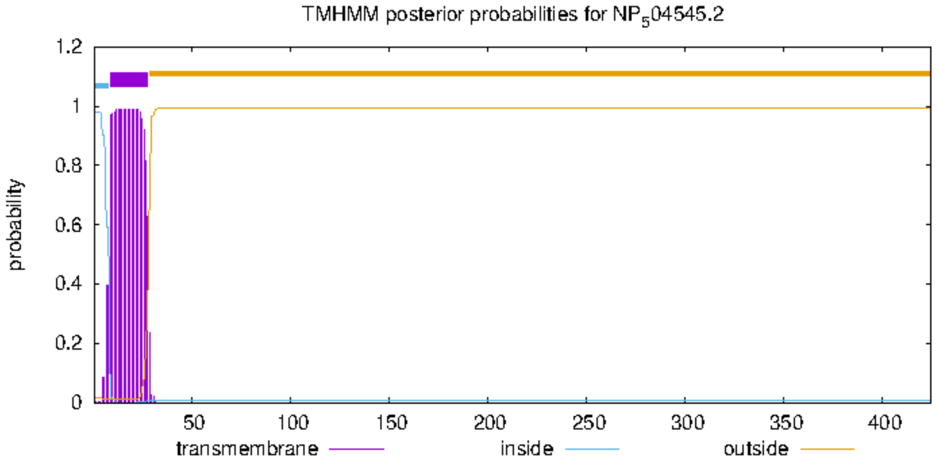


**unnamed protein product [Anisakis simplex]**

Sequence ID: VDK50184.1 Length: 411 Number of Matches: 1

Range 1: 126 to 372 GenPept Graphics ▼ Next Match ▲ Previc

| Score | Expect | Method | Identities | Positives | Gaps |
|---|---|---|---|---|---|
| 115 bits(289) | 6e-29 | Compositional matrix adjust. | 94/341(28%) | 137/341(40%) | 124/341(36%) |

```
>VDK50184.1 unnamed protein product [Anisakis simplex]
MTRLYIKLFLFIVLVFLFTIVFLFNQFQAYPAEVDYFDDHHIDAAFDSRHRAKHLNIVPHPSTRLSHLSV
SLSSDSLSQKIRSPNRGAAIRANRHRKKQQPRTQLEQPLEPGEFRSRKPNVPPRLLTSFVDKRNGNMDYA
CIRAGVVKYAYREARYWCKLDLVSDGGDKQIQIEASLYELAENHSQLYGTFILSCCCCSSNARVYSRSNT
SSSDQHHPAGNSDLDLDKIYEWTLTDGIHESRIPITYRIPAQTPIGTYSVEYAVCVPYLFGTVYSKERLV
ELLELVEYTPPVEDIWYHGQLITITDCLYRNETLITVNNVKSQPRLDNRFTKCVLRPEMVFEQGIHHTSR
VIQDHYKMPYMSPDDAILHHYKKDIANITSITSITSNHAQRFSARLNRTYQRTLQKIDQIN
```

```
# VDK50184.1 Length: 411
# VDK50184.1 Number of predicted TMHs:  1
# VDK50184.1 Exp number of AAs in TMHs: 21.47254
# VDK50184.1 Exp number, first 60 AAs:  21.42452
# VDK50184.1 Total prob of N-in:        0.99352
# VDK50184.1 POSSIBLE N-term signal sequence
VDK50184.1      TMHMM2.0        inside       1     4
VDK50184.1      TMHMM2.0        TMhelix      5    27
VDK50184.1      TMHMM2.0        outside     28   411
```

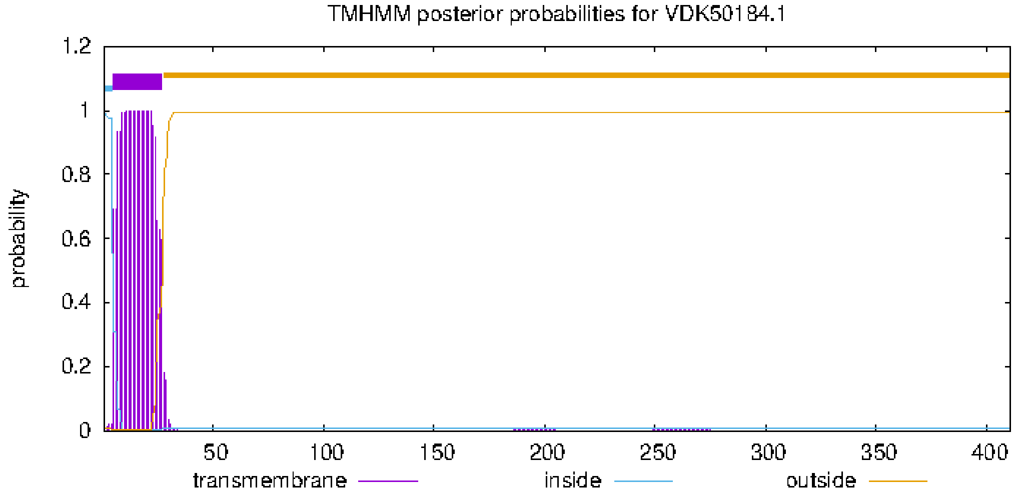


```
             1        10        20        30        40        50        60        70        80        90        100       110       120       130
             |--------+---------+---------+---------+---------+---------+---------+---------+---------+---------+---------+---------+---------|
NP_504545.2  MPRITASKIVLLIALSFCITVIYHFPIATRSSKEYDEYGNEYENVASIESDIKNVRRLLDEVPDPSQNRLQFLKLDEHAFAFSAYTDDRNGNMGYKYVRVLMFITSQDNFSCEINGRKSTDVSLYEFSEN
 VDK50184.1  MTRLYI-KLFLFIVLVFLFTIVFLFNQFQAYPAEVDYFDDHHIDAAF---DSRHRAKHLNIVPHPS-TRLSHLSVSLSSDSLSQKIRSPNRGAAIR--------ANRHRKKQQPRTQLEQPLEPGEF-RS
  Consensus  MpRita.KifLlIaLsFciT!!%hFnqaqaspaEyDe%d#ehe#aAf...DirnrarhL#eVPdPS.nRLqhLkldehadalSakirdrNrnaair........anrdrkkc#irgrleqdlelgEF.rn

             131     140       150       160       170       180       190       200       210       220       230       240       250       260
             |--------+---------+---------+---------+---------+---------+---------+---------+---------+---------+---------+---------+---------|
NP_504545.2  HKMKWQMFILNCKLPDGIDFNNVSSVKVIRSTTKQFVDVPIRYRIQDEKIITPDEYDYKMSICVPALFGNGYDAKRIVEFIELNTLQGIEKIYIYTNQKELDGSMKKTLKYYSDNHKITLIDYTLPFRED
 VDK50184.1  RKPNVPPRLLTSFVDKR---NGNMDYACIRAGVVKYAYREARYWCKLDLVSDGGDKQIQIEASLYELAEN-HSQLYGTFILSCCCCSSNARVYSRSNTSSSD-QHHPAGNSDLDLDKIYEWTLTDGIHES
  Consensus  rKmnwqmriLncklddr...NnnmdyacIRagtkq%adreaRYrcqd#l!idgd#k#iqieaclpaLaeN.hdalrgteiiecnccqgnar!YirsNqkelD.qhhkagnsdlDndKIteidlTdgirEd

             261     270       280       290       300       310       320       330       340       350       360       370       380       390
             |--------+---------+---------+---------+---------+---------+---------+---------+---------+---------+---------+---------+---------|
NP_504545.2  GVWYHGQLATVTDCLLRNTGITKYTFFNDFDEFFVPVIKSRTLFETISGLFEDPTIGSQRTALKYINAKIKSAPYSLKNIVSEKRIETRFTKCVVRPEMVFEQGIHHTSRVIQDNYKTVSHGGSLLRVYH
 VDK50184.1  RIPITYRIPAQTPIGTYSVEYAVCVPYLFGTVYSKERLVELLELVEYTPPVEDIWYHGQLITITDCLYRNETL-ITVNNVKSQPRLDNRFTKCVLRPEMVFEQGIHHTSRVIQDHYKMPYMSPDDAILHH
  Consensus  r!pihgriaaqTdcglrnteiakctf%ndgde%fkerikerleleeisglfEDitiggQriaikdcnarnesa.islnN!kS#kRi#nRFTKCVlRPEMVFEQGIHHTSRVIQDnYKmpshggddarlhH

             391     400       410       420       431
             |--------+---------+---------+---------+|
NP_504545.2  YKDKKYCCEDESLLKKRHGDQLREKFDSVVGLLDL
 VDK50184.1  YKKDIANITSITSITSNHAQRFSARLNRTYQRTLQKIDQIN
  Consensus  YKddiancedeslikkrHa#rlrarl#rtyqrldq......
```

### GLY-12

NP_741838.1

>GLY-12
MKRVLRRSASKHASTLFKIICILVLCSFIIYLKSEDHDLQKNTAIHAPRSNSREDEDDILIPADNKDQQFVAAVLVFCAT
RPDALRNHLSQILAQRPSHFQYHIIVSQDGNKTAVTQVAQKFVKDYKNVSHIQHEKTEIKKRNNYPAISAHYKWALDKAF
KGFRYDHVIVTEDDLDIGNDFFSYFRWGKQVLNSDDTIWCVSAWNDNGGGPLIDSTRGDLIWRTDFFPGLGWMLTKKLWN
ELSPGFPVAYWDDWMRKPEVRKSRSCIRPEISRTSHNMKLAGKGSSGGMFKDYLSKISASSANIDFSLLPVTLVQKSIYD
KRLIEQIENARPIDLQNTTGMEKTYNYKIVYKNIRDWHRLAAHFKLMTDIRGGMQRTAYYGVVTLMFQNCRVFLVPESTY
RNPSQLTSYVYDSEWDKQNRFIEFEAYYCKTKKYAGKCDPHSPEMIAFFKKKGWKKRLDDWGEMLVV

**Sequences producing significant alignments** Download Select columns Show 100

select all 3 sequences selected GenPept Graphics Distance tree of results Multiple alignment MSA Viewer

| | Description | Scientific Name | Max Score | Total Score | Query Cover | E value | Per. Ident | Acc. Len | Accession |
|---|---|---|---|---|---|---|---|---|---|
| ☑ | unnamed protein product [Anisakis simplex] | Anisakis simplex | 214 | 310 | 65% | 2e-64 | 41.97% | 478 | VDK49579.1 |
| ☑ | unnamed protein product [Anisakis simplex] | Anisakis simplex | 192 | 192 | 52% | 2e-58 | 38.91% | 263 | VDK67433.1 |
| ☑ | unnamed protein product [Anisakis simplex] | Anisakis simplex | 80.5 | 80.5 | 12% | 3e-19 | 61.11% | 57 | VDK30835.1 |

**Putative conserved domains have been detected, click on the image below for detailed results.**

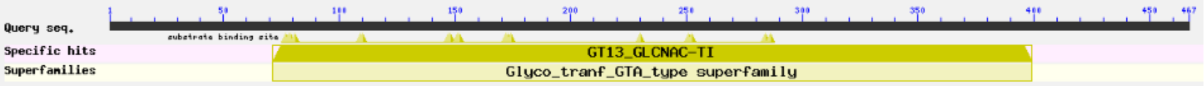


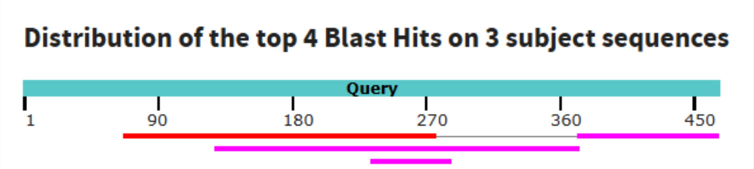


>VDK49579.1 unnamed protein product [Anisakis simplex]

MHSQSIFAMKIFSLVPTKLRRRLCGSVRTRQLIAVCSLTAVVFLLLHQGPLFSSVDEKPQMNDRKILESD

NVVVVPQVVDADNGDVKRLKHPKNGVRFTKQQSELSSGSNTVTDDQPVTVVLVICAIRPAAIKNHLQQLI

RLFLNYFLERISCSCLLRPSSKAFPIVVSQDGTLSSVTNVIKEFVNETEHIHFIQNIADCSSFVTVLLYQ

IKLFYCLSNDINQLSVKCGSYVLFFQHTERKAQNNEAAKKAKNYFFIAQHYKWALDKIFFEMNYDTAIIT

EDDLDIAEDFFSYFSATRYLLRSDPTIWCISAWNDNGGNNITDRSRSDILYRTDFFPGLGWELTVDLWRE

LSAKWPLTYWDDWLRRQDVRQGRVC**IRPE**SG**MARTAY**YGIVSFMLGSSRVYAISAHVYADPNFLSRPSSD

FYTEDWDKMSRYLDFQETYCKPGKFTGDCDPNNPKLKEWFAKKRLTKRLQSWGEMIVN

>VDK67433.1 unnamed protein product [Anisakis simplex]

MQHISGEKANVTAAPNMGRYLTYYMIARHYKLGLSYVFDTLGYKSVIITEDDLDIAPDFFDYFSATRYLL

ERDRTLYCVSAWNDNGKPNLIDTNAYDLLHRSDFFPGLGWMMTSELWQELGPIWPTGFWDDWIRDPARRN

NRS**CIRPE**ISRTAMTPEGKKGASNGLFFVNHLNKIVLNTKPVNFTLLNLDYLLKENYDQHFIDHVYSTPL

LPFSKLRARIARGVGNKSREYRIEYDRLERYLDIAKQLQIMGDFKVGRRLYKS

>VDK30835.1 unnamed protein product [Anisakis simplex]

MTVDLWRELSAKWPLTYWDDWLRRQDVRQGRVC**IRPE**VSRTAHNNKVAGKGTSGYVV

### GLY-13

NP_509566.1

>GLY-13
MHAVTKIFIIFIFVFILWTLYVENDITNRTRNTDNIDDLLESANRLERLLKFEAKKIAALAEDVHKIRANRKGKHVIMEE
MVSQDLKQWKDPIPVLVFSCNRAMAVRDHVEKLIRYRPSQEKFPIIVTQDCDNENVKNEVKKFGDKVEYIKHLAGDKANI
TIPPSHRQYTAYYRIARHYKLALNHVFVDKGYSSVIITEDDLDISPDFFSYFSSTRYLLENDEKLWCVTAWNDNGKQENI
DMTAASTLYRSDFFAGLGWMMSSKTWHELEPIWPVGFWDDWMRDPARRKDRQCIRPEISRTGMMSYGKEGASKGQFFSKH
LAKIKVNDKYINFGKIDLDYLLPANFAKKTNLEVMKEAVELSIDNVASFVLSSENKGKSVRVMYDGNIDYIRKADKLHIM
HDFKAGVPRTAYDGIVTCFINGIRIYLVPDRTKVSAYNPDWSVPPSFGE

select all   3 sequences selected   GenPept   Graphics   Distance tree of results   Multiple alignment   MSA Viewer

| | Description | Scientific Name | Max Score | Total Score | Query Cover | E value | Per. Ident | Acc. Len | Accession |
|---|---|---|---|---|---|---|---|---|---|
| ☑ | unnamed protein product [Anisakis simplex] | Anisakis simplex | 287 | 287 | 57% | 1e-95 | 54.86% | 263 | VDK67433.1 |
| ☑ | unnamed protein product [Anisakis simplex] | Anisakis simplex | 181 | 181 | 51% | 4e-52 | 34.93% | 478 | VDK49579.1 |
| ☑ | unnamed protein product [Anisakis simplex] | Anisakis simplex | 55.5 | 55.5 | 9% | 2e-10 | 50.00% | 57 | VDK30835.1 |

**Putative conserved domains have been detected, click on the image below for detailed results.**

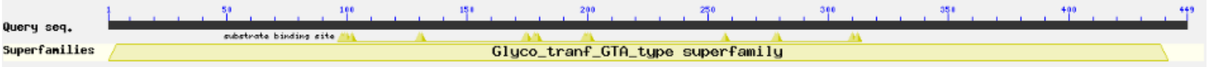


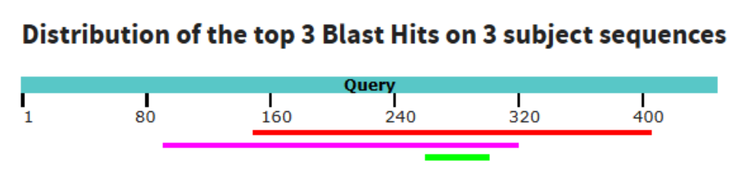


263 aa
>VDK67433.1 unnamed protein product [Anisakis simplex]
MQHISGEKANVTAAPNMGRYLTYYMIARHYKLGLSYVFDTLGYKSVIITEDDLDIAPDFFDYFSATRYLL
ERDRTLYCVSAWNDNGKPNLIDTNAYDLLHRSDFFPGLGWMMTSELWQELGPIWPTGFWDDWIRDPARRN
NRSCIRPEISRTAMTPEGKKGASNGLFFVNHLNKIVLNTKPVNFTLLNLDYLLKENYDQHFIDHVYSTPL
LPFSKLRARIARGVGNKSREYRIEYDRLERYLDIAKQLQIMGDFKVGRRLYKS

### GLY-14

NP_497719.1, 437 aa

>GLY-14
MHISSKITFIFVLIYFSWNFYIQNGFTNKAIVKEELRLLKEMDEAGSRLEKLLKVEKEHMERLSETMNRATRLRITPNSN
NPNNWPHPIPVIVFSCNRPDSVRAHVEKLIMYRPSAQQFPITVSQDCDNESVKKEVEKFGNSVNYVKHPPGESVKIEIPT
NLQKFKPYYYISRHYKLALNHIFSNSNNYSSVIITEDDLDIAPDFFSYFSNTRYLLEKDPSLWCVTAWNDNGKPENIDLK
SNATLYRSDFFAGLGWMMTRKTWEELEPIWPNGFWDDWMREPVQRKQRQCIRPEISRTGMMKYGKEGTSKGQFFSDHLEK
IKVNDLPVDFSQINLDYLQKNEFESRLSLDIRNAVPVDIDDITYPDWKPDYEGMKAIIYYTGRTDFVAKADRLSLMHDFK
AGVPRTAYNGIVTCFYKGTRIFLVPDRSKVPGYDSSW

select all 3 sequences selected GenPept Graphics Distance tree of results Multiple alignment MSA Viewer

| Description | Scientific Name | Max Score | Total Score | Query Cover | E value | Per. Ident | Acc. Len | Accession |
|---|---|---|---|---|---|---|---|---|
| unnamed protein product [Anisakis simplex] | Anisakis simplex | 252 | 252 | 46% | 4e-82 | 57.79% | 263 | VDK67433.1 |
| unnamed protein product [Anisakis simplex] | Anisakis simplex | 178 | 178 | 53% | 4e-51 | 36.18% | 478 | VDK49579.1 |
| unnamed protein product [Anisakis simplex] | Anisakis simplex | 54.7 | 54.7 | 12% | 3e-10 | 49.06% | 57 | VDK30835.1 |

**Putative conserved domains have been detected, click on the image below for detailed results.**

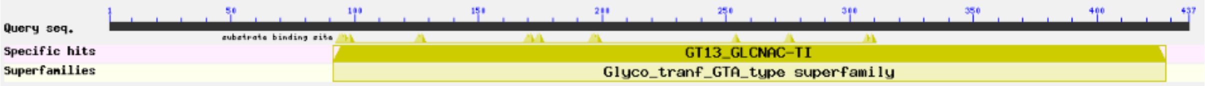


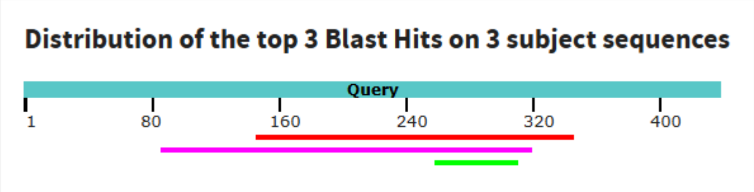

#### GLY-2

NP_491874.1, 669 aa

```
>GLY-2
MRRRHRCVALLFIFSAFITPLGFFYYTISNESKRYSEESEKNYGYQTLEFTESPEEISVDFDKYSQSECSRFPSNVEIEY
PECLNKMKWIKNGWKTHHCYIENHIDGSECSFRYYLSQVENYCPPMEHHGKRKGLAKISPSIRRLLPIFESIPHYMKTRI
NRLWKKWKEGAHEVMQKYPKSMIERRKLNVLVFIGFLANEQKLNMAKKSDHGGPLGELLQWSDLLATLSVIGHHLEVSTN
KNTLRNIVWKYMSRGPCQYVNNFRQQLDIIFTDIMGFNILRQHHRQFLLSNRCRIRLLDSFGTHAEFTTKTYFVQNKKSL
SGPFSQRNPWGGHGLDLRQHWTFYPHSDDNTFLGFVVDTEGIDKKNNQMIPSALVYGKEQYMWRDAEKPIDVLKRIVTVH
STVADLDLKDSNISSIFKKVQNHGFLNSEEISQLLDNITIFFGLGFPLEGPAPLEAMAHGAVFINAKFKEPKSRLNYKFL
AEKPTLRKWTSQNPYMEKIGEPHVITVDIFNELELEEAIKRAISLKPKHFVPFEFTPAGMLHRVALLLEKQELCDKIAYS
KRWPPIDQMKIFRTLNADDSCETICHSKQLLCEPSYFPIINSSPLLRRENLCSSTTSDSSPFAPFNCTIQQSAFLFSCAS
SPPISFEINRLCPCRDYIPEQHAICKKCL
```

```
# GLY-2 Length: 669
# GLY-2 Number of predicted TMHs:  1
# GLY-2 Exp number of AAs in TMHs: 22.05886
# GLY-2 Exp number, first 60 AAs:  22.05011
# GLY-2 Total prob of N-in:        0.99994
# GLY-2 POSSIBLE N-term signal sequence
GLY-2   TMHMM2.0        inside       1     6
GLY-2   TMHMM2.0        TMhelix      7    29
GLY-2   TMHMM2.0        outside     30   669
```

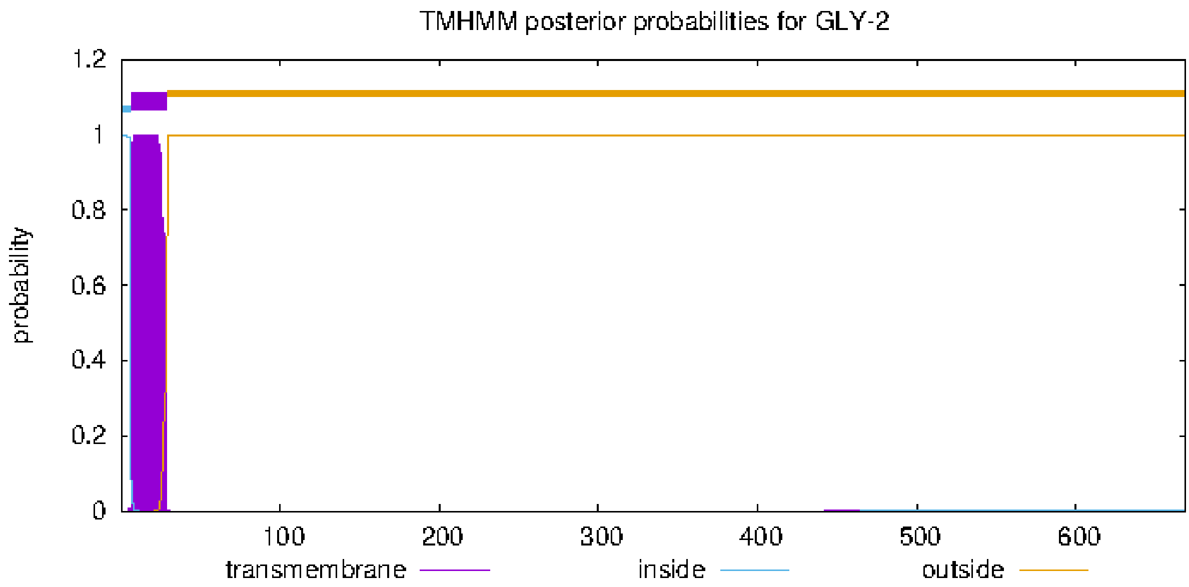


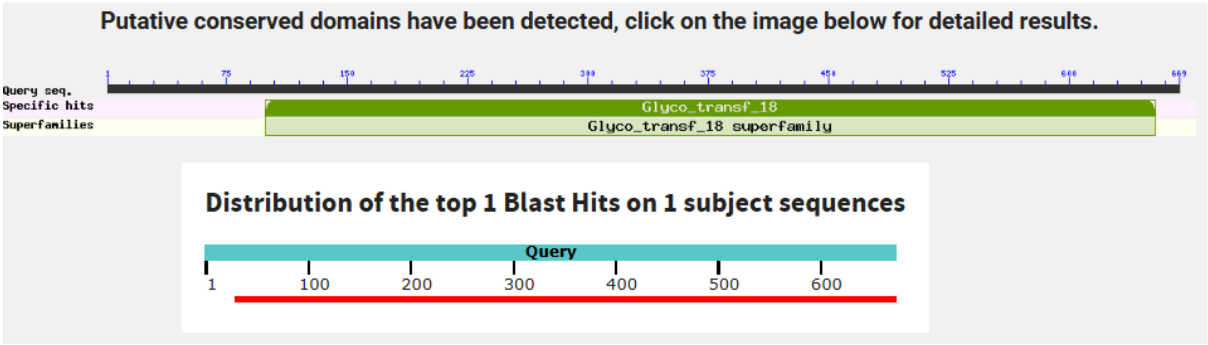

1268 aa

>VDK44765.1 unnamed protein product [Anisakis simplex]
MLPVNNLLFSVILIKIFPVSKVLQVAIFDSTFLRKNMVLMISDDYSLSSHLSEAGSKLVVIDFYADWCGP
CRYIAPIFEQFSQQYTEAVFVKVNVDLCRQTSAVYGVSAMPTFVFVRNNQEVDRMMGADVSELERKIMQH
LSRNTTSLDSQKVATHQERQFLEKFVQFSHRMEVYEDEVAQTLALSVMPSEKLKEKSCNLGEVNQLELLK
NLLRWFKVDFFNWVDTPKCEMCGTVTSSSSRIKGMPTDEEKEFGADRVEVYNCQHCSKEVRFPRYNDPVK
LLETRRGRCGEWANCFALCCRALQFETRWVHDESDHVWCEVWMNDMDRWIHCDPCENVLDTPLLYEKGWG
KKLNYVIAFGLDHIRDVTWRYTFDHLKTIKRRTACREAILRNFFKKLNARFERVMTAERKKELDRRYLNE
LIEFLSPNLQLRGESSSEYQGRTTGSVEWREKRNELGVKSESCKSTSTILRPNDNELADKIFRQFLFYRL
EYNCAKDEYKRGENVIGGWENLISKHKDVFRKEEVDWKMCYLCRREGKSNGELCWSFDLDGLSIKTFSVQ
LNGITKYEDGNILVLVCCGDTCNMLPESGHLTIESPKDGRIDVKVLFSGGIGQLAWQHAQLFRSELKSCE
ADFIGDDKARNHEHEYLNLKVFQEENSQERGMIGHTVDHAKHKDCSLSDSLLNAFPDCKSKLEWMQSGWK
THSCYAQNGVNGSLCSFVIYLSEVEHHCPVLEWRKRIFKPHERQYIPAKIQRNVSALMDLMFDNDVNYQF
IKGRISRLWPRWLSAYDDNLLRWPKTLINRRKLNIVIHMGFLSKESGFKFGEKATAGGPLGELVQWSDLI
SALYILGHNLFISTEYDAFKHNMATFSNVTPCDVSSNRSLGVVFTDIVGVRYMRRHMKQFFLEKKCLLRV
LDSFGTHAEFNSPLYFSTHKHELGGRTNPWGGNGLELQQFMTMYPHTDDNTFLGFVVETHSVNESSVRTN
DTLVYGKEIYMWNGSEKLLDKVNELSQLHATVADMNEFATQRVFNHGLLNGFEWHSLLRRVKIFLGLGFP
LEGPAPLEAIANGAIFINPIFKPSKSRKSYAFFAEKPTLRELTSQNPYVERFIGRPHVITVDITNMSAVE
KAVREALSYESIPYLPFEFTASGMLQRVNILINKQNFCGKSAFPPNQALKVVYANRSQSCEKACSTNGLI
CERSFFDLLNQDSVVNRTNDCEEIKKIASPLAPYKCHIQAERMLFSCASVPHDSWTERICPCRDFIRGQN
ALCSLCLF

```
# VDK44765.1 Length: 1268
# VDK44765.1 Number of predicted TMHs:  0
# VDK44765.1 Exp number of AAs in TMHs: 12.94758
# VDK44765.1 Exp number, first 60 AAs:  12.90259
# VDK44765.1 Total prob of N-in:        0.58357
# VDK44765.1 POSSIBLE N-term signal sequence
VDK44765.1      TMHMM2.0        outside      1  1268
```

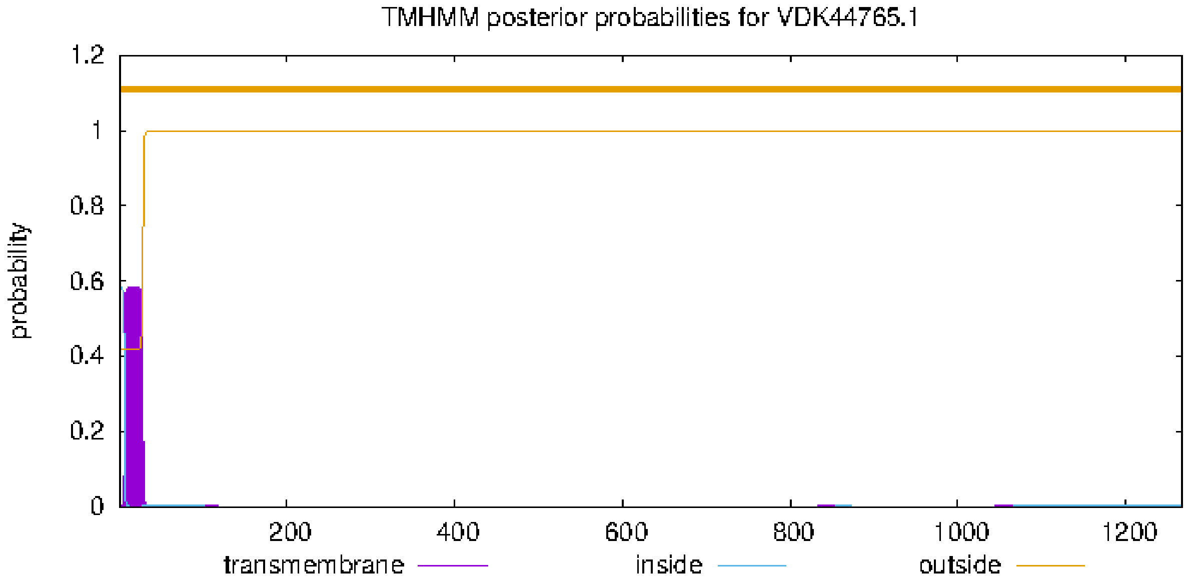

### GLY-20

NP_505864.1, 487 aa

```
>GLY-20
MMVYRRMHRLANAVIACCLFGFIVIFLKAPGEDQRLRDGVPVLSQNPVWSVNDWNGLNKEVVDLLKNESNRLSINEKPDL
SGWNFKTKEDMSLKGSEIVESVSFLNENFDILNAAKFGDLSTVKTILVIQVHDRPVYLQYLIESMRNTKGIEDTLLVFSH
DINVGIINEMIRNITFARVYQIFYPYNLQLFPTVFPGQSPSDCPEKMKRDKAQETNCSNWSSPDKYGNYRVAQLTQIKHH
WWWKMNFVFDGIVEKYSMKDPWVLLLEEDHMLAPDALHVLDIIVSNRPKYCENCEIISLGFYLKSTNKYGQDIAHLGVHP
WYSSKHNMGMALQKNTWQKIKGCSEMFCKWDDYNWDWSLMQISAKCLPQRFRVIFTKSPRVIHIGDCGVHTHRCEAHKAL
QSTQELFRQHKDLLFPTSLSVTDTSRRSLKPSKENGGWGDIRDRQLCEINKSPLVRVSSQSASVLHKLLNSKIQFSSSNK
TITSTTS
```

```
# GLY-20 Length: 487
# GLY-20 Number of predicted TMHs:  1
# GLY-20 Exp number of AAs in TMHs: 19.24938
# GLY-20 Exp number, first 60 AAs:  19.2113
# GLY-20 Total prob of N-in:        0.97542
# GLY-20 POSSIBLE N-term signal sequence
GLY-20  TMHMM2.0        inside       1     6
GLY-20  TMHMM2.0        TMhelix      7    26
GLY-20  TMHMM2.0        outside     27   487
```

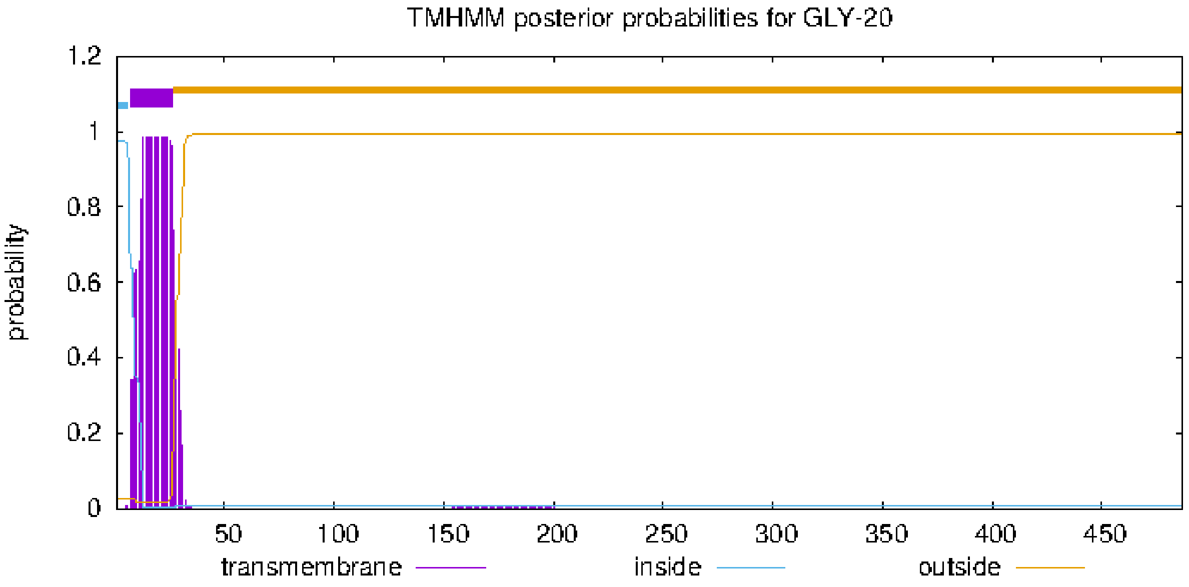


**Putative conserved domains have been detected, click on the image below for detailed results.**

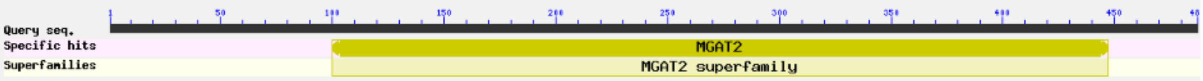


**Distribution of the top 1 Blast Hits on 1 subject sequences**

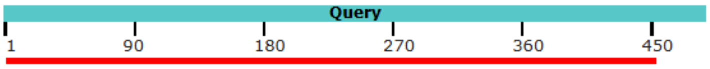

481aa

>VDK17387.1 unnamed protein product [Anisakis simplex]
MIRLKFVRLVNAIIVISFLGLLMVFIQNPYPPVDEHSEAAAAVAAVGSYESRGERLLSANSLPVQCYCCL
KAIEESLKESATFISSPLSINDVVSSIEFLNNNYEILNMHRFGPVSEAKFVIVVQVHNRIEYLRYLIESL
QKAKYINEVLSIFSHDYSSPDINELIKQIRFCRVLQIFYPYNIQLFPTVFPGTDPNDCAADMTKEKAKQQ
RCNNWAHPDKYGHYRVAKLTQIKHHWWWKINYVFDVVMKRYNLDKAWVILLEEDHYVAPDFIHVLQKVID
NKKQYCEQCQVVSLGLYLKQYSTYGDNIASLIVHPWFSSKHNMGMAFDVNTWNLIKNCSKEFCTYDDYNW
DWSLLHISMKCLPKKLRVIALKAPRVIHVGDCGVHTHRCNVQNAPKKAQELFSSVEKRLFPASLRVVENS
RRMLKPSKENGGWGDSRDHQLCINNSMVNVQPSLVDSLVVKGDYDSAIFINLNNSSNIMDL

```
# VDK17387.1 Length: 481
# VDK17387.1 Number of predicted TMHs:  1
# VDK17387.1 Exp number of AAs in TMHs: 20.0794099999999
# VDK17387.1 Exp number, first 60 AAs:  20.07647
# VDK17387.1 Total prob of N-in:        0.99511
# VDK17387.1 POSSIBLE N-term signal sequence
VDK17387.1      TMHMM2.0        inside       1     6
VDK17387.1      TMHMM2.0        TMhelix      7    26
VDK17387.1      TMHMM2.0        outside     27   481
```

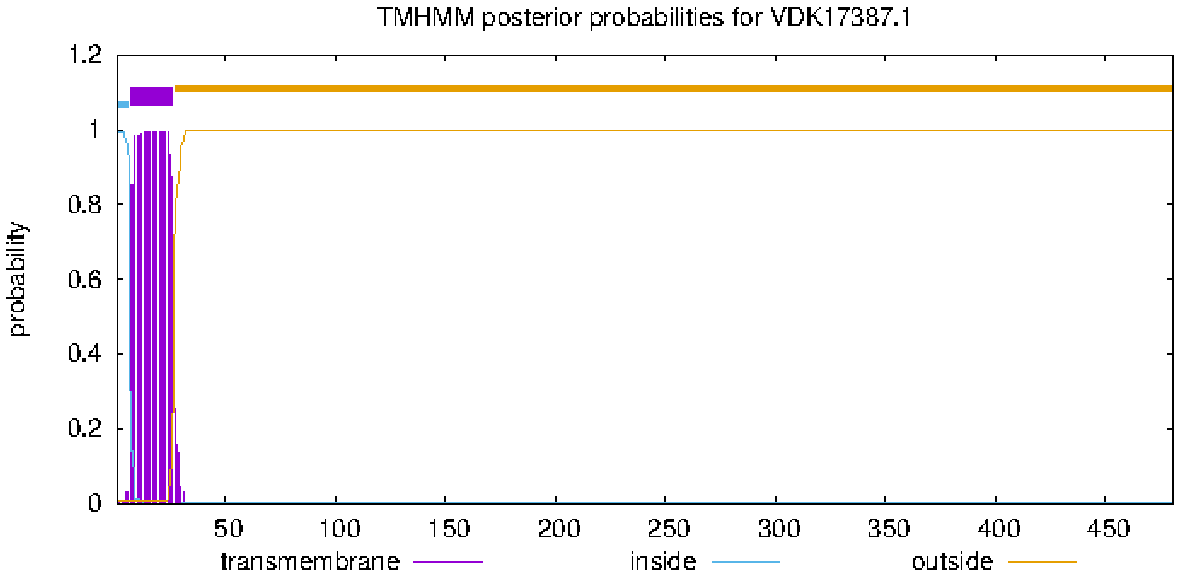

### MGAT3 (human)

NP_001091740.1, 533 aa

>MGAT3
MKMRRYKLFLMFCMAGLCLISFLHFFKTLSYVTFPRELASLSPNLVSSFFWNNAPVTPQASPEPGGPDLLRTPLYSH
SPLLQPLPPSKAAEELHRVDLVLPEDTTEYFVRTKAGGVCFKPGTKMLERPPPGRPEEKPEGANGSSARRPPRYLLS
ARERTGGRGARRKWVECVCLPGWHGPSCGVPTVVQYSNLPTKERLVPREVPRRVINAINVNHEFDLLDVRFHELG
DVVDAFVVCESNFTAYGEPRPLKFREMLTNGTFEYIRHKVLYVFLDHFPPGGRQDGWIADDYLRTFLTQDGVSRLR
NLRPDDVFIIDDADEIPARDGVLFLKLYDGWTEPFAFHMRKSLYGFFWKQPGTLEVVSGCTVDMLQAVYGLDGIRL
RRRQYYTMPNFRQYENRTGHILVQWSLGSPLHFAGWHCSWCFTPEGIYFKLVSAQNGDFPRWGDYEDKRDLNYI
RGLIRTGGWFDGTQQEYPPADPSEHMYAPKYLLKNYDRFHYLLDNPYQEPRSTAAGGWRHRGPEGRPPARGKLD
EAEV

No significant similarity found. For reasons why,click here

### MGAT4 (human)

NP_001253751.1, 548 aa

>MGAT4B

MRLRNGTFLTLLLFCLCAFLSLSWYAALSGQKGDVVDVYQREFLALRDRLHAAEQESLKRSKELNLVLDEIKRAVSER
QALRDGDGNRTWGRLTEDPRLKPWNGSHRHVLHLPTVFHHLPHLLAKESSLQPAVRVGQGRTGVSVVMGIPSV
RREVHSYLTDTLHSLISELSPQEKEDSVIVVLIAETDSQYTSAVTENIKALFPTEIHSGLLEVISPSPHFYPDFSRLRESFG
DPKERVRWRTKQNLDYCFLMMYAQSKGIYYVQLEDDIVAKPNYLSTMKNFALQQPSEDWMILEFSQLGFIGKMF
KSLDLSLIVEFILMFYRDKPIDWLLDHILWVKVCNPEKDAKHCDRQKANLRIRFKPSLFQHVGTHSSLAGKIQKLKDK
DFGKQALRKEHVNPPAEVSTSLKTYQHFTLEKAYLREDFFWAFTPAAGDFIRFRFFQPLRLERFFFRSGNIEHPEDKL
FNTSVEVLPFDNPQSDKEALQEGRTATLRYPRSPDGYLQIGSFYKGVAEGEVDPAFGPLEALRLSIQTDSPVWVILSE
IFLKKAD

```
# MGAT4B Length: 548
# MGAT4B Number of predicted TMHs:  1
# MGAT4B Exp number of AAs in TMHs: 21.83668
# MGAT4B Exp number, first 60 AAs:  21.78962
# MGAT4B Total prob of N-in:        0.99439
# MGAT4B POSSIBLE N-term signal sequence
MGAT4B  TMHMM2.0        inside       1     6
MGAT4B  TMHMM2.0        TMhelix      7    29
MGAT4B  TMHMM2.0        outside     30   548
```

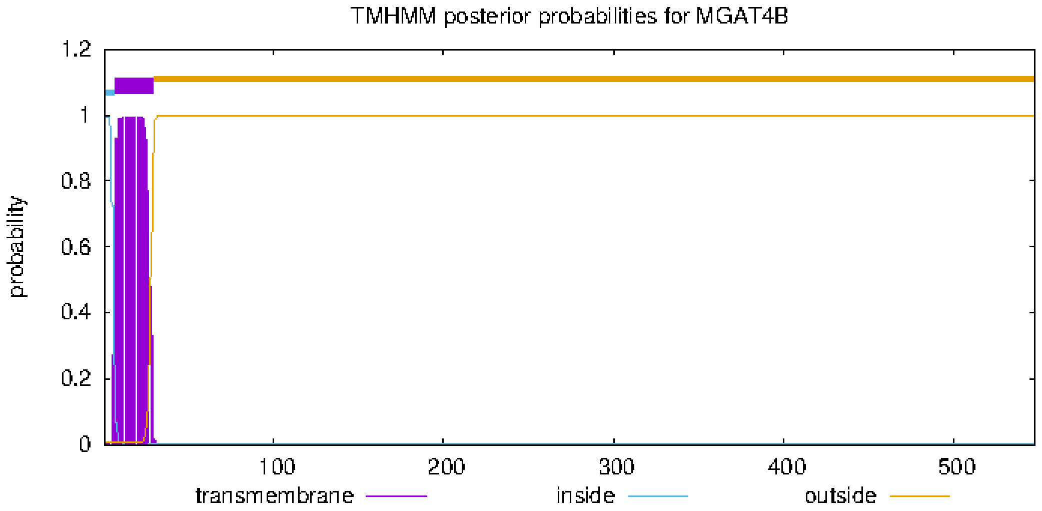


**Sequences producing significant alignments** Download Select columns Show 100

select all *3 sequences selected* GenPept Graphics Distance tree of results Multiple alignment MSA Viewer

| Description | Scientific Name | Max Score | Total Score | Query Cover | E value | Per. Ident | Acc. Len | Accession |
|---|---|---|---|---|---|---|---|---|
| unnamed protein product [Anisakis simplex] | Anisakis simplex | 349 | 349 | 80% | 1e-115 | 43.71% | 468 | VDK42989.1 |
| unnamed protein product [Anisakis simplex] | Anisakis simplex | 252 | 252 | 80% | 2e-77 | 33.76% | 526 | VDK43945.1 |
| alpha-1,3-mannosyl-glycoprotein 4-beta-N-acetylglucosaminyltransferase B [Anisakis pegreffii] | Anisakis pegreffii | 160 | 160 | 49% | 5e-46 | 34.33% | 252 | AZA15233.1 |

**Putative conserved domains have been detected, click on the image below for detailed results.**

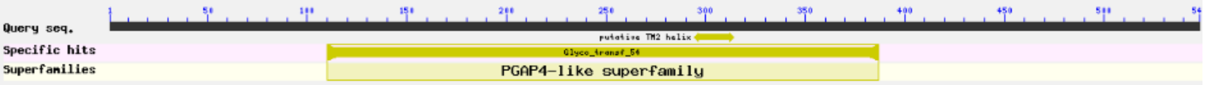


**Distribution of the top 3 Blast Hits on 3 subject sequences**

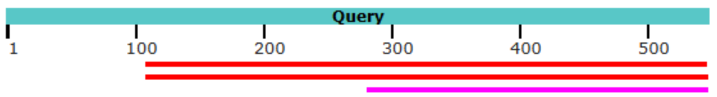

468 aa

>VDK42989.1 unnamed protein product [Anisakis simplex]
MLANVWRRLTDRDRELFLLKNSVTLCLNETPIPQLTISFPSFFDFLPHLYARSPSALRPAVIYPNDTFST
VAKVVIGIPTVARQNFSYLIPTLQSLISGMNAQEKAATLIVVLIADEHGAHSQFVNEQLHSIQSEFATDL
ESVMKRTVDLQLIVPPHSWYPPDLHAVPATFGDSEERVYWRTKQNLDYIFLMLYCQGKGEYYLQLEDDVV
TKPGFMSRIHDFIAQQPSDSWFMLEFSTLGFIGKLFRSSDLPLLTQFIVLFHRAKPVDWLLDLIFVNRYC
HPEKSSKECSQAVKQYRVRARPSLFQHIGVHSSLAGKLQKLREQDFGKVQLYVPHENNPPAVVTTNLVTY
KSHDIEGAYNGRNFFWSLAPHADDHISFDFKPAIKLRGFLFKTGNGEHQRDILNENAVVYLHRKNSKDFE
RITSFNEHGTARASFDGNDVALIDSLKIVVHNDSSNWVIFSEIFIKND

```
# VDK42989.1 Length: 468
# VDK42989.1 Number of predicted TMHs:  0
# VDK42989.1 Exp number of AAs in TMHs: 0.13907
# VDK42989.1 Exp number, first 60 AAs:  0.08231
# VDK42989.1 Total prob of N-in:        0.01293
VDK42989.1      TMHMM2.0        outside      1   468
```

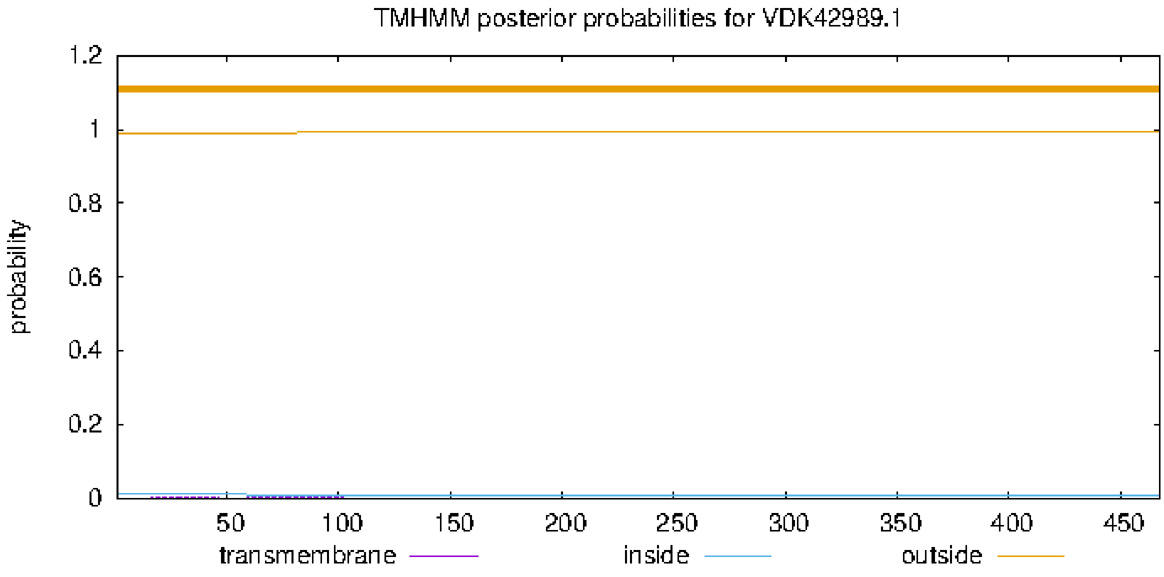

526 aa

>VDK43945.1 unnamed protein product [Anisakis simplex]

MLISPFARFPFRLILLVTLSVITIVCIVQTIPSTDEERYDNSKEMHIDRVSLNFHPSPVHHLDIQPNAPT
SSTPQSVRHKESVQSNDLTLPSLYDLLPHLRNNPDALRPSLVYANHTIRRNARLVIGVPTVPRDNHSYLL
QTLQSLAFGLDDVQQRNANIVIMIGSKDGADTTSVKQQISLIESDFKEFLDNGFFHIIVPPREWYPPDLD
SIEPTLGDPPQRMYWRTKQNLDYMFLLLYCQSLGEYYLQVEDDILAKEHYMDRIINFINEKGNFKWFTAE
FASLGFIGKLFHTKDLLYLINYIALLYRYKPVDWILDSVFMDRYCLPFEKAKPCYKRQKDYRVPGGTLFQ
HIGIHSSLEGKIQKLKEKNFGQSNTFSAHTDNPKAVVNTSLKQFKEYGNFNLNVFKIFMMADRCSGPQFR
RPLEIISASILLIKSMSKSVNLKTFSGFIIRSGNPEHPDDKFDNRTVVLTKSGAEGSAFEKLCTFNERGI
ARHHFAVPTSLQAIRLEVQTNYSNWLLLSELHIRHS

```
# VDK43945.1 Length: 526
# VDK43945.1 Number of predicted TMHs:  1
# VDK43945.1 Exp number of AAs in TMHs: 19.97639
# VDK43945.1 Exp number, first 60 AAs:  19.9062
# VDK43945.1 Total prob of N-in:        0.93646
# VDK43945.1 POSSIBLE N-term signal sequence
VDK43945.1      TMHMM2.0        inside       1     8
VDK43945.1      TMHMM2.0        TMhelix      9    31
VDK43945.1      TMHMM2.0        outside     32   526
```

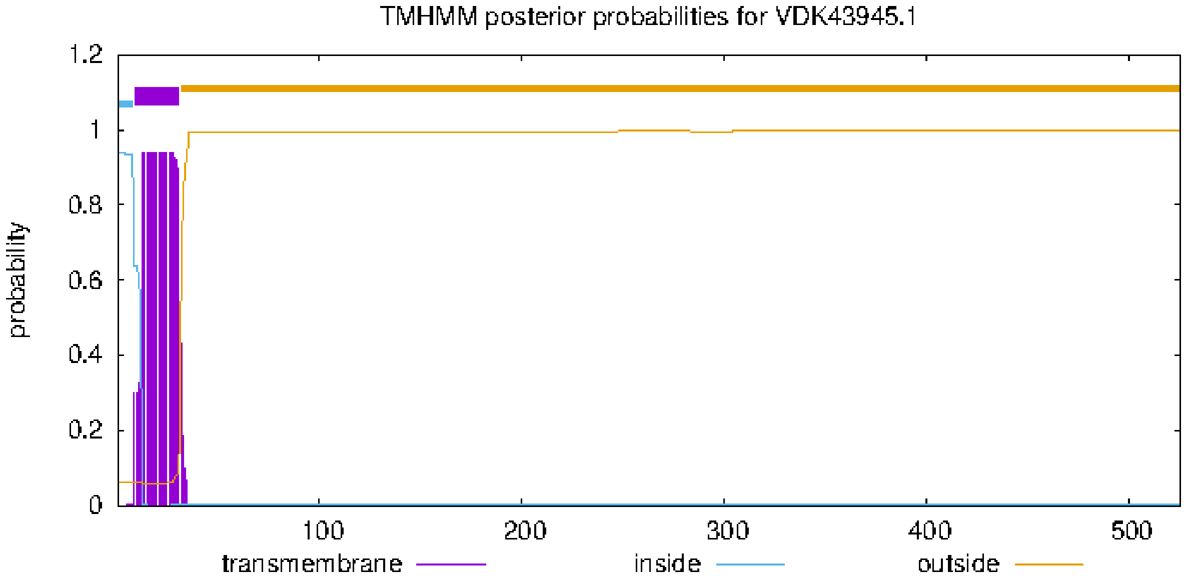

252 aa

>AZA15233.1 alpha-1,3-mannosyl-glycoprotein 4-beta-N-acetylglucosaminyltransferase B [Anisakis pegreffii]

MDRIINFINEKGNFKWFTAEFASLGFIGKLFHTKDLPYLINYIALLYRYKPVDWILDSVFMDRYCLPFEK
AKPCYKRQKDYRVPGGTLFQHIGIHSSLEGKIQKLKEKNFGQSNTFSAHTDNPKAVVNTSLKQFKEYGIQ
NFYDGRSLFWSSVPPTFGDYINIHFTDQVHVKGFIIRSGNPEHPDDKFDNRTVVLTKSGAEGSAFEKLCT
FNERGIARHHFAVPTSLQAIRLEVQTNYTNWLLLSELHIRHS

```
# AZA15233.1 Length: 252
# AZA15233.1 Number of predicted TMHs:  0
# AZA15233.1 Exp number of AAs in TMHs: 0.620799999999999
# AZA15233.1 Exp number, first 60 AAs:  0.61657
# AZA15233.1 Total prob of N-in:        0.09012
AZA15233.1      TMHMM2.0        outside      1   252
```

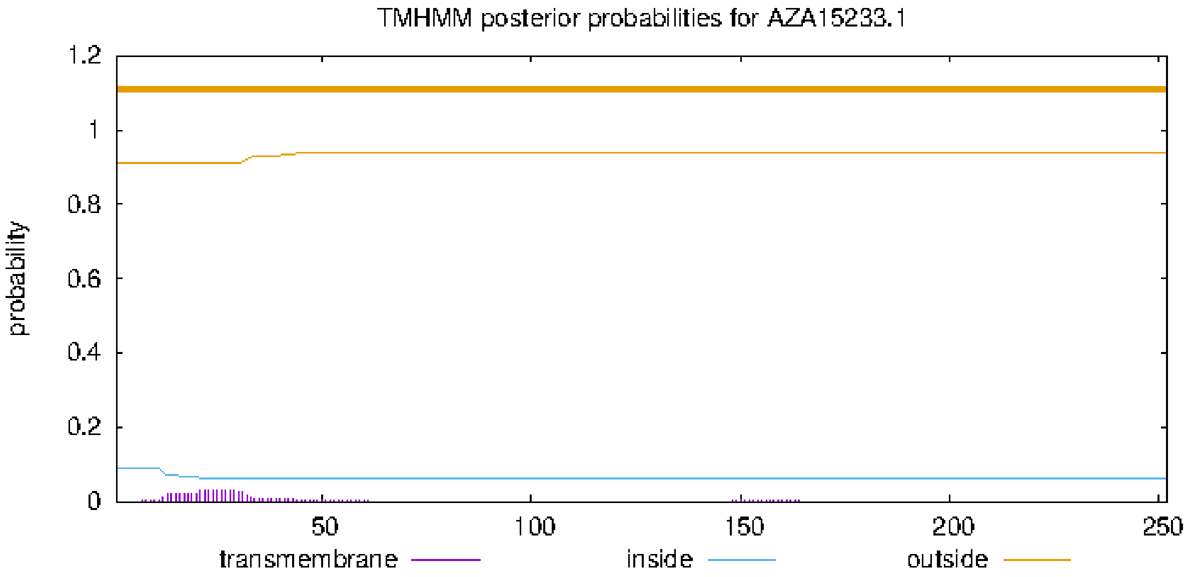

### MGAT5 (human)

NP_001358386.1, 741 aa

>MGAT5
MALFTPWKLSSQKLGFFLVTFGFIWGMMLLHFTIQQRTQPESSSMLREQILDLSKRYIKALAEENRNVVDGPYAGV
MTAYDLKKTLAVLLDNILQRIGKLESKVDNLVVNGTGTNSTNSTTAVPSLVALEKINVADIINGAQEKCVLPPMDGY
PHCEGKIKWMKDMWRSDPCYADYGVDGSTCSFFIYLSEVENWCPHLPWRAKNPYEEADHNSLAEIRTDFNILYS
MMKKHEEFRWMRLRIRRMADAWIQAIKSLAEKQNLEKRKRKKVLVHLGLLTKESGFKIAETAFSGGPLGELVQWS
DLITSLYLLGHDIRISASLAELKEIMKKVVGNRSGCPTVGDRIVELIYIDIVGLAQFKKTLGPSWVHYQCMLRVLDSFG
TEPEFNHANYAQSKGHKTPWGKWNLNPQQFYTMFPHTPDNSFLGFVVEQHLNSSDIHHINEIKRQNQSLVYGKV
DSFWKNKKIYLDIIHTYMEVHATVYGSSTKNIPSYVKNHGILSGRDLQFLLRETKLFVGLGFPYEGPAPLEAIANGCAF
LNPKFNPPKSSKNTDFFIGKPTLRELTSQHPYAEVFIGRPHVWTVDLNNQEEVEDAVKAILNQKIEPYMPYEFTCEG
MLQRINAFIEKQDFCHGQVMWPPLSALQVKLAEPGQSCKQVCQESQLICEPSFFQHLNKDKDMLKYKVTCQSSEL
AKDILVPSFDPKNKHCVFQGDLLLFSCAGAHPRHQRVCPCRDFIKGQVALCKDCL

select all *1 sequences selected* GenPept Graphics Distance tree of results Multiple alignment MSA Viewer

| | Description | Scientific Name | Max Score | Total Score | Query Cover | E value | Per. Ident | Acc. Len | Accession |
|---|---|---|---|---|---|---|---|---|---|
| ☑ | unnamed protein product [Anisakis simplex] | Anisakis simplex | 490 | 490 | 83% | 2e-158 | 43.15% | 1268 | VDK44765.1 |

Putative conserved domains have been detected, click on the image below for detailed results.

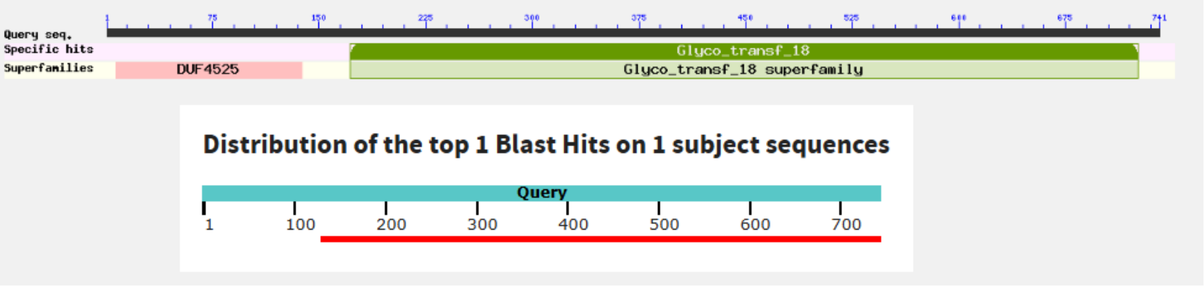


1268 aa (a closer homolog of GLY-2 than MGAT5)

>VDK44765.1 unnamed protein product [Anisakis simplex]
MLPVNNLLFSVILIKIFPVSKVLQVAIFDSTFLRKNMVLMISDDYSLSSHLSEAGSKLVVIDFYADWCGP
CRYIAPIFEQFSQQYTEAVFVKVNVDLCRQTSAVYGVSAMPTFVFVRNNQEVDRMMGADVSELERKIMQH
LSRNTTSLDSQKVATHQERQFLEKFVQFSHRMEVYEDEVAQTLALSVMPSEKLKEKSCNLGEVNQLELLK
NLLRWFKVDFFNWVDTPKCEMCGTVTSSSSRIKGMPTDEEKEFGADRVEVYNCQHCSKEVRFPRYNDPVK
LLETRRGRCGEWANCFALCCRALQFETRWVHDESDHVWCEVWMNDMDRWIHCDPCENVLDTPLLYEKGWG
KKLNYVIAFGLDHIRDVTWRYTFDHLKTIKRRTACREAILRNFFKKLNARFERVMTAERKKELDRRYLNE
LIEFLSPNLQLRGESSSEYQGRTTGSVEWREKRNELGVKSESCKSTSTILRPNDNELADKIFRQFLFYRL
EYNCAKDEYKRGENVIGGWENLISKHKDVFRKEEVDWKMCYLCRREGKSNGELCWSFDLDGLSIKTFSVQ
LNGITKYEDGNILVLVCCGDTCNMLPESGHLTIESPKDGRIDVKVLFSGGIGQLAWQHAQLFRSELKSCE
ADFIGDDKARNHEHEYLNLKVFQEENSQERGMIGHTVDHAKHKDCSLSDSLLNAFPDCKSKLEWMQSGWK
THSCYAQNGVNGSLCSFVIYLSEVEHHCPVLEWRKRIFKPHERQYIPAKIQRNVSALMDLMFDNDVNYQF
IKGRISRLWPRWLSAYDDNLLRWPKTLINRRKLNIVIHMGFLSKESGFKFGEKATAGGPLGELVQWSDLI
SALYILGHNLFISTEYDAFKHNMATFSNVTPCDVSSNRSLGVVFTDIVGVRYMRRHMKQFFLEKKCLLRV
LDSFGTHAEFNSPLYFSTHKHELGGRTNPWGGNGLELQQFMTMYPHTDDNTFLGFVVETHSVNESSVRTN
DTLVYGKEIYMWNGSEKLLDKVNELSQLHATVADMNEFATQRVFNHGLLNGFEWHSLLRRVKIFLGLGFP
LEGPAPLEAIANGAIFINPIFKPSKSRKSYAFFAEKPTLRELTSQNPYVERFIGRPHVITVDITNMSAVE
KAVREALSYESIPYLPFEFTASGMLQRVNILINKQNFCGKSAFPPNQALKVVYANRSQSCEKACSTNGLI
CERSFFDLLNQDSVVNRTNDCEEIKKIASPLAPYKCHIQAERMLFSCASVPHDSWTERICPCRDFIRGQN
ALCSLCLF

Multalin

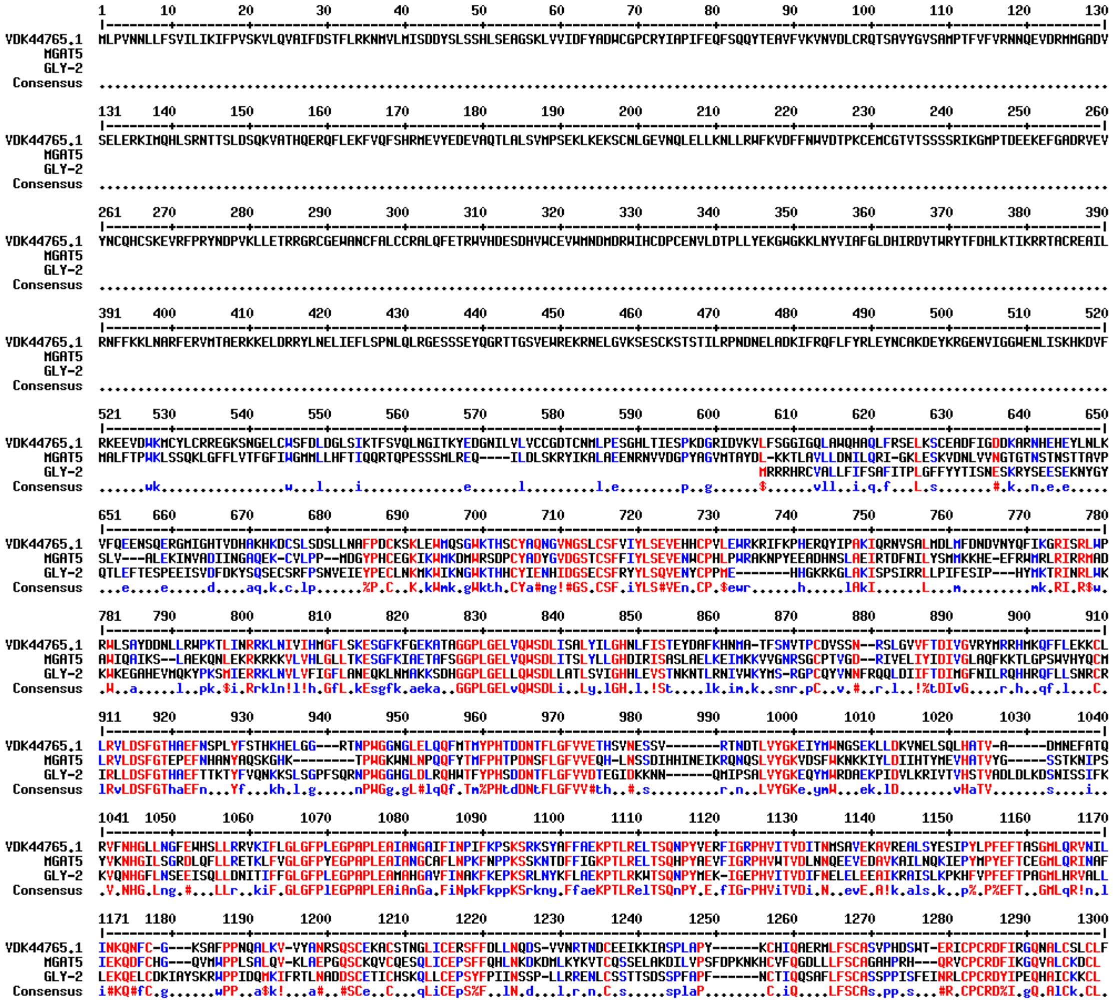

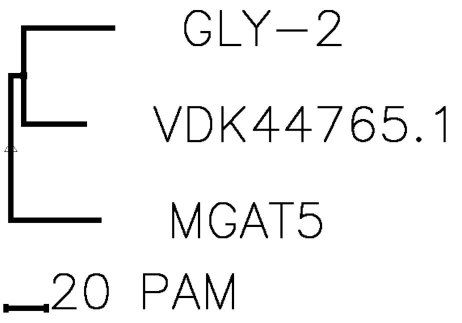

#### BRE-4

NP_490872.1

>NP_490872.1 Beta-1,4-N-acetylgalactosaminyltransferase bre-4 [Caenorhabditis elegans]
MAFRHLAVARLKSLLVLCAVLLLVHAMIYKIPSLYENLTIGSSTLIADVDAMEAVLGNTASTSDDLLDTW
NSTFSPISEVNQTSFMEDIRPILFPDNQTLQFCNQTPPHLVGPIRVFLDEPDFKTLEKIYPDTHAGGHGM
PKDCVARHRVAIIVPYRDREAHLRIMLHNLHSLLAKQQLDYAIFIVEQVANQTFNRGKLMNVGYDVASRL
YPWQCFIFHDVDLLPEDDRNLYTCPIQPRHMSVAIDKFNYKLPYSAIFGGISALTKDHLKKINGFSNDFW
GWGGEDDDLATRTSMAGLKVSRYPTQIARYKMIKHSTEATNPVNKCRYKIMGQTKRRWTRDGLSNLKYKL
VNLELKPLYTRAVVDLLEKDCRRELRRDFPTCF

```
# BRE-4, Length: 383
# BRE-4, Number of predicted TMHs:  1
# BRE-4, Exp number of AAs in TMHs: 21.81185
# BRE-4, Exp number, first 60 AAs:  21.80964
# BRE-4, Total prob of N-in:        0.98883
# BRE-4, POSSIBLE N-term signal sequence
BRE-4,   TMHMM2.0        inside       1     6
BRE-4,   TMHMM2.0        TMhelix      7    29
BRE-4,   TMHMM2.0        outside     30   383
```

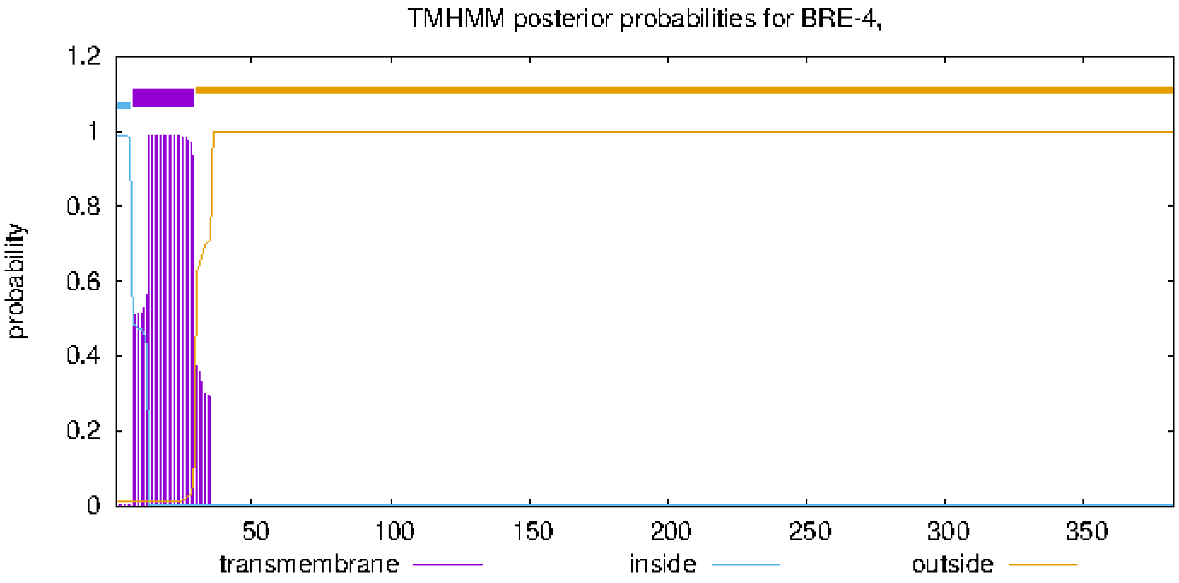

**Sequences producing significant alignments**

Download | Select columns | Show 100

select all 5 sequences selected

GenPept Graphics Distance tree of results Multiple alignment MSA Viewer

| | Description | Scientific Name | Max Score | Total Score | Query Cover | E value | Per. Ident | Acc. Len | Accession |
|---|---|---|---|---|---|---|---|---|---|
| ☑ | unnamed protein product [Anisakis simplex] | Anisakis simplex | 380 | 380 | 69% | 5e-132 | 65.33% | 329 | VDK43914.1 |
| ☑ | unnamed protein product [Anisakis simplex] | Anisakis simplex | 204 | 204 | 62% | 9e-63 | 39.50% | 367 | VDK57682.1 |
| ☑ | unnamed protein product [Anisakis simplex] | Anisakis simplex | 164 | 164 | 50% | 2e-48 | 41.03% | 288 | VDK50333.1 |
| ☑ | unnamed protein product [Anisakis simplex] | Anisakis simplex | 103 | 103 | 55% | 1e-25 | 31.96% | 291 | VDK57694.1 |
| ☑ | unnamed protein product [Anisakis simplex] | Anisakis simplex | 51.2 | 51.2 | 18% | 7e-08 | 33.80% | 198 | VDK65456.1 |

**Putative conserved domains have been detected, click on the image below for detailed results.**

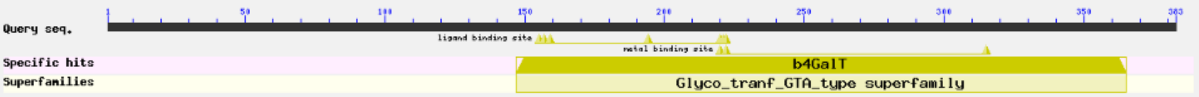


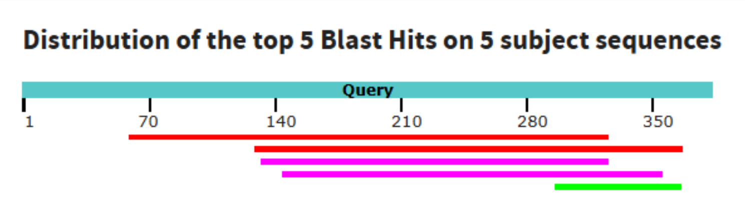


>VDK43914.1 unnamed protein product [Anisakis simplex]
MMNNKLKLVVLLGLCVALVHFLLSDCFISPDYHFWSPSLLTSIPKAFNVVDSSLKFSQSKDTVLSILNSF
DDKAISYSIAASLNTSFVNITDDGLVMEKQDERLAFCPLIPPKLVGPIAVWLDAPSFDDLERLYPKLQAG
GHGMPSGCRARHRVAIIVPYRDRETHLRVLLHNLHSLLMKQQLDYAIFVVEQVSNETFNRAKLMNVGYNE
AIKLHNWQCFIFHDVDLIPEDDRNLYSCPEQPRHMSVAVDKFQYKLPYGSIFGGISALTKEQFIRINGFS
NDYWGWGGEDDDLSTRVSLKGYKISRYPTEIARYKMIKHESEKENPVNK

### HEX-2

**Gene symbol: hex-2**

**Gene description: Beta-N-acetylhexosaminidase**

GenBank: NP_504489.3, 596 aa

```
>NP_504489.3 beta-N-acetylhexosaminidase [Caenorhabditis elegans]
MFPMRCIRRRSIDFVLKGVILTTICLFLFHMTSSYPKGGISQRALDSMQKEPQTPVLVQKRENFEESIDV
AVEQESKKENPFVNQQASTENAKPITQEVKIERPSRDNEFYKNVVIHFDLKGAPPKVDYFLDLLRLIAKG
GATGILLEWEDMFPWTGKLEQFKNTDAYSESDVDMILSEATKLKLDVIPLVQTFGHLEWILKYEEMRKYR
ENDAYPQVLCLGNEEGVEFVREMIRQVAKKHAKYGIPFFHIGADEAFEFGVCQESLDWIKKNGKNGRKQL
LALAHLKAIAEFAKQQTGDSTQILAWHDMLKDFDSRLIKNLELGQIIQPVVWDYSENIITLNDYIFSALA
ENFPTMWASSAYKGANYPSASTSEVRHYETNNRNWIRTKQNQERKFKNGFQGIIVTGWQRYDHLAGLCET
LPIGTASMMLQMQIALNAPALDLEGTRQKAATLLECQGFNVDGVKVVSNQCKYRGFQTYLIYQSEVPNLF
ARIDSELSKNHHLMGWANRYNRKYNISQNWYHREMLPFVQQLVGQYDRVESDLRASMKDLYFENTIDEFI
YENLGEMSEKLHGYLEEIQRLDKLRAWPKRHFPIKK
```

```
# CCD64497.1 Length: 596
# CCD64497.1 Number of predicted TMHs:  1
# CCD64497.1 Exp number of AAs in TMHs: 20.1106
# CCD64497.1 Exp number, first 60 AAs:  20.10864
# CCD64497.1 Total prob of N-in:        0.93438
# CCD64497.1 POSSIBLE N-term signal sequence
CCD64497.1      TMHMM2.0        inside       1    12
CCD64497.1      TMHMM2.0        TMhelix     13    35
CCD64497.1      TMHMM2.0        outside     36   596
```

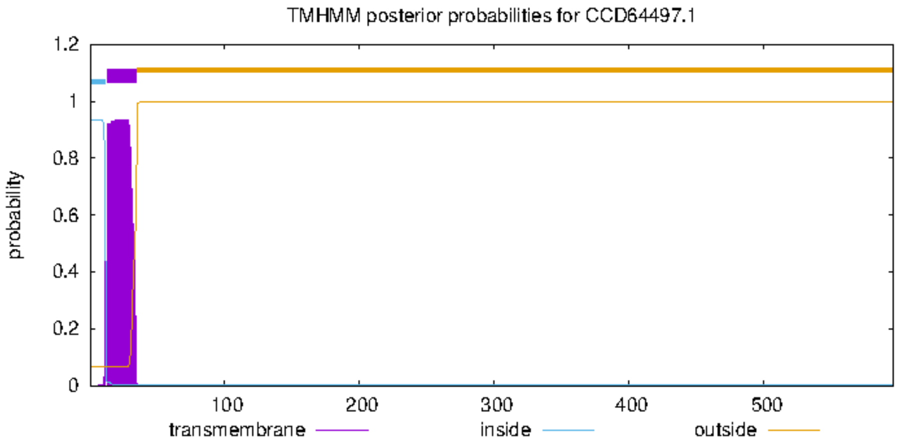

unnamed protein product [Anisakis simplex]

GenBank: VDK57457.1

**unnamed protein product [Anisakis simplex]**

Sequence ID: VDK57457.1 Length: 542 Number of Matches: 1

Range 1: 12 to 535 GenPept Graphics ▼ Next Match ▲ Pr

| Score | Expect | Method | Identities | Positives | Gaps |
|---|---|---|---|---|---|
| 385 bits(988) | 1e-127 | Compositional matrix adjust. | 207/529(39%) | 305/529(57%) | 45/529(8%) |

```
542 aa
>VDK57457.1 unnamed protein product [Anisakis simplex]
MQYFYANNCFQNSFYTNRIVHFDLKGAAPKVDYLKQVFRLIKENGATGILIEWEDMFPFNGILEENRCSD
AYTVTEVRDLLSTAQTLGLDVIPLVQTFGHLEWILKLDKFRKYRQSDQYAQVICMKDNDGVELVKEAIKQ
VVQMHKPYGIKYFHIGADEAFEKLFTFQYGTCAKDVAFLSQAGESGLENLAVGHIADIAKYVKSLLEPTA
KLVCNFKFSXLFATVNQYKLNDIIEPVVWDYSETLQQQNEYTWRELSAAFSNVWGASAFKGANNPSEQLI
DINHYLLNNVAWLDQKKEFGPLFQNFRGIILTGWQRYDHFAVLCELLPVSIPSLVVNLQVAVKGLDAHEK
SIARAVAKQLKCPYSVSLRHPESFGKCIFPGSSVFEHIQIEWPKMEETIKKEVSDNHQIRGWLNRFNIRH
NYTQLWYLKTLSDVINENYEQLVEMEHMIRHVVSFIKIRGIMALECKRAISSTVIFKKFSSFRHSMQQIY
KNDTIDEWIYQYFDPTMEKMKGYKSAIKRLQELRVFPKRTFAIRRDLESRTP
```

```
# VDK57457.1 Length: 542
# VDK57457.1 Number of predicted TMHs:  0
# VDK57457.1 Exp number of AAs in TMHs: 0.04211
# VDK57457.1 Exp number, first 60 AAs:  0.00039
# VDK57457.1 Total prob of N-in:        0.00390
VDK57457.1      TMHMM2.0        outside      1    542
```

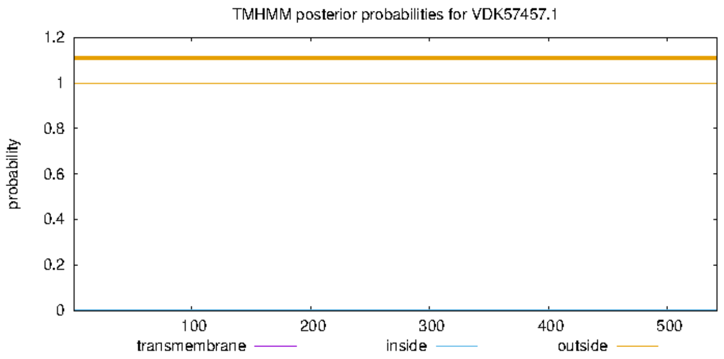


Multalin

```
           1        10        20        30        40        50        60        70        80        90       100       110       120       130
CCD64497.1 MFPMRCIRRRSIDFVLKGVILTTICLFLFHMTSSYPKGGISQRALDSMQKEPQTPVLVQKRENFEESIDVAVEQESKKENPFVNQQASTENAKPITQEVKIERPSRDNEFYKNVVIHFDLKGAPPKVDYF
VDK57457.1                                                                                                     MQYFYANNCFQNSFYTNRIVHFDLKGAAPKVDYL
Consensus  ...................................................................................................q#ykiarncr#NeFYkNr!!HFDLKGAaPKVDYl

           131      140       150       160       170       180       190       200       210       220       230       240       250       260
CCD64497.1 LDLLRLIAKGGATGILLEWEDMFPWTGKLEQFKNTDAYSESDVDMILSEATKLKLDVIPLVQTFGHLEWILKYEEMRKYRENDAYPQVLCLGNEEGVEFVREMIRQVAKKHAKYGIPFFHIGADEAFE--
VDK57457.1 KQVFRLIKENGATGILIEWEDMFPFNGILEENRCSDAYTVTEVRDLLSTAQTLGLDVIPLVQTFGHLEWILKLDKFRKYRQSDQYAQVICMKDNDGVELVKEAIKQVVQMHKPYGIKYFHIGADEAFEKL
Consensus  l#llRLIaenGATGILiEWEDMFPfnGiLE#nrnsDAYses#VrdiLSeAqkLgLDVIPLVQTFGHLEWILKl#emRKYR#nDaYaQViC$g###GVElVrEaIrQVaqkHakYGIk%FHIGADEAFE..

           261      270       280       290       300       310       320       330       340       350       360       370       380       390
CCD64497.1 ----FGVCQESLDWIKKNGKNGRKQLLALAHLKAIAEFAKQQTGDSTQILAWHDMLKDFDSRLIKNLELGQIIQPVVWDYSENIITLNDYIFSALAENFPTMWASSAYKGANYPSASTSEVRHYETNNRN
VDK57457.1 FTFQYGTCAKDVAFLSQAGESGLENL-AVGHIADIAKYVKSLLEPTAKLVCNFKFSXLFAT--VNQYKLNDIIEPVVWDYSETLQQQNEYTWRELSAAFSNVWGASAFKGANNPSEQLIDINHYLLNNVA
Consensus  ....%GtCaedlafikqaGenGre#L.AlaHiaaIAe%aKqqledsaqilanhdmlkdFas..!n#leLn#II#PVVWDYSEniqqqN#YifraLaaaFpnmWaaSA%KGANnPSaqli#!rHYelNNra

           391      400       410       420       430       440       450       460       470       480       490       500       510       520
CCD64497.1 WIRTKQNQERKFKNGFQGIIVTGWQRYDHLAGLCETLPIGTASMMLQMQIALNAPALDLEGTRQKAATLLECQGFNVDGVKVVSNQCKYRGFQTYLIYQSEVPNLFARIDSELSKNHHLMGWANRYNRKY
VDK57457.1 WLDQKKEFGPLFQN-FRGIILTGWQRYDHFAVLCELLPVSIPSLVVNLQVAVKGLDAHEKSIARAVAKQLKCPYSVSLRHPESFGKCIFPGSSVFEHIQIEWPKMEETIKKEVSDNHQIRGWLNRFNIRH
Consensus  WirqKq#qerlFqN.FrGIIlTGWQRYDHlAgLCElLP!giaS$ml#$Q!Alnalaadeegiaraa Akq LeCqgfnsdrhkesfnqCi%rGfqt%ehiQiEwPn$earIdkElSdNHqirGWaNR%Nrrh

           521      530       540       550       560       570       580       590       600       610       620       630       64042
CCD64497.1 NISQNWYHREMLPFVQQLVGQYDRVE--------------------------------SDLRASMKDLYFENTIDEFIYENLGEMSEKLHGYLEEIQRLDKLRAWPKRHFPIKK
VDK57457.1 NYTQLWYLKTLSDVINENYEQLVEMEHMIRHVVSFIKIRGIMALECKRAISSTVIFKKFSSFRHSMQQIYKNDTIDEWIYQYFDPTMEKMKGYKSAIKRLQELRVFPKRTFAIRRDLESRTP
Consensus  NisQnWYhre$ldf!##nyeQldrmE................................SdlRaSMq#iYk##TIDEfIY#nldemmEK$hGYleaIqRL#eLRafPKRhFaIrr.......
```

Uniprot: glycosyl hydrolase 20 family

#### HEX-3

Beta-N-acetylhexosaminidase [Caenorhabditis elegans]

GenBank: NP_499390.3, 591 aa

```
>NP_499390.3 beta-N-acetylhexosaminidase [Caenorhabditis elegans]
MLRGFFGRRSRSAWVRLIYLCIVTIIFLFATSQFKSTTTHARFVPEPSDPRLHPNSPLQAPKAQEPAAAE
TTKKPSRQPHHTTAKPRTENVEINGEKYPKLSPDGNLIPQRRIVHLDLKGAPYKPEFFTELFAFFNRIQA
TGILLEWEDMFPFKGRLRGAINKNAYSMETVEHILQEAQKHHLQIIPLVQTMGHLEWILKLEEFAHLRED
TRFPQVICFSDENAWELIKEMIEEVANVHKKYGMSYFHIGADEAFQIGICNASITQIKKEFTRERLMLWH
IARTARFVKEKYPETQVLAWHDMLASAMESDIEDYKLTELLQPVLWNYAEDLDIYLPRSTWMILRNFRNV
WGSSAWKGADGPARYSTNANHYLKNHESWIKQFTMVYKDFEVVEGLIMAGWSRYDHFAVLAETIPVALPT
LAMSMETMIEGRPLAGNYPVTSELLQCTPPLDLGFTATGCKFPGNRIYELINEMYQKQMQLRTYRLDDYE
LNGWLSRVADDYSVSSHWYIDKIENMIEMHATPLEQLADDLRFEMERIFFKDTVDEFIFTYLGEDLEWFN
RKRETIRKVSTSQTFPKRPFIESAKSEKTCT
```

```
# WEBSEQUENCE Length: 591
# WEBSEQUENCE Number of predicted TMHs:  1
# WEBSEQUENCE Exp number of AAs in TMHs: 21.2467299999999
# WEBSEQUENCE Exp number, first 60 AAs:  21.2366
# WEBSEQUENCE Total prob of N-in:        0.99905
# WEBSEQUENCE POSSIBLE N-term signal sequence
WEBSEQUENCE     TMHMM2.0        inside       1    11
WEBSEQUENCE     TMHMM2.0        TMhelix     12    34
WEBSEQUENCE     TMHMM2.0        outside     35   591
```

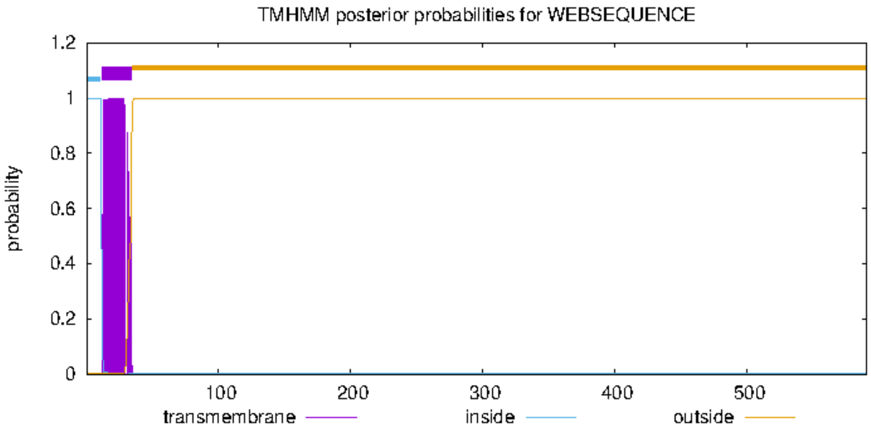

unnamed protein product [Anisakis simplex]

GenBank: VDK57457.1

#### unnamed protein product [Anisakis simplex]

**Sequence ID:** VDK57457.1 **Length:** 542 **Number of Matches:** 1

**Range 1: 18 to 531** GenPept Graphics ▼ Next Match ▲ Prev

| Score | Expect | Method | Identities | Positives | Gaps |
|---|---|---|---|---|---|
| 301 bits(771) | 2e-95 | Compositional matrix adjust. | 177/519(34%) | 277/519(53%) | 55/519(10%) |

```
>VDK57457.1 unnamed protein product [Anisakis simplex]
MQYFYANNCFQNSFYTNRIVHFDLKGAAPKVDYLKQVFRLIKENGATGILIEWEDMFPFNGILEENRCSD
AYTVTEVRDLLSTAQTLGLDVIPLVQTFGHLEWILKLDKFRKYRQSDQYAQVICMKDNDGVELVKEAIKQ
VVQMHKPYGIKYFHIGADEAFEKLFTFQYGTCAKDVAFLSQAGESGLENLAVGHIADIAKYVKSLLEPTA
KLVCNFKFSXLFATVNQYKLNDIIEPVVWDYSETLQQQNEYTWRELSAAFSNVWGASAFKGANNPSEQLI
DINHYLLNNVAWLDQKKEFGPLFQNFRGIILTGWQRYDHFAVLCELLPVSIPSLVVNLQVAVKGLDAHEK
SIARAVAKQLKCPYSVSLRHPESFGKCIFPGSSVFEHIQIEWPKMEETIKKEVSDNHQIRGWLNRFNIRH
NYTQLWYLKTLSDVINENYEQLVEMEHMIRHVVSFIKIRGIMALECKRAISSTVIFKKFSSFRHSMQQIY
KNDTIDEWIYQYFDPTMEKMKGYKSAIKRLQELRVFPKRTFAIRRDLESRTP
```

```
# VDK57457.1 Length: 542
# VDK57457.1 Number of predicted TMHs:  0
# VDK57457.1 Exp number of AAs in TMHs: 0.04211
# VDK57457.1 Exp number, first 60 AAs:  0.00039
# VDK57457.1 Total prob of N-in:        0.00390
VDK57457.1      TMHMM2.0        outside      1   542
```

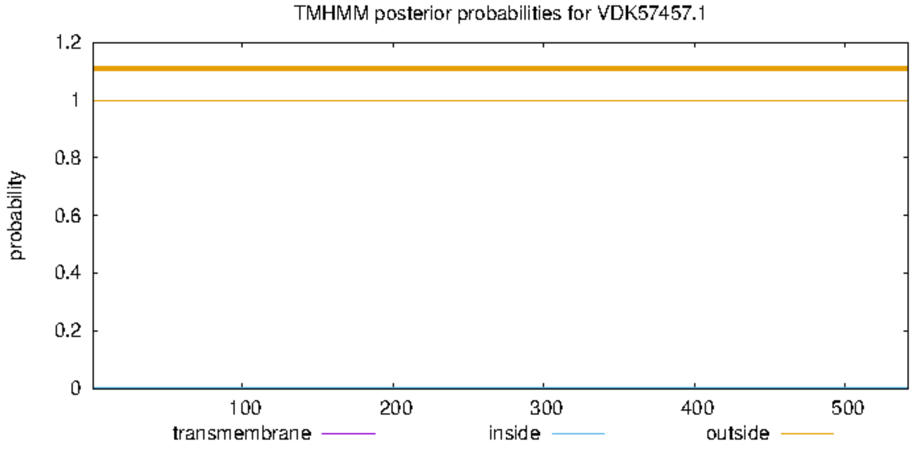


```
            1       10        20        30        40        50        60        70        80        90        100       110       120       130
            |--------+---------+---------+---------+---------+---------+---------+---------+---------+---------+---------+---------+---------|
CAA19506.3  MLRGFFGRRSRSAWVRLIYLCIVTIIFLFATSQFKSTTTHARFVPEPSDPRLHPNSPLQAPKAQEPAAAETTKKPSRQPHHTTAKPRTENVEINGEKYPKLSPDGNLIPQRRIVHLDLKGAPYKPEFFTE
VDK57457.1                                                                                                MQYFYANNCFQNSFYTNRIVHFDLKGAAPKVDYLKQ
Consensus   ..............................................................................................g#k%panncdqNlipqrRIVHlDLKGAapKp#%lk#

            131    140       150       160       170       180       190       200       210       220       230       240       250       260
            |--------+---------+---------+---------+---------+---------+---------+---------+---------+---------+---------+---------+---------|
CAA19506.3  LFAFFNRIQATGILLEWEDMFPFKGRLRGAINKNAYSMETVEHILQEAQKHHLQIIPLVQTMGHLEWILKLEEFAHLREDTRFPQVICFSDENAWELIKEMIEEVANVHKKYGMSYFHIGADEAF-----
VDK57457.1  VFRLIKENGATGILIEWEDMFPFNGILEENRCSDAYTVTEVRDLLSTAQTLGLDVIPLVQTFGHLEWILKLDKFRKYRQSDQYAQVICMKDNDGVELVKEAIKQVVQMHKPYGIKYFHIGADEAFEKLFT
Consensus   lFalinrnqATGILiEWEDMFPFnGrLrearnk#AYsmeeVrdiLqeAQkhgL#!IPLVQTmGHLEWILKL#eFahlR#ddr%aQVICmkD##awEL!KEaIe#Va#mHKkYGikYFHIGADEAF.....

            261    270       280       290       300       310       320       330       340       350       360       370       380       390
            |--------+---------+---------+---------+---------+---------+---------+---------+---------+---------+---------+---------+---------|
CAA19506.3  -QIGICNASITQIKK--EFTRERLMLWHIARTARFVKEKYPETQVLAWHDMLASAMESDIEDYKLTELLQPVLWNYAEDLDIYLPRSTWMILRNFRNVWGSSAWKGADGPARYSTNANHYLKNHESWIKQ
VDK57457.1  FQYGTCAKDVAFLSQAGESGLENLAVGHIADIAKYVKSLLEPTAKLVCNFKF-SXLFATVNQYKLNDIIEPVVWDYSETLQQQNEYTWRELSAAFSNVWGASAFKGANNPSEQLIDINHYLLNNVAWLDQ
Consensus   .QiGiCaad!aqikq..EfgrErLalgHIAriAr%VKelleeTakLacndkl.Sa$ead!##YKLn#ii#PVlW#YaEdL#qqnerstreilaaFrNVWGaSAfKGA#nParqli#aNHYLlNneaWidQ

            391    400       410       420       430       440       450       460       470       480       490       500       510       520
            |--------+---------+---------+---------+---------+---------+---------+---------+---------+---------+---------+---------+---------|
CAA19506.3  FTMVYKDFEVVEGLIMAGWSRYDHFAVLAETIPVALPTLAMSMETMIEGRPLAGNYPVTSELLQCTPPLDLGF----TATGCKFPGNRIYELINEMYQKQMQ-LRTYRLDDYELNGWLSRVADDYSVSSH
VDK57457.1  KKEFGPLFQNFRGIILTGWQRYDHFAVLCELLPVSIPSLVVNLQVAVKGLDAHEKSIARAVAKQLKCPYSVSLRHPESFGKCIFPGSSVFEHIQIEWPKMEETIKKEVSDNHQIRGWLNRFNIRHNYTQL
Consensus   kkefgkdF#nfrGiI$aGWqRYDHFAVLaEliPVaiPsLamn$#ta!eGrdaaensiaraealQckcPldlgl....saggCiFPGnr!%EhI#eewqKqe#.irkerlD#h#irGWLnRfadrhnysqh

            521    530       540       550       560       570       580       590       600       610       620       630    637
            |--------+---------+---------+---------+---------+---------+---------+---------+---------+---------+---------+------|
CAA19506.3  WYI--------DKIENMIEMH-------------------------ATPLEQLADDLRFEMERIFFKDTVDEFIFTYLGEDLEWFNRKRETIRKVSTSQTFPKRPFIESAKSEKTCT
VDK57457.1  WYLKTLSDVINENYEQLVEMEHMIRHVVSFIKIRGIMALECKRAISSTVIFKKFSSFRHSMQQIYKNDTIDEWIYQYFDPTMEKMKGYKSAIKRLQELRVFPKRTFAIRRDLESRTP
Consensus   WYi........#niE#$!EMe.........................aTpieqladdlRheM#rI%knDT!DEfI%qYlded$EkmnrkreaIrrlqelrtFPKRpFaeradlEkrcp
```

#### HEX-4

NP_740792.3, 491 aa

>HEX-4
MHKMSKLCFLALLSVTFMLIFVLTTPYSNDRSSYAAYEGGIDPRKTRQFKNIIVHLDLKGAPPRVEYLIEFFKLLSKHHV
DGILIEYEDMFPYSGDIEEIRRDLHYSENDIRRIIQAAEVHNLEVIPLIQSFGHLEFVLKKSKFMGLSEDLIDLNTICIS
DSKSIDIVKQMIEQIRRLHPNSTRIHIGADEAYHVAEDQRCIERMEKESIGKSDLKLEHIAKIGKFARENAGFETVFAWN
DMFDKESEETIRKSKINKFIVPVVWGYRTDVTENGYFPDGLFERIFNVFDRFYVASAFKGADGARQQFSNISRYLENQKS
YVNLMDLHKNAAAQKVDGIFVTGWSRFNHFNALCELLPVAIPSLIVDLFYLNYQLTEKDAWRAMKSSLECENRRHLRGIL
AESTVHGCKFPGADVFEIIMHDWKRAVDRRIFGKPDQQPSPDSEAIEILKNLKKSLNQSILYKTDADEVFNQYLHDYHSL
IAQNTRTEITN

>NP_740792.3 beta-N-acetylhexosaminidase [Caenorhabditis elegans]
MHKMSKLCFLALLSVTFMLIFVLTTPYSNDRSSYAAYEGGIDPRKTRQFKNIIVHLDLKGAPPRVEYLIE
FFKLLSKHHVDGILIEYEDMFPYSGDIEEIRRDLHYSENDIRRIIQAAEVHNLEVIPLIQSFGHLEFVLK
KSKFMGLSEDLIDLNTICISDSKSIDIVKQMIEQIRRLHPNSTRIHIGADEAYHVAEDQRCIERMEKESI
GKSDLKLEHIAKIGKFARENAGFETVFAWNDMFDKESEETIRKSKINKFIVPVVWGYRTDVTENGYFPDG
LFERIFNVFDRFYVASAFKGADGARQQFSNISRYLENQKSYVNLMDLHKNAAAQKVDGIFVTGWSRFNHF
NALCELLPVAIPSLIVDLFYLNYQLTEKDAWRAMKSSLECENRRHLRGILAESTVHGCKFPGADVFEIIM
HDWKRAVDRRIFGKPDQQPSPDSEAIEILKNLKKSLNQSILYKTDADEVFNQYLHDYHSLIAQNTRTEIT
N

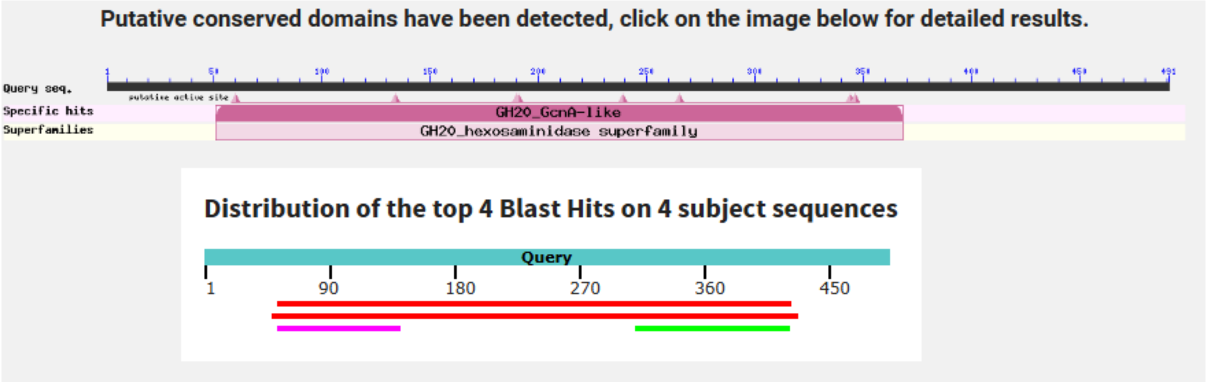

>VDK49770.1 unnamed protein product [Anisakis simplex]

MQKIVHFDLKGAPPKLEYYAELFPMLKRLNVDGVLMEYEDMFPYSGDIAILKRSIAYTRDQIASILSMAK

QNDLEVIPLVQTFGHMEFALKYNRFAYLREDQEQIDTICPSESGSWILITEMLRQVRALHPESKRIHIGS

DEAWQIAKDQRCLDRLRSELGSSTERLKLSHISRTANFAKHELGFDEVLAWNDMFGETNEQLLLQYKLGE

LLTPVVWGYSVNVTQPNYFPGGMFERYSKVFDRLMFASAFKGANGKDQKFANADRYLANQLSYVDLYRIN

ETHLIGRLKGIILTGWQRYSHTLPLCEILPVGIPSLVMDIVYLNNTTVAVDELRRLTREFLDCRSPEPEF

APIMLGTESYYPPSDIAYTNCTFPGHQIYDIVIISYYLTNIYVH

```
# VDK49770.1 Length: 394
# VDK49770.1 Number of predicted TMHs:  0
# VDK49770.1 Exp number of AAs in TMHs: 0.27093
# VDK49770.1 Exp number, first 60 AAs:  0.0001
# VDK49770.1 Total prob of N-in:        0.01120
VDK49770.1      TMHMM2.0        outside      1   394
```

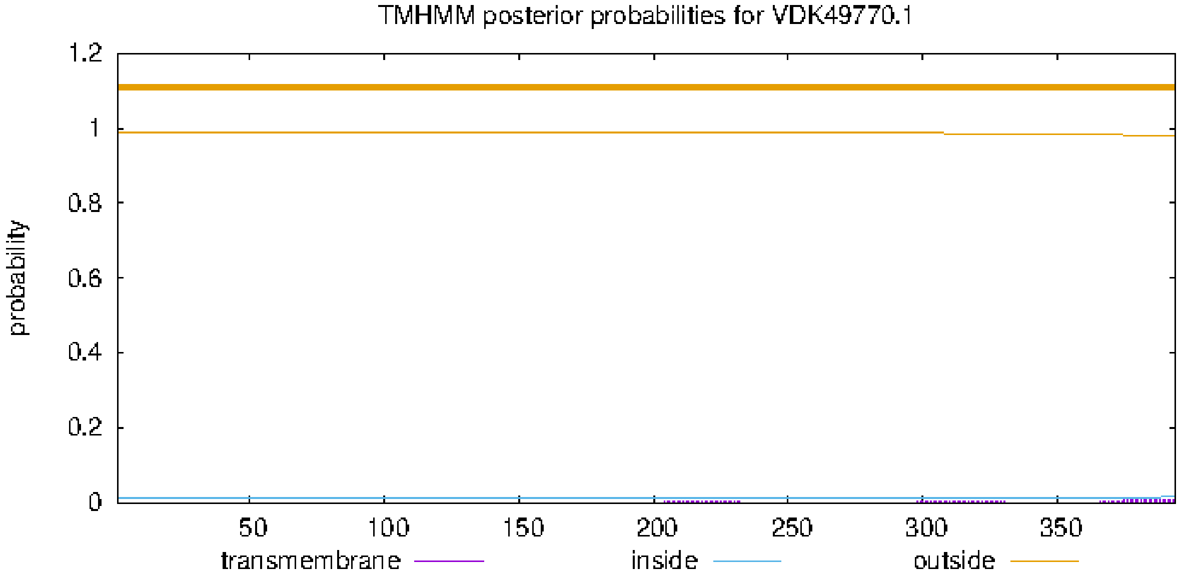

### HEX-5

>HEX-5
MLLRRTICILACIVQFATCGYQRSIVHFDMKGAPPKVAYFKQLLTTISGLGATGVLLEWEDMFPYQGGLSRVVNKNAYTE
EEVISVLEHAQQLQLEVIPLVQTLAHMEWILKTEEYSVLREDERYPMVACIGNPESLDIILDSVNQLMRIHSNFNTGYVH
IGADEAFQVGICDADREILPVKYDNNKLRMIFDHLRMVSLNITEEYPSTKVLMWYDELKSAPLELIKEYNLDNLVIPVVW
KYTANLDNDLPSEMWKNMSYSFKEVWGGSAFKGADGASRYWNRLKPYILNNKEWYLQNEKYKPQFTTFDSIIITGWQRYD
HFASLCELWPTSMVSLALNLIVLTKFHIDKESAEQVIQALNCPQTTTLDQLVAGSDRCRFPGYRVRDSIRDYVQLKTFFE
NSTWVHNRENGWLQSSHMRISASNPYYIDAIGKAYERTLKKLDTVSNSLSTSFSEVFYPDVIEEFKTDYFQPFYEDLQKR
KESVDNIDTKRFYVPRPWFRR

NP_001366664.1, 501 aa

>NP_001366664.1 beta-N-acetylhexosaminidase [Caenorhabditis elegans]
MLLRRTICILACIVQFATCGYQRSIVHFDMKGAPPKVAYFKQLLTTISGLGATGVLLEWEDMFPYQGGLS
RVVNKNAYTEEEVISVLEHAQQLQLEVIPLVQTLAHMEWILKTEEYSVLREDERYPMVACIGNPESLDII
LDSVNQLMRIHSNFNTGYVHIGADEAFQVGICDADREILPVKYDNNKLRMIFDHLRMVSLNITEEYPSTK
VLMWYDELKSAPLELIKEYNLDNLVIPVVWKYTANLDNDLPSEMWKNMSYSFKEVWGGSAFKGADGASRY
WNRLKPYILNNKEWYLQNEKYKPQFTTFDSIIITGWQRYDHFASLCELWPTSMVSLALNLIVLTKFHIDK
ESAEQVIQALNCPQTTTLDQLVAGSDRCRFPGYRVRDSIRDYVQLKTFFENSTWVHNRENGWLQSSHMRI
SASNPYYIDAIGKAYERTLKKLDTVSNSLSTSFSEVFYPDVIEEFKTDYFQPFYEDLQKRKESVDNIDTK
RFYVPRPWFRR

select all *4 sequences selected* GenPept Graphics Distance tree of results Multiple alignment MSA Viewer

| Description | Scientific Name | Max Score | Total Score | Query Cover | E value | Per. Ident | Acc. Len | Accession |
|---|---|---|---|---|---|---|---|---|
| unnamed protein product [Anisakis simplex] | Anisakis simplex | 276 | 276 | 97% | 6e-87 | 31.76% | 542 | VDK57457.1 |
| unnamed protein product [Anisakis simplex] | Anisakis simplex | 201 | 201 | 77% | 6e-60 | 31.09% | 394 | VDK49770.1 |
| unnamed protein product, partial [Anisakis simplex] | Anisakis simplex | 109 | 109 | 18% | 4e-28 | 53.33% | 186 | VDK64756.1 |
| unnamed protein product [Anisakis simplex] | Anisakis simplex | 75.9 | 75.9 | 34% | 3e-16 | 29.78% | 186 | VDK21806.1 |

**Putative conserved domains have been detected, click on the image below for detailed results.**

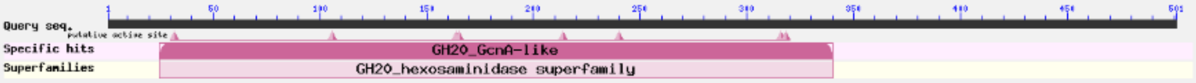


**Distribution of the top 4 Blast Hits on 4 subject sequences**

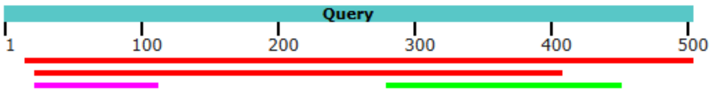

542 aa

>VDK57457.1 unnamed protein product [Anisakis simplex]

MQYFYANNCFQNSFYTNRIVHFDLKGAAPKVDYLKQVFRLIKENGATGILIEWEDMFPFNGILEENRCSD
AYTVTEVRDLLSTAQTLGLDVIPLVQTFGHLEWILKLDKFRKYRQSDQYAQVICMKDNDGVELVKEAIKQ
VVQMHKPYGIKYFHIGADEAFEKLFTFQYGTCAKDVAFLSQAGESGLENLAVGHIADIAKYVKSLLEPTA
KLVCNFKFSXLFATVNQYKLNDIIEPVVWDYSETLQQQNEYTWRELSAAFSNVWGASAFKGANNPSEQLI
DINHYLLNNVAWLDQKKEFGPLFQNFRGIILTGWQRYDHFAVLCELLPVSIPSLVVNLQVAVKGLDAHEK
SIARAVAKQLKCPYSVSLRHPESFGKCIFPGSSVFEHIQIEWPKMEETIKKEVSDNHQIRGWLNRFNIRH
NYTQLWYLKTLSDVINENYEQLVEMEHMIRHVVSFIKIRGIMALECKRAISSTVIFKKFSSFRHSMQQIY
KNDTIDEWIYQYFDPTMEKMKGYKSAIKRLQELRVFPKRTFAIRRDLESRTP

```
# WEBSEQUENCE Length: 542
# WEBSEQUENCE Number of predicted TMHs:  0
# WEBSEQUENCE Exp number of AAs in TMHs: 0.04211
# WEBSEQUENCE Exp number, first 60 AAs:  0.00039
# WEBSEQUENCE Total prob of N-in:        0.00390
WEBSEQUENCE	TMHMM2.0	outside	     1   542
```

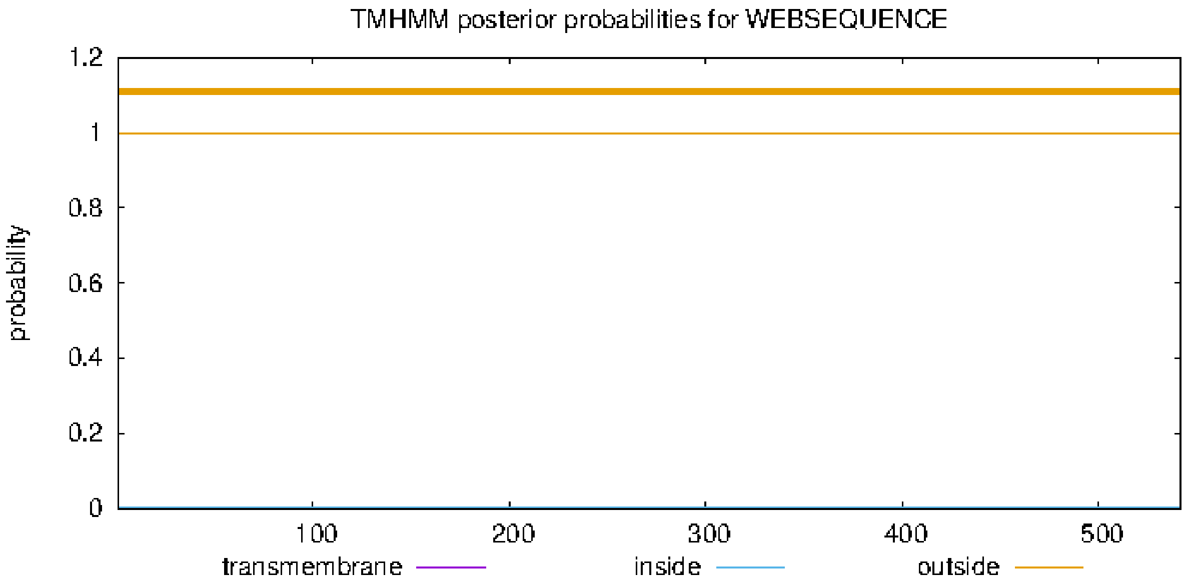

### AMAN-2

NCBI Reference Sequence: NP_505995.2

```
Wormbase
>AMAN-2
MGKRNFYIILCLGVFLTVSLYLYNGIETGAEALTKRQRYVDDLRRKIGNLEHVAEENGRTIDRLEQEVQRAKAEKSVDFD
EEKEKTEEKEVEKEEKEVAPVPVRGNRGEMAHIHQVKQHIKPTPSMKDVCGIRENVSIAHSDLQMLDLYDTWKFENPDGG
VWKQGWKIEYDAEKVKSLPRLEVIVIPHSHCDPGWIMTFEEYYNRQTRNILDGMAKHLAEKDEMRFIYAEISFFETWWRD
QADEIKKKVKGYLEAGKFEIVTGGWVMTDEANAHYHSMITELFEGHEWIQNHLGKSAIPQSHWSIDPFGLSPSMPHLLTS
ANITNAVIQRVHYSVKRELALKKNLEFYWRQLFGSTGHPDLRSHIMPFYSYDIPHTCGPEPSVCCQFDFRRMPEGGKSCD
WGIPPQKINDDNVAHRAEMIYDQYRKKSQLFKNNVIFQPLGDDFRYDIDFEWNSQYENYKKLFEYMNSKSEWNVHAQFGT
LSDYFKKLDTAISASGEQLPTFSGDFFTYADRDDHYWSGYFTSRPFYKQLDRVLQHYLRSAEIAFTLANIEEEGMVEAKI
FEKLVTARRALSLFQHHDGVTGTAKDHVVLDYGQKMIDALNACEDILSEALVVLLGIDSTNKMQMDEHRVNENLLPEKRV
YKIGQNVVLFNTLSRNRNEPICIQVDSLDAGVEADPPIKKQQVSPVIAYDEEKKTLVVKNGIFELCFMLSLGPMESVSFR
LVKNTTTSKVEIITNNAAEFKETSFKSSSTSGDFTVKNDKVEAEFDGENGMIKRATSLVDDKPIDLNSHFIHYGARKSKR
KFANGNEDNPAGAYLFLPDGEARELKKQSSDWILVKGEVVQKVFATPNNDLKILQTYTLYQGLPWIDLDNEVDVRSKENF
ELALRFSSSVNSGDEFFTDLNGMQMIKRRRQTKLPTQANFYPMSAGVYIEDDTTRMSIHSAQALGVSSLSSGQIEIMLDR
RLSSDDNRGLQQGVRDNKRTVAHFRIVIEPMSSSSGNKKEERVGFHSHVGHLATWSLHYPLVKMIGDATPKSISSKNVEQ
ELNCDLHLVTFRTLASPTTYEANERSTAAEKKAAMVMHRVVPDCRSRLTLPDTSCLATGLEIEPLKLISTLKSAKKTSLT
NLYEGNKAEQFRLQPNDISSILVSF

>NP_505995.2 Alpha-mannosidase [Caenorhabditis elegans]
MGKRNFYIILCLGVFLTVSLYLYNGIETGAEALTKRQRYVDDLRRKIGNLEHVAEENGRTIDRLEQEVQR
AKAEKSVDFDEEKEKTEEKEVEKEEKEVAPVPVRGNRGEMAHIHQVKQHIKPTPSMKDVCGIRENVSIAH
SDLQMLDLYDTWKFENPDGGVWKQGWKIEYDAEKVKSLPRLEVIVIPHSHCDPGWIMTFEEYYNRQTRNI
LDGMAKHLAEKDEMRFIYAEISFFETWWRDQADEIKKKVKGYLEAGKFEIVTGGWVMTDEANAHYHSMIT
ELFEGHEWIQNHLGKSAIPQSHWSIDPFGLSPSMPHLLTSANITNAVIQRVHYSVKRELALKKNLEFYWR
QLFGSTGHPDLRSHIMPFYSYDIPHTCGPEPSVCCQFDFRRMPEGGKSCDWGIPPQKINDDNVAHRAEMI
YDQYRKKSQLFKNNVIFQPLGDDFRYDIDFEWNSQYENYKKLFEYMNSKSEWNVHAQFGTLSDYFKKLDT
AISASGEQLPTFSGDFFTYADRDDHYWSGYFTSRPFYKQLDRVLQHYLRSAEIAFTLANIEEEGMVEAKI
FEKLVTARRALSLFQHHDGVTGTAKDHVVLDYGQKMIDALNACEDILSEALVVLLGIDSTNKMQMDEHRV
NENLLPEKRVYKIGQNVVLFNTLSRNRNEPICIQVDSLDAGVEADPPIKKQQVSPVIAYDEEKKTLVVKN
GIFELCFMLSLGPMESVSFRLVKNTTTSKVEIITNNAAEFKETSFKSSSTSGDFTVKNDKVEAEFDGENG
MIKRATSLVDDKPIDLNSHFIHYGARKSKRKFANGNEDNPAGAYLFLPDGEARELKKQSSDWILVKGEVV
QKVFATPNNDLKILQTYTLYQGLPWIDLDNEVDVRSKENFELALRFSSSVNSGDEFFTDLNGMQMIKRRR
QTKLPTQANFYPMSAGVYIEDDTTRMSIHSAQALGVSSLSSGQIEIMLDRRLSSDDNRGLQQGVRDNKRT
VAHFRIVIEPMSSSSGNKKEERVGFHSHVGHLATWSLHYPLVKMIGDATPKSISSKNVEQELNCDLHLVT
FRTLASPTTYEANERSTAAEKKAAMVMHRVVPDCRSRLTLPDTSCLATGLEIEPLKLISTLKSAKKTSLT
NLYEGNKAEQFRLQPNDISSILVSF
```

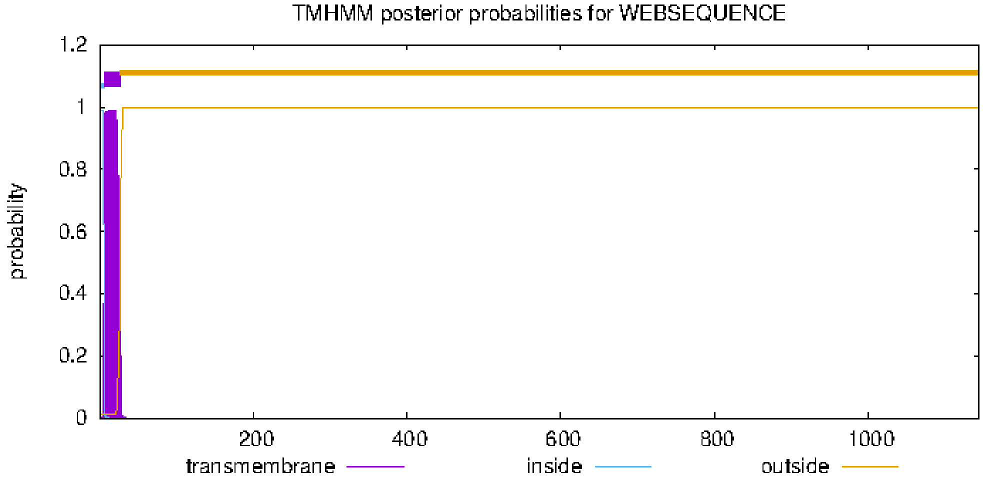

#### unnamed protein product [Anisakis simplex]

**Sequence ID:** VDK44985.1 **Length:** 1122 **Number of Matches:** 1

**Range 1: 186 to 1117** GenPept Graphics ▼ Next Match ▲ Previous Match

| Score | Expect | Method | Identities | Positives | Gaps |
|---|---|---|---|---|---|
| 809 bits(2090) | 0.0 | Compositional matrix adjust. | 458/1054(43%) | 620/1054(58%) | 157/1054(14%) |

#### AMAN-3

Wormbase

>AMAN-3, isoform a
MHRLALSAKRKVLLNPKTVSIYFFAILFTFLLAYHQRLGQHNNELHISRVVNMRSFVKEANNLSNSQKNPNIFEPKNEVC
QRPLTESSTNFNTFDLFESVVAKGNTLPPASKKSRTEKLKVYVLPFTHVDPGWLETFERYTKSTNQILDNMHQFMMKNEK
MRFMWAEFVFFERWWSLQNEQVKEDVKKLVTEGRLELATGSWVMTDEANPYFPVSVDNIVEGFQFIHKNFGIKPQTMWSN
DPFGYSNSVPYLFKKSGVHRTVINRIHHKLKQTLQSQKAIPFKWRQYFDATGEDDVLTQILPYTHYDILNSCGSDASVCC
EFDFKRMTHWSCPGPKPEKITNANVAAKAEKLVNQLEKMSEMYKAPVILMMHGDDFRFDMIEEWNQQHDNFIPVFDEINK
GSRVEIRFGTFTDYFNDLEKWYSNNKDSEPPTVSGDFFPYMCALGDYWTGYYTTRPFFKRQGRLLHSLIRNGDIMLSMLR
VNLKQRKIGENVKRLEAARRNLALFQHHDAITGTSKVSVMDNYSELLHASIVSTNIVLENLTKSDIDLYPRIHDGIELQT
IVDLESSEKEIRIFNSHLFEITDVFKIRVKEREVIVSIDGKIIEAQLEPFFQKSKVEKDSFLLLFQATVLPLSMMKVKVQ
RGDSGGLTKMAKIEAKDTNAWDLGSSWTIASSTSAPNLETPYFKVSNDPITGAIQSVNKLSDNSKFHCQQSFYNYKEAGG
GAYLMRLHTNPKEIIEYQWLKVSGPLRQSIYQKSTNVLQRLSIHNVEGPSGEEVDISMSIDITKERNTELMTRFSTKWDK
PLTYTDSVGMQLLRRDFYKLPVQSNYYPMPTAAVLQSGKQRLSIVSNVEHGARFLESGTVEINIDRILNQDDGKGLGTGP
DAIPIDMKPVDMKFKLIFDSLESDPTENSRYSTHSFRAQQAVQTIIYPPMMFNRPPKKEEDIEEIEEISNLKFPCDIQLL
TVRPLEDNKQLLILYRHATVCSSQKVGNCGGELKSSLMDLLIKLGAKQVQKTDLSGVTRVGGIIKVEYLPDYELKTFDFL
TLILYR

NP_001361919.1, 1046 aa

```
>NP_001361919.1 Alpha-mannosidase [Caenorhabditis elegans]
MHRLALSAKRKVLLNPKTVSIYFFAILFTFLLAYHQRLGQHNNELHISRVVNMRSFVKEANNLSNSQKNP
NIFEPKNEVCQRPLTESSTNFNTFDLFESVVAKGNTLPPASKKSRTEKLKVYVLPFTHVDPGWLETFERY
TKSTNQILDNMHQFMMKNEKMRFMWAEFVFFERWWSLQNEQVKEDVKKLVTEGRLELATGSWVMTDEANP
YFPVSVDNIVEGFQFIHKNFGIKPQTMWSNDPFGYSNSVPYLFKKSGVHRTVINRIHHKLKQTLQSQKAI
PFKWRQYFDATGEDDVLTQILPYTHYDILNSCGSDASVCCEFDFKRMTHWSCPGPKPEKITNANVAAKAE
KLVNQLEKMSEMYKAPVILMMHGDDFRFDMIEEWNQQHDNFIPVFDEINKGSRVEIRFGTFTDYFNDLEK
WYSNNKDSEPPTVSGDFFPYMCALGDYWTGYYTTRPFFKRQGRLLHSLIRNGDIMLSMLRVNLKQRKIGE
NVKRLEAARRNLALFQHHDAITGTSKVSVMDNYSELLHASIVSTNIVLENLTKSDIDLYPRIHDGIELQT
IVDLESSEKEIRIFNSHLFEITDVFKIRVKEREVIVSIDGKIIEAQLEPFFQKSKVEKDSFLLLFQATVL
PLSMMKVKVQRGDSGGLTKMAKIEAKDTNAWDLGSSWTIASSTSAPNLETPYFKVSNDPITGAIQSVNKL
SDNSKFHCQQSFYNYKEAGGGAYLMRLHTNPKEIIEYQWLKVSGPLRQSIYQKSTNVLQRLSIHNVEGPS
GEEVDISMSIDITKERNTELMTRFSTKWDKPLTYTDSVGMQLLRRDFYKLPVQSNYYPMPTAAVLQSGKQ
RLSIVSNVEHGARFLESGTVEINIDRILNQDDGKGLGTGPDAIPIDMKPVDMKFKLIFDSLESDPTENSR
YSTHSFRAQQAVQTIIYPPMMFNRPPKKEEDIEEIEEISNLKFPCDIQLLTVRPLEDNKQLLILYRHATV
CSSQKVGNCGGELKSSLMDLLIKLGAKQVQKTDLSGVTRVGGIIKVEYLPDYELKTFDFLTLILYR
```

```
# VWL57857.1 Length: 1045
# VWL57857.1 Number of predicted TMHs:  1
# VWL57857.1 Exp number of AAs in TMHs: 22.16184
# VWL57857.1 Exp number, first 60 AAs:  22.16148
# VWL57857.1 Total prob of N-in:        0.99720
# VWL57857.1 POSSIBLE N-term signal sequence
VWL57857.1      TMHMM2.0        inside       1    11
VWL57857.1      TMHMM2.0        TMhelix     12    34
VWL57857.1      TMHMM2.0        outside     35  1045
```

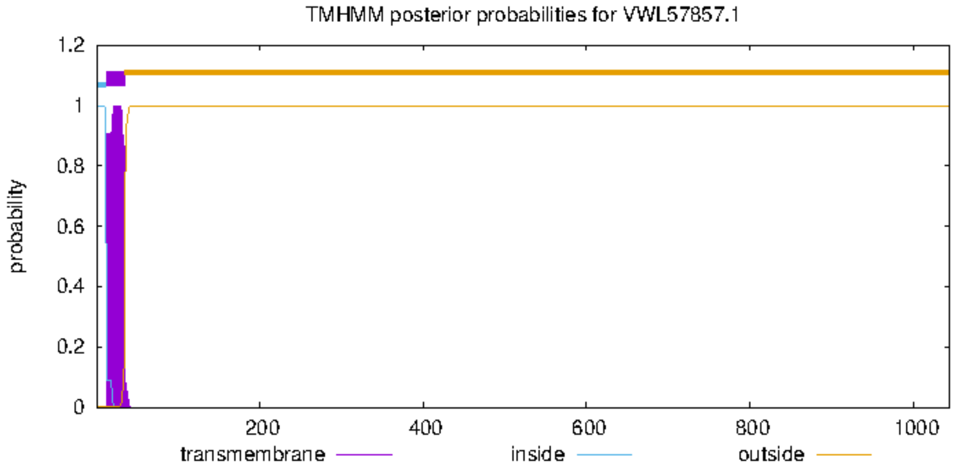


#### unnamed protein product [Anisakis simplex]

**Sequence ID:** VDK44985.1 **Length:** 1122 **Number of Matches:** 1

**Range 1: 233 to 1056** GenPept Graphics ▼ Next Match ▲ Previous M

| Score | Expect | Method | Identities | Positives | Gaps |
|---|---|---|---|---|---|
| 401 bits(1030) | 2e-122 | Compositional matrix adjust. | 285/928(31%) | 449/928(48%) | 160/928(17%) |

```
>VDK44985.1 unnamed protein product [Anisakis simplex]
MRVNTRKWLFGGGLCVFVFVELQKLELKLKTLEKEIKNNDAAMMEMRQKLRAERLKIKEFENMKKKISEA
SADGGENNVDSAGGGAPGVLQLPKKAEQIRKEQENVAAAREHAEEEEQAKRGVIAEPLNNNNDNNHIDNE
VPVKSDRVSNSRGVLLNRNGQTPTNKGAIVKKFFADLVMPYAEVGQDMCRAISNMSAARSDIQMLDLYNT
IPFDDPDGGVWKQGFDIKYDEEKVKQEKRLEIIVTPHSHTDPGWITTFEGYYNSQTKYIFENMLLSLQSM
ERMRFIYAEMSFFEKWWAEINDEQRAAVQRLLHQGRLEIVSGAWVMTDEANAHYFATVSEWIEGHEWIAN
HIPDYKPKNHWSIDPFGLSSTLAFVVSKANLSNALVQRVHYSVKKHLAEQKQLEFKWRQLWSGADASNDL
FTHVMPFYSYDVPHTCGPDPKVCCQFDFWRLSGQAACPWGVPPEEITERNLARRAAILYDQYRKKAQLFR
RNVLFVPLGDDFRYGSAEEWRLQHDNYIRLFDYINAKKEWNVHVRFGTLGDYFELDHARVEEFKGEADGE
VPVLSGDFFTYADRNDHYWSGYYTSRPFYKRMDRVLQHYLRSAEIIVSLAISKDSRAESIIQEMYGLLVE
ARRHMSLFQHHDGVTGTAKDDVVVDYGQNEVVVFNSLAQPRHEVVCVQTKHLKTEVSRPSYPNIPVPQQI
APVLKRTSRNIEFEFGKYELCFMATVPPFGFEVYKLHSTPNAVGGGASKVSLRSRKKITSSDFMTELIQE
PYFDLDNEFVEAHFDATTGLLKSVTPSDGHEVSVNLSFVQYGVRSKNPGKFQGGDDLSGAYLFLPNGPAR
EMRPQNGYQYVVVDGPIMKKTDIYDCCHGLLLLAISDCYCYFVIIFQMIKRKRFAKLPLQAHFYPMPTSA
FIEDNSNRMTIFSAQSLGVASLESGWLEVMLDRRLNQDDGRGLFQGVTDNKITRSKFRLLIEPLDVNKRL
NREPDSKSTTTAYHSLVGHYVSLQLQYPLMSMFSSAGQSETMASPTDYNEESNKKYAPAKSKALILHRLG
AECGSKLRLHSFCSTSDGKVKVSKLFEGKPTRIRETSLTLLYEGNSNDNFDEVQIKPMDMRTFRLDFTRG
FS
```

```
# VDK44985.1 Length: 1122
# VDK44985.1 Number of predicted TMHs:  0
# VDK44985.1 Exp number of AAs in TMHs: 7.90134999999999
# VDK44985.1 Exp number, first 60 AAs:  1.06627
# VDK44985.1 Total prob of N-in:        0.06930
VDK44985.1      TMHMM2.0        outside      1  1122
```

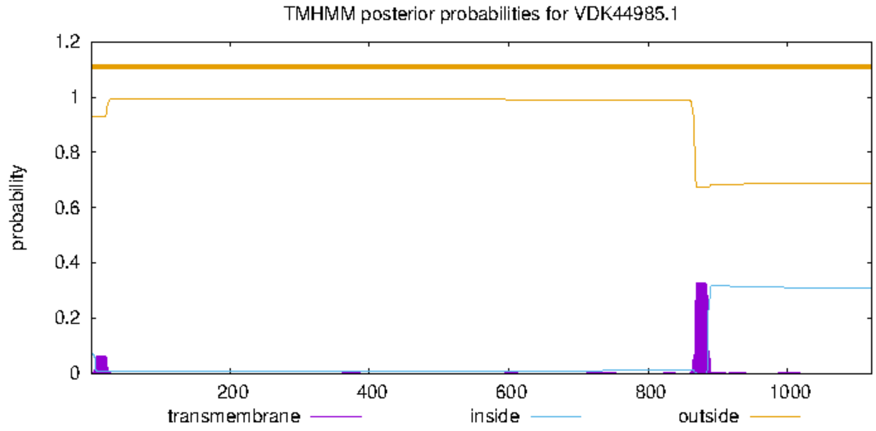


```
MSF:   1187    Check:   0        ..
Name: VWL57857.1      Len:  1187  Check: 4701  Weight:  1.00
Name: VDK44985.1      Len:  1187  Check:   75  Weight:  1.00
Name: Consensus       Len:  1187  Check: 9216  Weight:  0.00

//
```

```
              1       10        20        30        40        50        60        70        80        90        100       110       120       130
              |--------+---------+---------+---------+---------+---------+---------+---------+---------+---------+---------+---------+---------|
VWL57857.1                                                                                                                         MHRLALSAKRKVL
VDK44985.1    MRVNTRKWLFGGGLCVFVFVELQKLELKLKTLEKEIKNNDAAMMEMRQKLRAERLKIKEFENMKKKISEASADGGENNVDSAGGGAPGVLQLPKKAEQIRKEQENVAAAREHAEEEEQAKRGVIAEPLNN
 Consensus    .........................................................................................................................qarraliAerlnn

              131     140       150       160       170       180       190       200       210       220       230       240       250       260
              |--------+---------+---------+---------+---------+---------+---------+---------+---------+---------+---------+---------+---------|
VWL57857.1    LNPKTVSIYFFAILFTFLLAYHQRLGQHNNELHISRVVNMRSFVKEANNLSNSQKNPNIFEPKNEVCQRPLTESSTNFNTFDLFE---SVVAKGNTLPPASKKSRTEK-LKVYVLPFTHVDPGWLETFER
VDK44985.1    NNDNNHIDNEVPVKSDRVSNSRGVLLNRNGQTPTNKGAIVKKFFADLVMPYAEVGQDMCRAISNMSAARSDIQMLDLYNTIPFDDPDGGVWKQGFDIKYDEEKVKQEKRLEIIVTPHSHTDPGWITTFEG
 Consensus    nNdnnhidnefa!lfdrllasrqrLg#rNn#lhinrganmrkFfa#annlsaeqg#dncraikNesaaRpdi#mldn%NTidld#...gVwaqGndikpaeeKsrqEK.Le!iVlPhsHtDPGWieTFEr

              261     270       280       290       300       310       320       330       340       350       360       370       380       390
              |--------+---------+---------+---------+---------+---------+---------+---------+---------+---------+---------+---------+---------|
VWL57857.1    YTKS-TNQILDNMHQFMMKNEKMRFMWAEFVFFERWWSLQNEQVKEDVKKLVTEGRLELATGSWVMTDEANPYFPVSVDNIVEGFQFIHKNF-GIKPQTMWSNDPFGYSNSVPYLFKKSGVHRTVINRIH
VDK44985.1    YYNSQTKYIFENMLLSLQSMERMRFIYAEMSFFEKWWAEINDEQRAAVQRLLHQGRLEIVSGAWVMTDEANAHYFATVSEWIEGHEWIANHIPDYKPKNHWSIDPFGLSSTLAFVVSKANLSNALVQRVH
 Consensus    YtnS.TnqIl#NMhqf$qknErMRFiwAEmsFFErWWaeqN##qraaVqrLlh#GRLEiasGaWVMTDEANah%fasVd#i!EGh#fIanni.diKPqnhWSnDPFGlSnsla%lfkKanlhral!#R!H

              391     400       410       420       430       440       450       460       470       480       490       500       510       520
              |--------+---------+---------+---------+---------+---------+---------+---------+---------+---------+---------+---------+---------|
VWL57857.1    HKLKQTLQSQKAIPFKWRQYFD-ATGEDDVLTQILPYTHYDILNSCGSDASVCCEFDFKRMT-HWSCP-GPKPEKITNANVAAKAEKLVNQLEKMSEMYKAPVILMMHGDDFRFDMIEEWNQQHDNFIPV
VDK44985.1    YSVKKHLAEQKQLEFKWRQLWSGADASNDLFTHVMPFYSYDVPHTCGPDPKVCCQFDFWRLSGQAACPWGVPPEEITERNLARRAAILYDQYRKKAQLFRRNVLFVPLGDDFRYGSAEEWRLQHDNYIRL
 Consensus    hklKqhLaeQKaieFKWRQlfd.Adae#DllTq!$P%thYD!lnsCGpDakVCC#FDFkR$s.qaaCP.GpkPEeIT#aNlAarAaiLy#QlrKka#$%ranVilmmhGDDFR%dmaEEWrqQHDN%Irl

              521     530       540       550       560       570       580       590       600       610       620       630       640       650
              |--------+---------+---------+---------+---------+---------+---------+---------+---------+---------+---------+---------+---------|
VWL57857.1    FDEIN--KGSRVEIRFGTFTDYF---NDLEKWYSNNKDSEPPTVSGDFFPYMCALGDYWTGYYTTRPFFKRQGRLLHSLIRNGDIMLSML--RVNLKQRKIGENVKRLEAARRNLALFQHHDAITGTSKV
VDK44985.1    FDYINAKKEWNVHVRFGTLGDYFELDHARVEEFKGEADGEVPVLSGDFFTYADRNDHYWSGYYTSRPFYKRMDRVLQHYLRSAEIIVSLAISKDSRAESIIQEMYGLLVEARRHMSLFQHHDGVTGTAKD
 Consensus    FDeIN..KesrVe!RFGTlgDYF...nareee%kn#aDgEpPtlSGDFFpYadanddYWsGYYTsRPF%KRqdRlLqhliRna#IilS$a..rdnra#riIqEnygrLeaARRn$aLFQHHDa!TGTaKd

              651     660       670       680       690       700       710       720       730       740       750       760       770       780
              |--------+---------+---------+---------+---------+---------+---------+---------+---------+---------+---------+---------+---------|
VWL57857.1    SVMDNYSELLHASIVSTNIVLENLTKSDIDLYPRIHDGIELQTIVDLESSEKEIRIFNSHLFEITDVFKIRVKEREVIVSIDGKIIEAQLEPFFQKSKVEKDSFLLLFQATVLPLSMMKVKVQRGDS---
VDK44985.1    DVVVDYGQ-------NEVVVFNSLAQ------PR-HEVVCVQT--------KHLK---------TEVSRPSYPNIPVPQQIAPVLKRTSRNIEFEFGKYE-----LCFMATVPPFGFEVYKLHSTPNAVG
 Consensus    dVmd#Yg#.......nen!Vl#nLaq......PR.H#g!clQT........Keir.........T#Vfriryk#reViqqIagkiiraqr#ieF#kgKyE.....LcFqATVlPlgmekyKlqrgdn...

              781     790       800       810       820       830       840       850       860       870       880       890       900       910
              |--------+---------+---------+---------+---------+---------+---------+---------+---------+---------+---------+---------+---------|
VWL57857.1    GGLTKMAKIEAKDTNAWDLGSWTIASSTSAPNLETPYFKVSNDPITGAIQSVNKLSDNSKFHCQQSFYNYKEAGGGAYLMRLHTNPKEIIEYQWLKVSGPLRQSIYQKSTNVLQRLSIHNVEGPSGEEVD
VDK44985.1    GGASKVSLRSRKKITSSDFMTELIQEPYF--DLDNEFVEAHFDATTGLLKSVTP-SDGHEVSVNLSFVQY---GVRSKNPGKFQGGDDLSGAYLFLPNGPAREMRPQNGYQYVV------VDGP------
 Consensus    GGasKmalreaKdinasDlgselIaeptf..#L#ne%feahnDaiTGaiqSVnk.SDnhefhc#qSFy#Y...Ggraknmrlhqngd#iieaqlllpnGPaR#mrpQngt#ylq......V#GP......

              911     920       930       940       950       960       970       980       990       1000      1010      1020      1030      1040
              |--------+---------+---------+---------+---------+---------+---------+---------+---------+---------+---------+---------+---------|
VWL57857.1    ISMSIDITKERNTELMTRFSTKWDKPLTYTDSVGMQLLRRDFYKLPVQSNYYPMPTAAVLQSGKQRLSIVSNVEHGARFLESGTVEINIDRILNQDDGKGLGTGPDAIPIDMKPVDMKFKLIFDSLESDP
VDK44985.1    IMKKTDIYDCCHGLLLLAISDCY----CYFVIIFQMIKRKRFAKLPLQAHFYPMPTSAFIEDNSNRMTIFSAQSLGVASLESGWLEVMLDRRLNQDDGRGLFQGVT----DNKITRSKFRLLIEPLDVNK
 Consensus    ImkkiDItdcrngeL$laiSdcw....cYfdi!gqqilRrrFaKLPlQan%YPMPTaAfi#dnk#R$sIfSaqehGaafLESGtlE!niDRrLNQDDGrGLgqGpd....DnKitrmKFrLii#pL#s#k

              1041 1050       1060      1070      1080      1090      1100      1110      1120      1130      1140      1150      1160      1170
              |--------+---------+---------+---------+---------+---------+---------+---------+---------+---------+---------+---------+---------|
VWL57857.1    TENSRYSTHSFRAQQAVQTIIYPPMMFNRPPKKEEDIEEIEEISNLKFPCDIQLLTVRPLEDNK-QLLILYRHATVCSSQKVGNCGGELKSSLMDLLIKLGAKQVQKTDLSGVTRVGGIIKVEYLPDYEL
VDK44985.1    RLNREPDSKSTTTAYHSLVGHYVSLQLQYPLMSM--FSSAGQSETMASPTDYNEESNKKYAPAKSKALILHRLGAECGSKLRLHSFCSTSDGKVKVSKLFEGKPTRIRETSLTLLYEGNSNDNF-DEVQI
 Consensus    reNrrpdshSfraaqasqtghYpp$ql#rPlkke..ieeae#ien$afPcDi#eesnrkladaK.qaLILhRhaaeCgSqlrgncgcelkdglmdllilleaKqtrir#lSgtlryeGnind#%.d#y#i

              1171 1180   1187
              |--------+------|
VWL57857.1    KTFDFLTLILYR
VDK44985.1    KPMDMRTFRLDFTRGFS
 Consensus    KpmDmrTlrLdr.....
```
